# The Signaling Pathways Project: an integrated ‘omics knowledgebase for mammalian cellular signaling pathways

**DOI:** 10.1101/401729

**Authors:** Scott Ochsner, David Abraham, Kirt Martin, Wei Ding, Apollo McOwiti, Wasula Kankanamge, Zichen Wang, Kaitlyn Andreano, Ross A. Hamilton, Yue Chen, Angelica Hamilton, Marin L. Gantner, Michael Dehart, Shijing Qu, Susan G. Hilsenbeck, Lauren B. Becnel, Dave Bridges, Avi Ma’ayan, Janice M. Huss, Fabio Stossi, Charles E. Foulds, Anastasia Kralli, Donald P. McDonnell, Neil J. McKenna

## Abstract

Integrated mining of public transcriptomic and ChIP-Seq datasets has the potential to illuminate facets of mammalian cellular signaling pathways not yet explored in the research literature. Here, we designed a web knowledgebase, the Signaling Pathways Project (SPP), which incorporates stable community classifications of the four major categories of signaling pathway node (receptors, enzymes, transcription factors and co-nodes) and their cognate bioactive small molecules (BSMs). We then mapped over 10,000 public transcriptomic or cistromic experiments to their relevant signaling pathway node, BSM or biosample of study. To provide for prediction of pathway node-target transcriptional regulatory relationships, we generated consensus ‘omics signatures, or consensomes, based on measures of significant differential expression of genomic targets across all underlying transcriptomic experiments. To expose the SPP knowledgebase to researchers, a web browser interface accommodates a variety of routine data mining strategies. Consensomes were validated using alignment with literature-based knowledge, gene target-level integration of transcriptomic and ChIP-Seq data points, and in bench experiments that confirmed previously uncharacterized node-gene target regulatory relationships. SPP is freely accessible at https://beta.signalingpathways.org.

## Introduction

Signal transduction pathways describe functional interdependencies between distinct classes of molecules that collectively determine the response of a given cell to its afferent endocrine, paracrine and cytokine signals ^1^. The bulk of readily accessible information on these pathways resides in the conventional research literature and in knowledgebases that curate such information ^2^. Many such articles are based in part upon discovery-scale datasets documenting, for example the effects of genetic or small molecule perturbations on gene expression in transcriptomic (expression array or RNA-Seq) datasets, and DNA promoter occupancy in cistromic (ChIP-Seq) datasets. Conventionally, only a small fraction of data points from such datasets are characterized in any level of detail in associated hypothesis-driven articles. Although the remaining data points in ‘omics datasets possess potential collective re-use value for validating experimental data or gathering evidence to model cellular signaling pathway, the findability, accessibility, interoperability and re-use (FAIR) status of these datasets ^3, 4^ has been historically limited.

We previously described our efforts to biocurate transcriptomic datasets involving genetic or small molecule manipulation of nuclear receptors ^5^. Here we describe here a novel and distinct web knowledgebase, the Signaling Pathways Project (SPP), that enhances the FAIR status of public cell signaling ‘omics datasets along three dimensions. Firstly, SPP encompasses datasets involving genetic and small molecule perturbations of a broad range of cellular signaling pathway modules - receptors, enzymes, transcription factors and their co-nodes. Secondly, SPP integrates transcriptomic datasets with biocurated ChIP-Seq datasets, documenting genomic occupancy by transcription factors, enzymes and other factors. Thirdly, we have developed a meta-analysis technique that surveys across transcriptomic datasets to generate consensus ranked signatures, referred to as consensomes, which allow for prediction of signaling pathway node-target regulatory relationships. We have validated the consensomes using alignment with literature knowledge, integration of transcriptomic and ChIP-Seq evidence, and using bench experimental use cases that corroborate signaling pathway node-target regulatory relationships predicted by the consensomes. Finally, we have designed a user interface that makes the entire data matrix available for routine data browsing, mining and hypothesis generation by the mammalian cell signaling research community at https://beta.signalingpathways.org.

## Results

### Signaling Pathways Project user interface

Mammalian signal transduction pathways comprise four major categories of pathway module: activated transmembrane or intracellular receptors, which initiate the signals; intracellular enzymes, which propagate and modulate the signals; transcription factors, which give effect to the signals through regulation of gene expression; and co-nodes, a broad variety of classes, such as many transcriptional coregulators ^6^, that do not fall into the other three categories. Fig.1 shows the major signaling pathway module categories, classes and/or families of node and biosamples to which publically archived transcriptomic and ChIP-Seq datasets were mapped. Table 1 shows representative examples of the hierarchical relationships within each of the signaling pathway module categories. Having defined relationships within each major signaling pathway module, we next designed a dataset biocuration pipeline (Supplementary File 1) that would classify publically archived transcriptomic and ChIP-Seq datasets according to the signaling pathway node(s) whose transcriptional functions they were designed to interrogate. To make the results of our biocuration efforts routinely and freely available to the research community, we developed a web interface for the SPP knowledgebase that would provide for browsing of datasets, as well as for mining of the underlying data points. A comprehensive walkthrough file containing instructions on the use of the SPP interface is shown in Supplementary File 2.

**Fig. 1.**
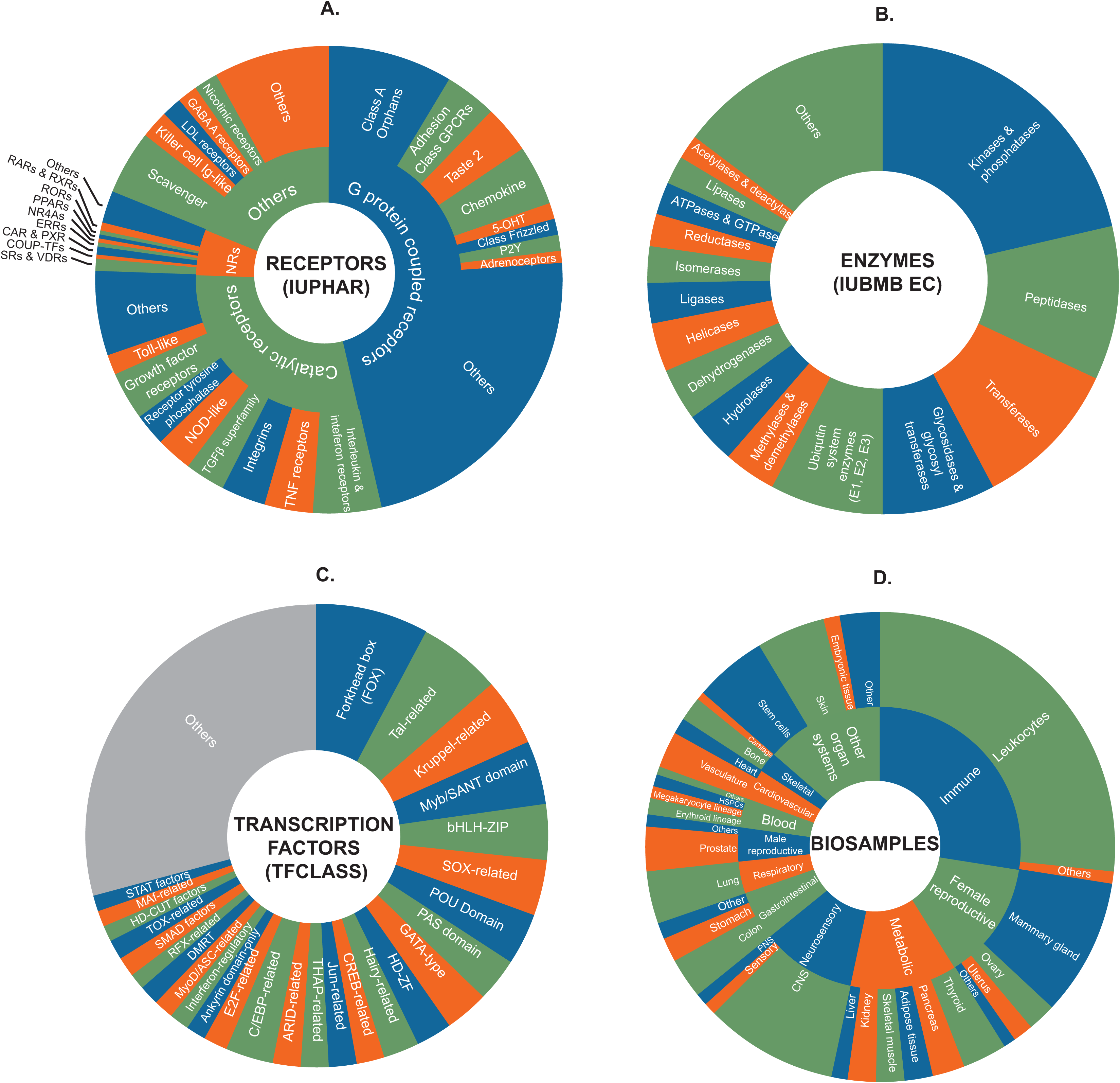
Major signaling pathway module and biosample classifications in the Signaling Pathway Project knowledgebase. Stable community-endorsed classifications for: (A) cellular receptors (International Union of Pharmacology, IUPHAR); (B) enzymes (International Union of Biochemistry and Molecular Biology, IUBMB) and (C) transcription factors (TFClass ^47^) make up the foundation of the Signaling Pathways Project data model. In addition, categorization of tissue and cell line biosamples according to their organ and physiological system of origin (D) facilitates an appreciation of tissue-specific patterns of transcriptional regulation. 5OHTRs, 5 hydroxytryptamine receptors; LDL, low density lipoprotein; NRs, nuclear receptors. For purposes of clarity, omitted from the transcription factors sunburst are factors with > 3 adjacent zinc fingers (482 genes), Hox-related factors (180 genes), multiple dispersed zinc finger factors (140 genes) and other factors with up to three adjacent zinc fingers (24 genes).

**Table 1.**
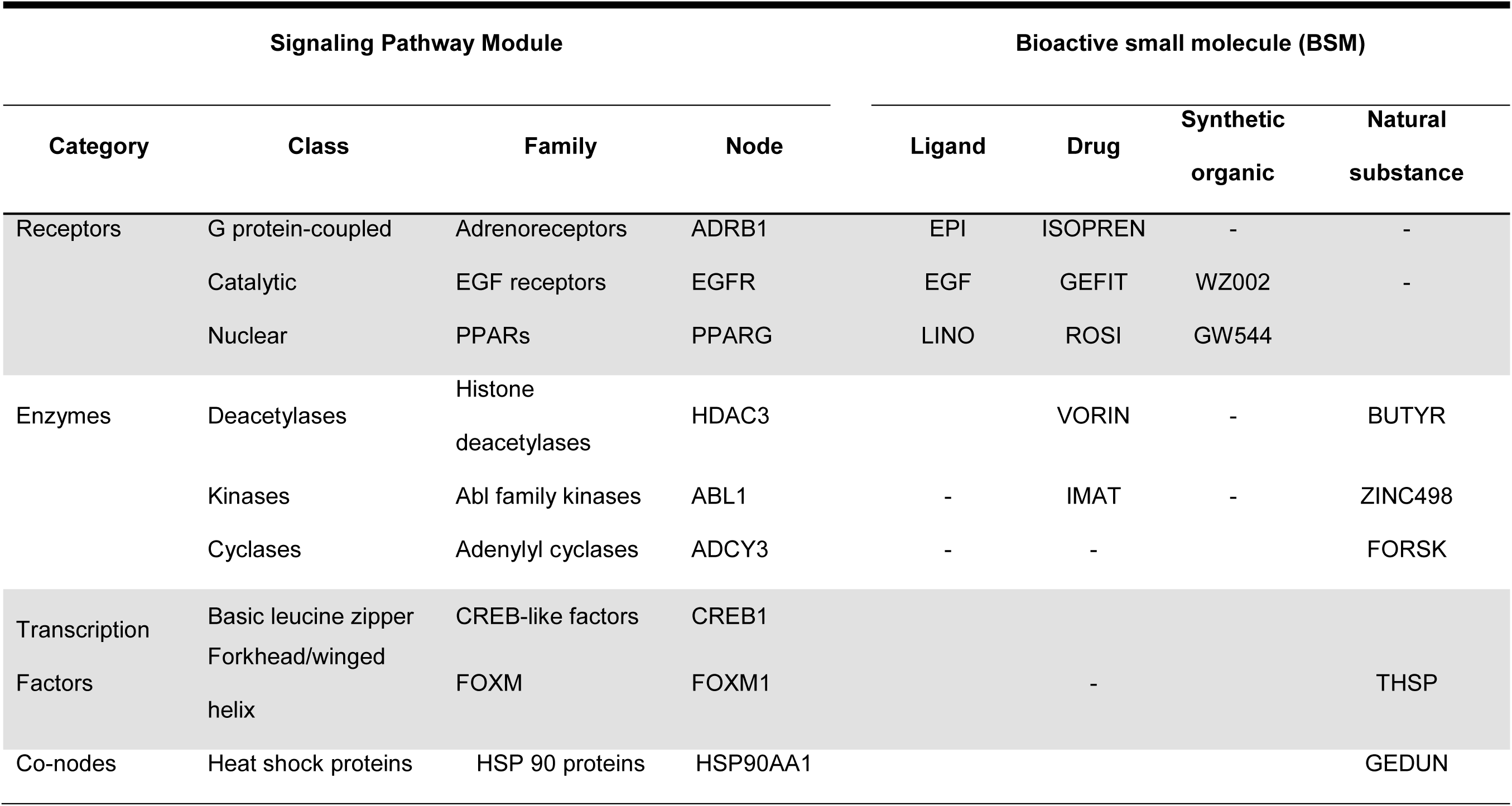
Examples of signaling pathway module hierarchies in the SPP knowledgebase. Nodes and peptide BSMs are represented by approved HGNC symbol for the encoding gene. BSMs are abbreviated as follows: BUTYR, butyric acid; EPI, epinephrine; FORSK, forskolin; GEDUN, gedunin; GEFIT, gefitinb; GW544, GW409544; IMAT, imatinib; ISOPREN, isoprenaline; LINO, linoleic acid; ROSI, rosiglitazone; THSP, thrombospondin; VORIN, vorinistat; WZ002, WZ4002; ZINC498, ZINC08764498;

### Browsing of SPP datasets

The full dataset listing (https://beta.signalingpathways.org/datasets/index.jsf) can be filtered using any combination of: ‘omics dataset type; SPP category (receptor, enzyme, transcription factor, co-node); class or family; biosample physiological system and organ; or species. Individual dataset pages enable integration of SPP with the research literature via digital object identifier (DOI)-driven links from external sites, as well as for citation of datasets to enhance their FAIR status ^3, 4^.

### Mining of SPP datasets in Ominer

The SPP query interface, Ominer, allows a user to specify single gene target, GO term or a custom gene list in the “Gene(s) of Interest” drop-down, and to dial in additional node and biosample regulatory parameters in subsequent drop-down menus as required (Supplementary File 3A). Examples of single gene and GO term queries are shown in Table 2 and Table 3, respectively. Results are returned in an interface referred to as the Regulation Report, a detailed graphical summary of evidence for transcriptional regulatory relationships between signaling pathway nodes and genomic target(s) of interest (Supplementary File 3B & C). To allow users to share links to SPP Regulation Reports with colleagues, or to embed them in research manuscripts or grant applications, the Reports are accessed by a constructed URL defining all of the individual query parameters. The vertical organization of the default Category view in both transcriptomic and cistromic/ChIP-Seq Regulation Reports reflects conventional schematic depictions of cellular signaling pathways, with Receptors on top, followed by Enzymes, Transcription Factors, followed by Co-nodes (Supplementary File 3B & C).

**Table 2:** Examples of Single Gene queries in the Signaling Pathways Project knowledgebase. The Ominer query form accommodates any level of detail required, from broad discovery queries across multiple nodes and organs to specific regulatory contexts at more stringent differential expression or significance cut-offs.

| 'Omics Category | Signaling Pathway Module |  |  | Biosample |  | Target |
| --- | --- | --- | --- | --- | --- | --- |
|  | Category | Class | Family | System | Organ |  |
| Transcriptomic | All |  |  | All |  | PDK4 |
|  | All |  |  | Metabolic | Liver | ALDH3A2 |
|  | Receptor | Catalytic | Insulin receptor | All |  | PFKFB3 |
|  | Enzymes | Kinases | Cyclin-dependent | All |  | DHRS1 |
| ChIP-Seq | All |  |  | All |  | LHPP |
|  | All |  |  | Male Repro | Prostate | TMPRSS2 |
|  | Enzymes | Acetyltransferase | CBP/p300 | All |  | MAMDC2 |
|  | Transcription factors | BZIP | C/EBP family | Immune | Leukocytes | TRIB1 |

**Table 3:**
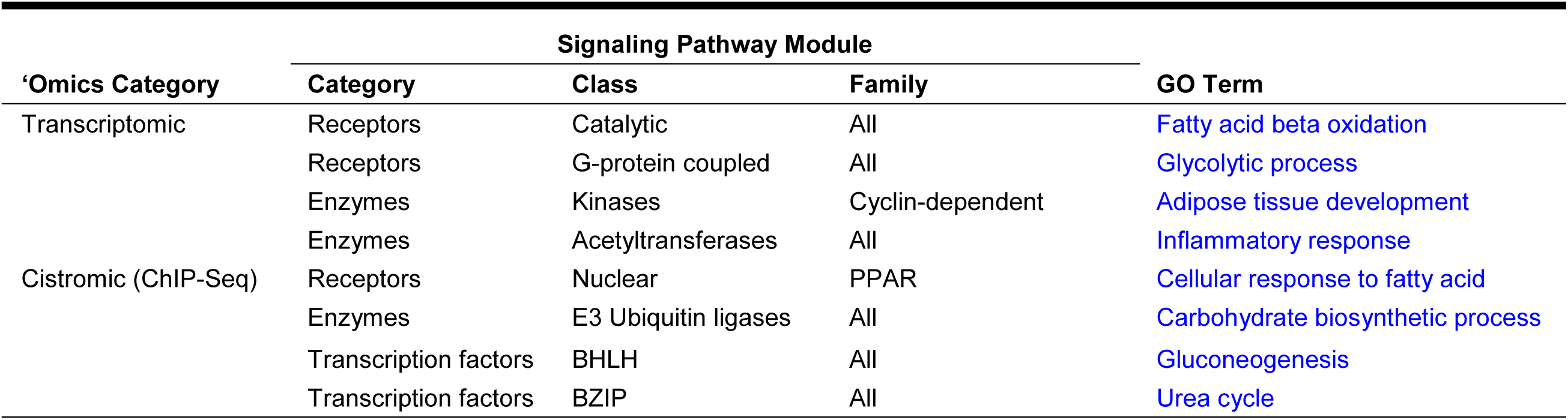
Examples of GO Term queries in the SPP knowledgebase.

Reflecting the hierarchy in Table 1, each Regulation Report category is subdivided into classes (depicted as **Category | Class** in the UI, Supplementary File 3B & C) which are in turn subdivided into families, which in turn contain member nodes, which are themselves mapped to BSMs (Supplementary File 3B (transcriptomic) & 3C (cistromic/ChIP-Seq)). The transcriptomic Regulation Report displays differential expression levels of a given target in experiments involving genetic (rows labelled with italicized node AGS) or BSM (rows labelled with bold BSM symbol) manipulations of nodes within a given family (Supplementary File 3B). Below the node sections, the transcriptomic Regulation Report contains sections in which data points from related animal and cell model experiments are consolidated to convey evidence for previously underappreciated roles of a target transcript in specific physiological contexts. To accommodate users seeking a perspective on regulation of a target in a specific organ, tissue, cell line or species, users can select the “Biosample” and “Species” views from the dropdown. The cistromics/ChIP-Seq Report displays MACS2 peak values within 10 kb of a given promoter transcriptional start site (TSS) in ChIP-Seq experiments named using the convention IP Node AGS | **BSM Symbol** | *Other Node AGS* (Supplementary File 3C). To accommodate users seeking a perspective on regulation of a target in a specific organ, tissue, cell line or species, users can select the “Biosample” and “Species” views from the dropdown (Supplementary File 3B).

Each data point in either Regulation Report links to a pop-up window containing the essential experimental information (Supplementary File 3D, upper = transcriptomic, lower = cistromic). This in turn links to a window summarizing the pharmacology of any BSMs used in the experiment (Supplementary File 3E), or a Fold Change Details window that places the experiment in the context of the parent dataset (Supplementary File 3F), linking to the full SPP dataset page and associated journal article. The Fold Change Details window also provides for citation of the dataset, an important element of enhancing the FAIR status of ‘omics datasets ^3, 4^.

### Consensomes: discovering downstream targets of signaling pathway nodes

An ongoing challenge for the cellular signaling bioinformatics research community is the meaningful integration of the universe of ‘omics data points to enable researchers lacking computational expertise to develop focused research hypotheses in a routine and efficient manner. A particularly desirable goal is unbiased meta-analysis to define community consensus reference signatures that allow users to predict regulatory relationships between signaling pathway nodes and their downstream genomic targets. Accordingly, we next set out to design a meta-analysis pipeline that would leverage our biocurational platform to reliably rank signaling pathway node - target gene regulatory relationships in a given biosample context. Since this analysis was designed to establish a consensus for a node or node family across distinct datasets from different laboratories, we referred to it as consensomic analysis, and the resulting node-target rankings as consensomes. A detailed description of the biocurational and statistical methodologies behind transcriptomic and cistromic consensome analysis is provided in the Methods section.

Table 4 shows examples of the consensomes available in the initial version of the SPP knowledgebase. Supplementary File 4 shows the full list of consensomes available in the initial release of SPP. Consensomes are accessed through Ominer, in which the user selects the “Consensome” from “Genes of Interest”, then either “Transcriptomic” or “Cistromic (ChIP-Seq)” from the “’Omics Category” menu (Supplementary File 2A). Subsequent menus allow for selection of specific signaling pathway families, physiological systems or organs of interest, or species. To accommodate researchers interested in a specific physiological system or organ rather than a specific signaling node, consensomes are also calculated across all experiments mapping to a given physiological system (metabolic, skeletal, etc.) and organ (liver, adipose tissue, etc.), providing for identification of targets under the control of a broad spectrum of signaling nodes in those organs. To maximize their distribution, exposure and citation in third party resources, consensomes can also by accessed by direct DOI-resolved links.

**Table 4:**
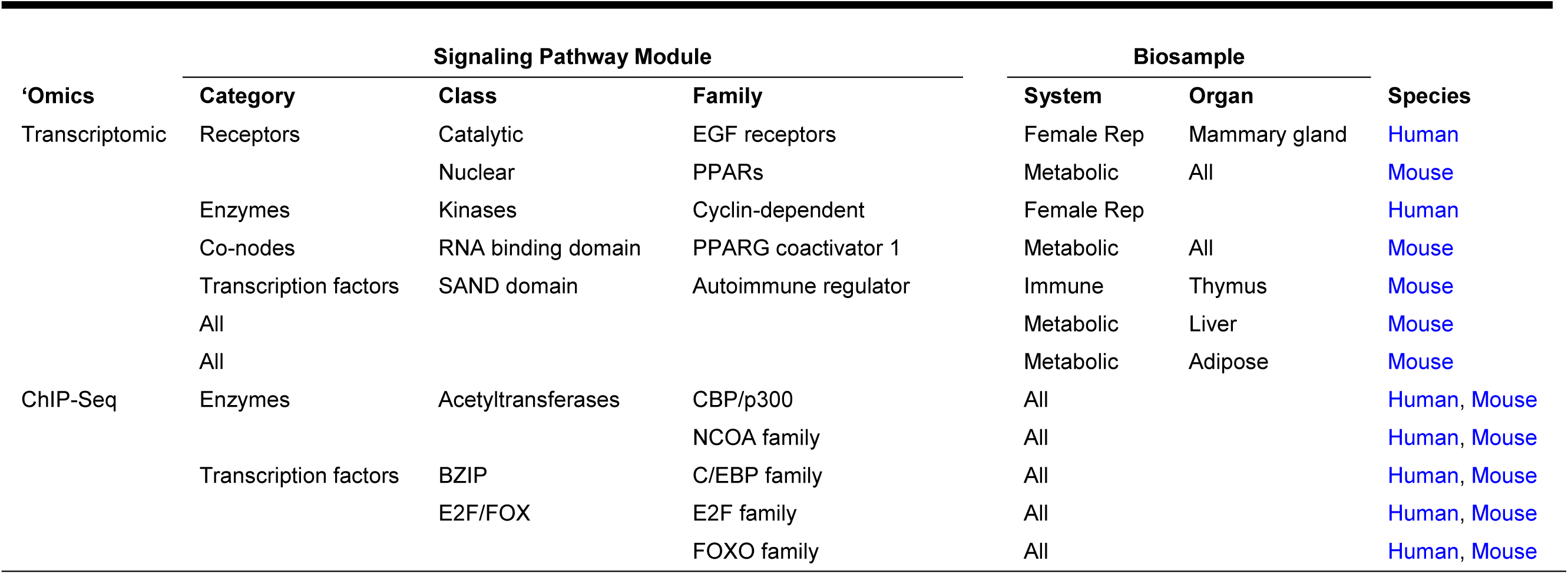
Examples of consensomes in the SPP knowledgebase. Consensomes are calculated at either: the signaling pathway node family level, ranking targets based on their transcriptional sensitivity to manipulation of nodes in a given gene family; or across all experiments in a given organ biosample, to indicate frequently regulated targets in a given organ.

Consensomes are displayed in an accessible tabular format (Supplementary File 3G) in which the default ranking is in ascending order of consensome p-value (CPV; see Methods), although targets can be ranked by any column desired. To reflect the frequency of differential expression of a target relative to others in a given consensome, the percentile ranking of each target within the consensome is displayed. Targets in the 90th percentile of a given consensome – the highest confidence predicted genomic targets for a given node family - are accessible through the web interface, and the entire list of targets is available for download in spreadsheet format for import into custom analysis programs. As previously discussed, to suppress the diversity of experimental designs as a confounding variable in consensome analysis, the direction of differential expression is omitted when calculating the ranked signatures. An appreciation of the pharmacology of a specific node-target gene relationship is essential however to allow researchers to place the ranking in a specific biological context and to design subsequent experiments in an informed manner. To accommodate this, consensome targets link to transcriptomic or cistromic Regulation Reports filtered to display those data points that contributed to the calculation of the CPV for each target.

### Validation of consensomes

We next wished to verify that consensomes were reliable reference datasets for modeling regulatory relationships between cellular signaling pathway nodes and their downstream genomic targets. To do this we designed a validation strategy comprising four components: comparison of consensomes with existing canonical (i.e. literature-defined) node-target relationships; reciprocal validation of node-target relationships between transcriptomic and ChIP-Seq consensomes; validation of pan-node organ consensomes; and in three experimental use cases that functionally validate predicted node-target relationships.

### Canonical signaling node downstream targets are highly ranked in transcriptomic and cistromic nuclear receptor consensomes

Two considerations recommended members of the nuclear receptor (NR) superfamily of physiological ligand-regulated transcription factors for selection for initial proof-of-principle validation of the consensomes. Firstly, as the largest single class of drug targets, they are the subject of a large body of dedicated research literature, affording considerable opportunity for testing the consensomes against existing canonical knowledge of their downstream targets. Secondly, as ligand-regulated transcription factors, members of this superfamily are prominently represented in both publically archived transcriptomic and ChIP-Seq experiments, enabling meaningful cross-validation of consensomes between these two experimental categories. We selected the ten top ranked targets in the following consensomes: estrogen receptors in human mammary gland (ERs-Hs-MG); the androgen receptor in human prostate gland (AR-Hs-Prostate); the glucocorticoid receptor in mouse liver (GR-Mm-Liver); and the peroxisome proliferator-activated receptor (PPAR) family in the mouse metabolic system (PPARs-Mm-Metabolic). Encouragingly, we found that 36/40 (90%) of the most highly ranked targets across all four consensomes had been previously identified as targets of members of those node families in the research literature, and that of these same 40 genes, 82% (33/40) were in the 90th percentile or higher in comparable ChIP-Seq consensomes (Supplementary File 5).

### Frequently regulated hepatic transcripts are enriched for critical regulators of metabolic pathways

Substantial literature evidence indicates that genomic targets encoding hepatic metabolic enzymes are subject to dynamic transcriptional regulation by numerous afferent metabolic and endocrine cues ^7^. If consensome analysis were biologically valid, we anticipated that targets with elevated rankings in the murine hepatic transcriptomic consensome – that is, genes that are preferentially responsive to multiple hepatic signaling pathways - would be enriched for genes encoding enzymes with prominent roles in hepatic metabolism. To test this hypothesis, we first identified genes in the 99^th^ percentile of the All nodes-Mm-liver transcriptomic consensome, which represents the top 1% of genes that are significantly differentially expressed in expression profiling experiments in a murine hepatic biosample, irrespective of the perturbed signaling node (*n* = 258; Supplementary File 6). Based upon a set of 1647 murine metabolic enzymes curated by the Mammalian Metabolic Enzyme Database resource (Corcoran, 2017 #250} we found that 38% (99/258) of the top 1% genes encoded metabolic enzymes, a 6.5-fold enrichment over the frequency of metabolic enzyme-encoding genes in the entire All nodes-Mm-liver transcriptomic consensome (5.7%, 1490/25922).

We next speculated that transcripts under fine control by hepatic signaling pathways would be enriched for enzymes whose deficiency would have a critical impact upon hepatic metabolism. Using the OMIM resource ^8^, we identified a set of unique human genes whose deficiency has a literature-supported connection to a human genetic disorder or trait (*n* = 5277). Using this reference gene set, we established that human orthologs of 42% (41/97) of metabolic enzyme-encoding targets in the 99^th^ percentile of the All Nodes-Mm-liver transcriptomic consensome had documented deficiencies in a human metabolic disorder (Table 5 and Supplementary File 7), compared with a frequency of 15% (3865/25922) for such genes in the entire consensome.

**Table 5.**
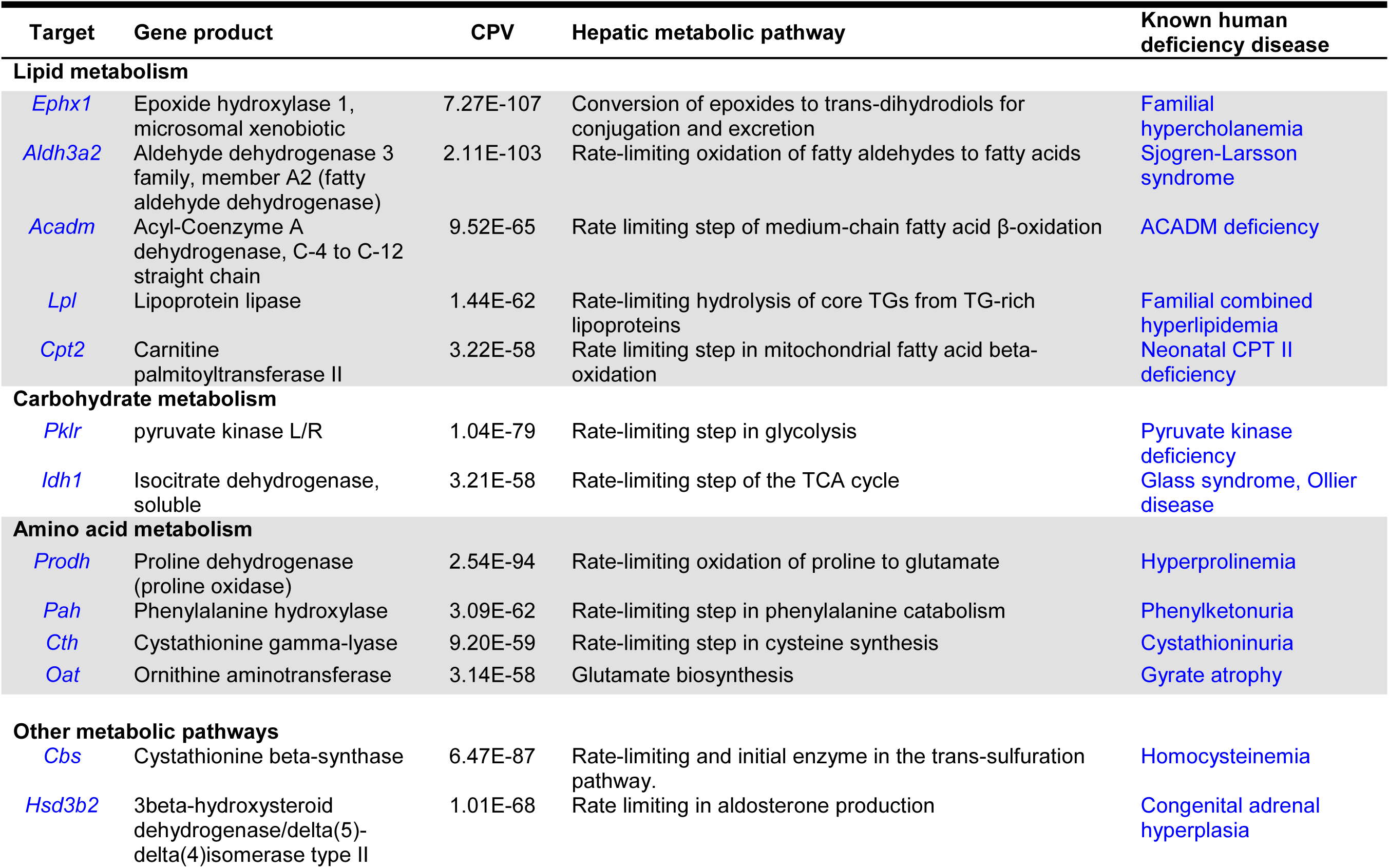
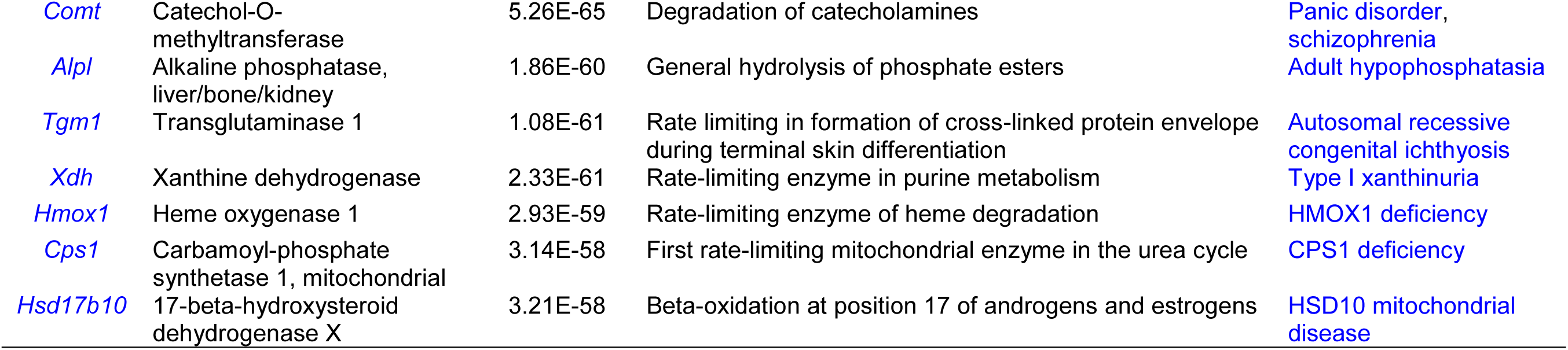
Selected genes encoding metabolic enzymes in the 99^th^ percentile of the All nodes-Mm-liver transcriptomic consensome whose deficiency is associated with a human metabolic disorder. Gene symbol links point to SPP transcriptomic Regulation Reports filtered for mouse liver. Disease links point to OMIM entries highlighted for the corresponding human gene. See Supplementary File 7 for the full list.

Rate-limiting enzymes (RLEs) play critical roles in determining mammalian metabolic flux ^9^. We surmised that hepatic signaling pathways might preferentially target RLE-encoding genes to exert efficient control over hepatic metabolism, and that this would be reflected in the enrichment of RLE genes among enzyme-encoding targets in the top 1% of the All Nodes-Mm-liver transcriptomic consensome. Consistent with this notion, 40/97 (41%) of the metabolic enzymes encoded by targets in this group of genes catalyze rate-limiting steps in the pathways in which they participate (Supplementary File 7), a nearly 3-fold enrichment over the previously published estimate of 14% (96/687) for the proportion of hepatic metabolic enzymes made up by RLEs ^9^. This analysis demonstrates the ability of pan-node organ consensomes to illuminate factors that are downstream targets of multiple distinct signaling nodes in a specific organ and, by inference, have pivotal, tightly-regulated roles in the function of that organ.

We next wished to establish whether the biological significance implied by elevated rankings in consensomes for cellular signaling pathway nodes was reflected in both gain- and loss-of-function validation experiments at the bench.

### Bench validation use case 1: elevated consensome rankings predict functional roles for targets in signaling node pathways

In the first use case, we used Q-PCR analysis to verify ER-dependent regulation of a panel of both characterized and uncharacterized ER targets that were highly ranked in the ERs-Hs-All consensome (Fig. 2A). We next identified targets that were assigned very high consensome rankings, but whose functional importance in the context of signaling by the corresponding signaling nodes was previously uncharacterized in the research literature. The tumor protein D52-like 1 (*TPD52L1*) gene encodes a little-studied protein that bears sequence homology to members of the TPD52 family of coiled-coil motif proteins that are overexpressed in a variety of cancers ^10^. Despite a ranking in the transcriptomic (ERs-Hs-All-TC CPV = <1E-130, 99.99^th^ percentile) and ChIP-Seq (ERs-Hs-All-CC, 99^th^ percentile) ER consensomes that was comparable to or exceeded that of canonical ER target genes such as *GREB1* or *MYC*, and subsequent experimental bench validation of the ER family-*TPD52L1* regulatory relationship (Fig. 2A), the functional role of *TPD52L1* in ER signaling has gone unexplored in the research literature. Suggestive of a role for TPD52L1 in ER regulation of cell division, we identified 17BE2-dependent association of TPD52L1 with structures resembling stress fibers (Fig. 2B), which play an important role in mitosis orientation during cell division ^11^, and found that depletion of *TPD52L1* resulted in a significant decrease in 17BE2-induced proliferation of MCF-7 cells (Fig. 2C).

**Fig. 2.**
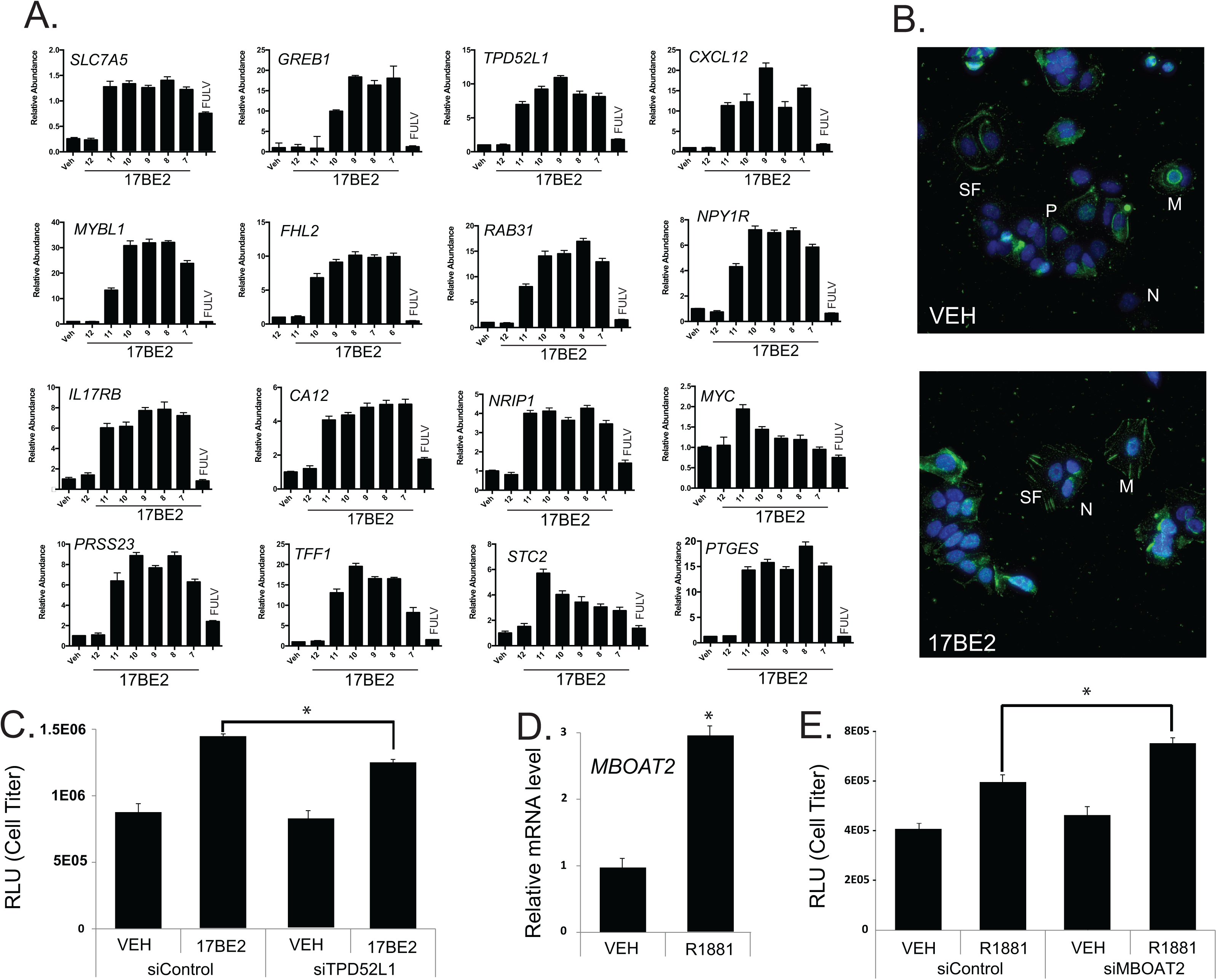
Validation of ER regulation of *TPD52L1* (A-C) and AR regulation of *MBOAT2* (D, E) A) Q-PCR analysis of dose dependent induction by 17BE2 in MCF-7 cells of targets with elevated rankings in the ER-Hs-MG transcriptomic consensome. Cells were treated for 18 h with varying concentrations of 17BE2 alone or 1 nM 17BE2 in combination with 100 nM of the selective ER downregulator FULV. Consistent with the strong ER family node dependence of regulation predicted by the ER-Hs-mammary gland transcriptomic and ChIP-Seq consensomes, FULV completely abolishes 17BE2 induction of all target genes tested. Each number is representative of –log[17BE2] such that the number 9 is equivalent to 1 nM 17BE2. Data are representative of three independent experiments. B) MCF-7 cells were immunolabeled with TPD52L1 antibody (green) and imaged by deconvolution widefield microscopy. Images shown are max intensity projections, where DAPI (blue) stains DNA. Scale bar is 10μm. (in the inset, 5μm). M, membrane; N, nucleus; P, perinuclear junctions; SF, stress fibers. C. Depletion of TPD52L1 restricts MCF-7 cell viability. D. Induction of *MBOAT2* in LNCaP prostate epithelial cells upon treatment with 0.1 nM R1881. E. AR-stimulated viability of LNCaP cells is enhanced by depletion of *MBOAT2.* Cells were harvested on Day 5. Gene expression of *KLK3* and *FKBP5*, known canonical AR target genes, was slightly reduced or unaffected, respectively, by *MBOAT2* siRNA knockdown (data not shown). Statistical significance was determined using PRISM by One-way ANOVA with Tukey’s multiple comparison test. * p < 1E-04.

*MBOAT2* encodes an enzyme catalyzing cycles of glycerophospholipid deacylation and reacylation to modulate plasma membrane phospholipid asymmetry and diversity ^12^. *MBOAT2* has rankings in both the AR-Hs-All TC (CPV = 3.7E-38, 99.96^th^ percentile) and ChIP-Seq (99^th^ percentile) consensomes comparable to those of canonical AR target genes such as *KLK3* and *TMPRSS2*. In contrast to the large volume of literature devoted to these targets however, with the exception of a mention in a couple of androgen expression profiling studies ^13, 14^, the functional role of MBOAT2 in the context of AR signaling has been entirely unstudied. In validation of its elevated AR consensome ranking, we confirmed that *MBOAT2* was an AR-regulated gene in cultured prostate cancer cell lines (Fig. 2D), and that depletion of *MBOAT2* significantly increased LNCaP cell numbers at growth day 5 in in R1881-treated celIs, but not untreated cells (Fig. 2E).

### Bench validation use case 2: GR, ERR family members and insulin receptor regulate targets encoding glycogen synthase phosphatase and kinase regulatory subunits

The first two experimental validation studies focused on distinct single node-target regulatory relationships. We next wished to validate the use of consensome intersection analysis to highlight convergence of multiple signaling nodes on targets involved in a common downstream biological process. Glycogen synthase, which catalyzes the rate-limiting step in the interconversion of glucose and glycogen in metabolic organs, is subject to tandem activation by protein phosphatase 1 (PP1), ^15^, and deactivation by 5’AMP-activated protein kinase (AMPK} ^16^. Although regulation of glycogen metabolism in a variety of organs is known to be under the control of signaling mediated by the glucocorticoid (GR) ^17^, estrogen receptor-related (ERR) ^18^ and insulin (IR) ^19^ receptor families, the respective underlying mechanisms are incompletely understood. We wished to use SPP consensomes to investigate the hypothesis that regulation of glycogen metabolism by members of these distinct receptor families might involve convergent regulation of glycogen synthase activity. Surveying *p* < 0.05 genes in the mouse ERR and GR and human IR transcriptomic consensomes mapping to the GO terms “AMP-activated protein kinase activity” or “protein phosphatase regulator activity”, we isolated two targets, *Ppp1r3c* (GR-Mm-All CPV = 1.89E-15; ERR-Mm-All CPV 5.98E-06) and *Prkab2* (ERR-Mm-All CPV =1.84E-04, IR-Hs-All CPV = 4.9E-04). These encode, respectively, the PTG regulatory subunit of the PP1 holoenzyme ^20, 21^, and the AMPKβ2 regulatory subunit of the AMPK holoenzyme ^22, 23^. Corroborating these predicted regulatory relationships, we identified conserved GR and ERR response elements in the *Ppp1r3c* promoter (Supplementary File 8, Use Case 2). We also confirmed that *Ppp1r3c* was a target of GR in mouse Hepa-1-c liver cells (Fig. 3A) and that both *Ppp1r3c* and *Prkab2* were transcriptionally regulated in response to either loss- or gain-of function of Esrra in skeletal muscle (Fig. 3B, C & D, left panel). Finally, consistent with inhibition of AMPK activity by IR signaling ^24^, treatment of myoblasts for 24 h with the IR agonist IGF1 stimulated the *Prkab2* promoter, an effect that was further enhanced by expression of both Esrra and Esrrg isoforms (Fig. 3D, right panel).

**Fig. 3.**
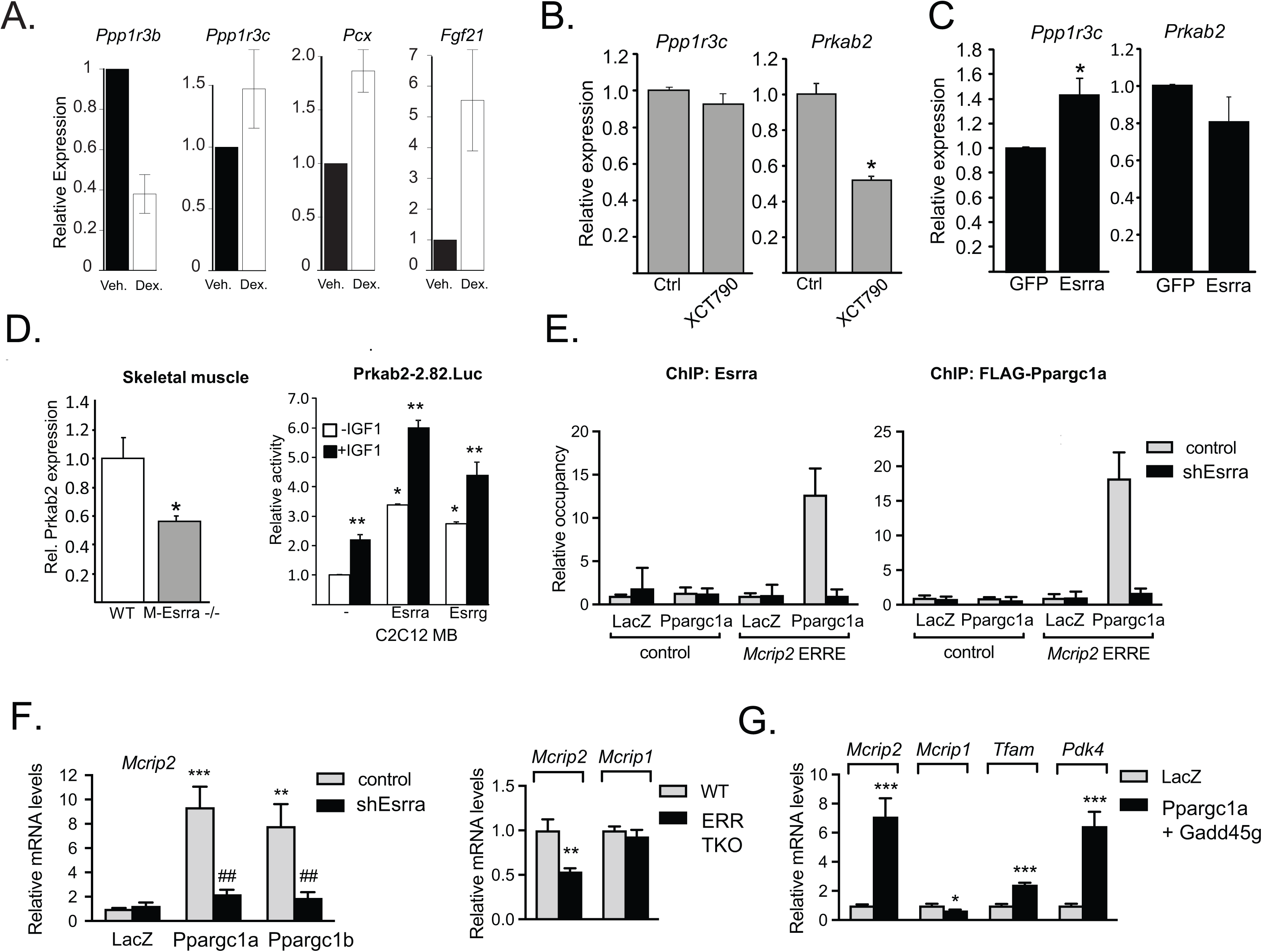
Validation of consensome predictions of genomic targets for GR, ERR and IR. **A**. *Ppp1r3c* is regulated by GR. Mouse Hepa-1-c hepatoma cells were treated with 250 nM dexamethasone (DEX) for 48 h, followed by qPCR of glycogenic genes including *Ppp1r3c*, *Ppp1r3b* and the established GR/NR3C1 targets pyruvate carboxylase (*Pcx*) and *Fgf21*. **B**. Endogenous *Ppp1r3c* and *Prkab2* transcripts were measured by quantitative real-time PCR in C2C12 day 3 myotubes treated with vehicle or 5 μM of the Esrra inverse agonist XCT790 (IC50∼0.5mM) for 24 h. **C.** Endogenous *Ppp1r3c* and *Prkab2* transcripts were measured by quantitative real-time PCR in C2C12 day 3 myotubes transduced with recombinant adenovirus expressing GFP or human ERRα. Experimental transcript levels were normalized to 36B4 expression and results are expressed as the mean ± S.E.M. Asterisks * indicate significant difference vehicle vs. treatment groups, (p ≤ 0.05, n=3). **D.** Left panel. Expression of *Prkab2* transcript is reduced by 40% in Esrra-depleted skeletal muscle compared to wild-type tissue. Endogenous *Prkab2* expression was assayed by Q-PCR in vastus lateralis muscles of male wild-type (WT) or Esrr1-/- mice (ERRα-/-). *Prkab2* transcript was normalized to 36B4 expression and results are expressed as the mean (± S.E.M). Asterisk * indicates significant difference between groups (p < 0.05, n=4). Right panel. Activity of the Prkab2.-2.82.Luc promoter-reporter in C2C12 myoblasts (MB) cotransfected with vector, ERRα/Esrra or ERRγ/Esrrg, as indicated. One day post-transfection MB were then cultured in 0.1% FBS overnight -/+ 10nM IGF1 treatment for 24 hours. Data are reported as mean luciferase/renilla values normalized to control (± S.E.M.) for three trials. Asterisks indicate significant differences between transfection conditions (*) or IGF1 treatment (**), (p ≤ 0.05, n=3). **E.** ChIP of Esrra (left panel) and Ppargc1a (right panel) at the Mcrip2 ERRE. C2C12 myotubes were treated as described in Methods. Relative occupancy represents the amount of Mcrip2 DNA (or of a control genomic region that has no ERR binding sites) that is immunoprecipitated by anti-ERRa or anti-Flag (detecting the Flag-tagged Ppargc1a in the different myotubes, relative to the DNA immunoprecipitated in LacZ/control shRNA cells (which has been set as 1 for each DNA region). Data are mean ± SD (n=3). **F. Left panel**. *Mcrip2* is induced by Ppargc1 co-nodes in C2C12 myotubes in an Esrra-dependent manner. RNA (isolated 24 hrs after Ppargc1a/b expression) was analyzed by RT-qPCR. Data are normalized to 36B4, and expressed relative to levels in LacZ/shGFP cells. **Right panel.** Expression levels of *Mcrip2*, but not the related gene *Mcrip1*, are decreased in primary brown adipocytes lacking Esrra, Esrrb and Esrrg, relative to ERR WT mice. mRNA levels were determined as in left panel. **G.** Simulation of chronic adrenergic stimulation of primary brown adipocytes by overexpression of Ppargc1a and Gadd45g significantly increases expression of *Mcrip2* relative to mock-transfected adipocytes. Included as controls are the OXPHOS genes *Tfam* and Pdk4, encoding pyruvate dehydrogenase kinase isoform 4, a characterized ERR target and the third highest ranked target in the All nodes-Mm-adipose transcriptomic consensome signature. Differentiated adipocytes were infected with adenoviruses expressing Ppargc1a and Gadd45g and mRNA levels measured as described in the Methods section.

### Bench validation use case 3: the murine ERR, PPARGC and adipose tissue consensomes implicate *Mcrip2* in adipocyte oxidative metabolism

The control of cellular mitochondrial content and oxidative capacity is important for cellular and organismal energy homeostasis ^25^. For example, brown and beige adipocytes generate new mitochondria and increase their oxidative and thermogenic capacity in response to norepinephrine (NE), which is secreted locally when the organism senses a cold environment ^26^. NE (adrenergic) stimulation elicits an acute transcriptional response, exemplified by the induction of genes such as the uncoupling protein *Ucp1*, the Pparg co-node *Ppargc1a* and the signaling regulator *Gadd45g* ^26, 27^. *In vivo*, chronic or repeated exposure to cold (or to adrenergic agonists) also leads to higher mitochondrial DNA content, increased cristae density and enhanced expression of oxidative enzymes (OxPhos complexes) and *Ucp1* ^26^.

Mitochondrial biogenesis is regulated by a variety of nuclear receptors, including members of the ERR family, as well as Pparg and its co-nodes Pppargc1 and Ppargc1b, members of the PPARGC family of RNA-binding transcriptional coregulators ^28^. A highly ranked gene in the ERRs-Mm-All transcriptomic consensome was *Mcrip2* (CPV = 1.54E-12), which has no literature-characterized function or role other than a report identifying it as an interacting partner of Ddx6 that was localized to RNA stress granules ^29^. Interestingly, we noted that *Mcrip2* was also very highly ranked in the All nodes-Mm-adipose (CPV = 6.58E-37), PPARs-Mm-Adipose (CPV = 1.17E-30), PPARGCs-Mm-Metabolic (CPV = 1.96E-06) and All nodes-Mm-liver (CPV = 2.2E-88) transcriptomic consensomes, indicating an influential role in adipose biology, as well as a potentially broader function in systemic metabolic homeostasis. Inspection of the *Mcrip2* transcriptomic and cistromic Regulation Reports indicated that it was regulated under conditions of mitochondrial biogenesis in adipose tissue, as well as loss and gain of function of a variety of known signaling node regulators of mitochondrial biogenesis. Corroborating this evidence, we identified a conserved consensus ERR binding site in the first intron of *Mcrip2* (Supplementary File 9, Use Case 3) and confirmed that Esrra and Ppargc1a were recruited to the *Mcrip2* ERRE (Fig. 3E). We also confirmed the interdependence of Esrra, Ppargc1a and Ppargc1b in regulation of *Mcrip2* in mouse muscle cells (Fig. 3F; Supplementary File 7, Use Case 3), and brown adipocytes (Fig. 3F), and demonstrated induction of *Mcrip2* in response to conditions mimicking chronic adrenergic stimulation of brown adipocytes (Fig. 3G).

## Discussion

Effective re-use of ‘omics datasets in the field of cellular signaling relies upon the ability of bench researchers to ask sophisticated questions across this data universe in a routine manner. Many excellent tools and resources have been developed in the field of cell signaling ‘omics ^30–41^. Here, we set out to complement these resources to allow researches to routinely answer questions such as: What cell cycle-related factors are regulated by FGF receptors in the liver? What downstream targets are most responsive to insulin receptor signaling in the liver? What E3 ubiquitin ligases regulate targets in my gene set of interest? To fill this gap we designed a knowledgebase, SPP, which allows bench researchers to routinely evaluate public transcriptomic or ChIP-Seq dataset evidence for regulatory relationships between cellular signaling pathway nodes and their downstream targets.

The SPP resource is characterized by a number of unique features. Previous transcriptomic meta-analysis approaches in the field of cellular signaling have been perturbation-centric, and applied to experiments involving a single unique perturbant ^42, 43^. Consensomic analysis differs from these approaches in that it is node-centric: that is, it is predicated upon the functional relatedness of *any* genetic or small molecule manipulation of a given pathway node, and allows experiments to be grouped for meta-analysis accordingly. In doing so, it lends the meta-analysis greater statistical power, and calls potential node-target relationships with a higher degree of confidence than would otherwise be possible. An additional unique aspect is that many other primary analysis and meta-analysis studies describing integration of transcriptomic and ChIP-Seq datasets, although insightful, are limited in scope and exist only as stand-alone literature studies. Ours is to our knowledge the first meta-analysis to be sustainably integrated into an actively-biocurated public web resource in a manner supporting routine use by researchers lacking formal informatics training.

Our resource has a number of limitations. SPP is currently based upon transcriptomic and ChIP-Seq data since these are the most numerous and informatically mature of the various types of ‘omics data. Future versions of the knowledgebase will incorporate datasets from additional ‘omics modalities such as metabolomics, as these accrue in greater numbers and their data stewardship matures. Secondly, bias in publically archived datasets towards specific nodes and biosamples is to some extent reflected in SPP. Other limitations of the consensomes relate to the design of available archived experiments. For example, certain targets may be regulated by a given node only under specific circumstances (e.g. acute BSM administration, or in a specific organ or tissue) and if such experiments do not exist or are otherwise publically unavailable, these targets would not rank highly in the corresponding node consensome. Moreover, a low ranking for a target in a consensome does not necessarily imply the complete absence of a regulatory relationship, and may reflect the requirement for a quite specific cellular context (e.g. specific organ) for such regulation to take place. Finally, since SPP is based upon transcriptional methodologies, effects exerted by signaling pathway nodes at the protein level, such as enhanced stabilization or degradation of protein, or modulation of the rate of translation, will not be reflected at the mRNA level. Caveats such as these notwithstanding, we believe SPP adds value to the currently available tool set enabling bench investigators to re-use archived datasets for hypothesis generation and data validation.

## Acknowledgements

This work was supported by the National Institute of Diabetes, Digestive and Kidney Diseases (DK097748, DK48807, DK107535, DK56338, DK095686 and DK105126), the National Institute of Child Health and Development (DK097748), the National Cancer Institute (CA125123) and the National Heart, Lung and Blood Institute (HL127624). Additional funding was provided by the Dan L. Duncan NCI Comprehensive Cancer Center at Baylor College of Medicine. Microscopic analysis was supported by the Integrated Microscopy Core at Baylor College of Medicine with funding from the Cancer Prevention Research Institute of Texas (CPRIT, RP150578) and the John S. Dunn Gulf Coast Consortium for Chemical Genomics. We thank Dr Bethany Hazen and Dr Erin Brown for technical help. Finally we extend our thanks to all investigators who archived their ‘omics datasets, without whom the Signaling Pathways Project would not be possible.

## Author contributions

**Knowledgebase concept and design**: NM, SO

**Consensome algorithm design**: SH, NM, SO

**Data processing:** SO, ZW, AM

**Biocuration:** NJM, SAO, DA, KM

**Software development:** LB, AM, WD, WK

**Database and system administration**: MD, SQ

**Bench validation:** KA, RH, YC, AH, MG, DB, JH, FS, CF, AK, DM

**Manuscript drafting:** NM, DB, JH, CF, AK, DM

## Data availability statement

The Signaling Pathways Project is freely accessible at https://beta.signalingpathways.org. Programmatic access to underlying data points and their associated metadata are is supported by a RESTful API at https://beta.signalingpathways.org/docs/. All Signaling Pathways Project datasets are biocurated versions of publically archived datasets, are formatted according to the recommendations of the FORCE11 Joint Declaration on Data Citation Principles, and are made available under a Creative Commons CC 3.0 BY license. The original datasets are available are linked to from the corresponding SPP datasets using the original repository accession identifiers. These identifiers are for transcriptomic datasets, the Gene Expression Omnibus (GEO) Series (GSE); and for cistromic/ChIP-Seq datasets, the NCBI Sequence Read Archive (SRA) study identifier (SRP).

*Note for reviewers: Per discussion with Dr Andrew Hufton, the requirement for the availability of full download of SPP site content at initial submission has been waived. We commit to full availability of this functionality as a condition in the event of publication.*

## Source code availability statement

*Note for reviewers: Per discussion with Dr Andrew Hufton, the requirement for the availability of SPP source code at initial submission has been waived. We commit to full availability of SPP source code as a condition in the event of publication.*

## Methods

### Data model design

The goal of the Signaling Pathways Project (SPP) is to give bench scientists routine access to biocurated public transcriptomic and ChIP-Seq datasets to infer or validate cellular signaling pathways operating within their biological system of interest. Although such pathways are diverse and dynamic in nature, they typically describe functional interdependencies between molecules belonging to three major categories of pathway module: activated transmembrane or intracellular receptors, which initiate the signals; intracellular enzymes, which propagate and modulate the signals; and transcription factors, which give effect to the signals through regulation of gene expression ^44^. Accordingly, we first set out to design a knowledgebase that would reflect this modular architecture. To ensure that our efforts were broadly aligned with established community standards, we started by integrating existing, mature classifications for receptors (International Union of Pharmacology, IUPHAR; ^45^), enzymes (International Union of Biochemistry and Molecular Biology Enzyme Committee ^46^) and transcription factors (TFClass ^47^). Table 1 shows representative examples of the hierarchical relationships within each of the signaling pathway module categories. To harmonize and facilitate data mining across different signaling pathway modules, top level categories were subdivided firstly into functional classes, which in turn were subdivided into gene families, to which individual gene products were assigned. Fig. 1A-C summarizes the major classes and/or families in each category, collectively comprising 174 families of receptors, 616 families of enzymes and 371 families of transcription factors. Note that some families contain only a single known node. Fig.1D summarizes the hierarchy of physiological systems and organs into which experimental biosamples (tissues and cultured or primary cell lines) were classified. Consistent with terminology in use in the cellular signaling field ^1, 48^, we refer to these individual gene products as nodes. Molecular classes that are relevant to cellular signaling pathways but do not fall into any of the three categories referred to above, such as regulatory RNAs, chromatin factors and cytoskeletal components, were assigned to a Co-nodes category. Impacting the functions of nodes in all four categories are bioactive small molecules (BSMs), encompassing: physiological ligands for receptors; prescription drugs, targeting almost exclusively nodes in the receptor and enzyme categories; synthetic organics, representing experimental compounds and environmental toxicants; and natural products (Table 1). BSM-node mappings were retrieved from an existing pharmacology biocuration initiative, the IUPHAR Guide To Pharmacology ^45^, or annotated by SPP biocurators *de novo* with reference to a specific PubMed identifier (PMID).

### Dataset biocuration

For knowledgebase design purposes, we defined a dataset as a collection of individual experiments encompassed by a specific GEO series (GSE, for transcriptomic datasets) or SRA Project (SRP, for ChIP-Seq datasets).

#### Transcriptomic datasets

We previously described our efforts to biocurate Gene Expression Omnibus (GEO) transcriptomic datasets pertinent to nuclear receptor signaling as part of the Nuclear Receptor Signaling Atlas ^5^. In order to expand this collection to encompass datasets involving perturbation of the full range of signaling pathway nodes, we carried out a systematic survey of Gene Expression Omnibus and the research literature to identify an initial population of transcriptomic datasets representing a reasonable cross-section of the various classes of signaling pathway node referred to in Figure 1. We next carried out a three step QC check to filter for datasets that (i) included all files required to calculate gene differential expression values; (ii) contained biological replicates to allow for calculation of associated significance values; and (iii) whose samples clustered appropriately by principal component analysis. Typically, 20-25% of archived transcriptomic datasets were discarded at this step for failure to meet one or more of the above criteria.

The remaining datasets were diverse in design, typically involving genetic (single or multi-node node overexpression, knockdown, knockin or knockout) or BSM (physiological ligand, drug or synthetic organic or natural substance; single or multi-BSM; time course; agonist, antagonist or tissue-selective modulator) manipulation of a signaling node across a broad range of human, mouse and rat biosamples. To maximize the amount of biological information extracted from each transcriptomic dataset, we calculated differential expression values for all possible contrasts, and not just those used by the investigators in their original publications. Next, transcriptomic experiments were mapped where appropriate to approved symbols (AGSs) for human, mouse and rat genes, representing genetically perturbed signaling nodes, and/or to unique identifiers for BSMs, as well as to the previously described biosample controlled vocabulary. Model experiments representing a variety of physiological and metabolic processes (e.g. inflammation, adipogenesis, fasting-fed) were annotated where appropriate. Gene differential expression values were calculated for each experiment using an industry standard Bioconductor pipeline ^5^. Finally, experiments were organized into datasets for which digital object identifiers (DOIs) were minted as previously described ^4^.

#### ChIP-Seq datasets

In addition to integration of transcriptomic datasets with each other, their integration with related ChIP-Seq datasets was desirable since it would provide for cross validation of predicted node-target relationships, as well as providing for more detailed mechanistic modeling of such relationships than would be possible using either omics platform individually. The ChIP-Atlas resource ^49^ supports re-use of ChIP-Seq datasets by carrying out uniform MACS2 peak-calling across ChIP-Seq datasets archived in NCBI’s Short Read Archive (SRA). We therefore next set out to identify and annotate ChIP-Atlas-processed SRA ChIP-Seq datasets relevant to mammalian signaling pathway nodes. Individual SRA experiments were first mapped to SRA Study Identifier (prefix SRP, DRP or ERP), which represents the SPP cistromic dataset unit. Individual SRA experiments (prefix SRX, DRX or ERX) were then mapped to the AGS of the immunoprecipitation (IP) node and any other genetically manipulated nodes (e.g. knockdown or knockout background), to any BSMs represented in the experimental design, and to the biosample in which the experiment was carried out. Experimental data and associated metadata were then loaded into the SPP Oracle 12c database.

### Generation of consensomes

#### Transcriptomic consensomes

Transcriptomic ‘datasets’ are collections of data from different sources (i.e. different GEO datasets). Experiments, or contrasts in statistical terminology, are pairs of conditions (i.e. control and treatment) within data sets, each of which has multiple observations, that are used to generate nominal p-values and fold changes in expression for each gene (target) represented in the pair. These are all pre-computed and stored in the SPP Oracle database. Experiments are the unit of analysis, of which a single dataset can have one or more. differential expression values and associated significance measures were generated from appropriate experimental contrasts in GEO Series as previously described ^50^.

Large scale meta-analysis pipeline of publically archived transcriptomic datasets is confronted primarily by the sheer heterogeneity of genetic and pharmacological perturbation designs represented in these datasets. We hypothesized that irrespective of the nature of the perturbation impacting a given pathway node, downstream targets with a greater dependence on the integrity of that node would be more likely to be differentially expressed in response to its perturbation than those with a weaker regulatory relationship with the node. Accordingly, to enhance the statistical power of the analysis, we initially binned transcriptomic experiments for meta-analysis on the basis of genetic or pharmacological manipulation of a given signaling node. To further extend statistical power, experiments involving manipulation of all nodes in a defined gene family were combined for meta-analysis. Next, we further classified experiments according to the biosample and species in which they were carried out, prior to committing them to the consensome pipeline as described before.

#### Computation of target & experiment-specific nominal p-values & fold changes

Although RNA-Seq datasets are growing in number, expression arrays remain in use and the vast majority of expression profiling datasets archived in Gene Expression Omnibus are on array platforms. We first therefore set out to develop an algorithm that would establish consensus across array datasets. Although much less than 1% of genes in any particular array experiment are represented by more than 1 probeset, a few genes had 2-5 probesets and a very few had as many as 15 or 20. In such cases, we combined probeset-specific fold changes and probeset-specific p-values to generate gene level fold-changes and p-values. Briefly, we used the fold changes to convert the individual probeset-specific two-tailed nominal p-values into z-scores that capture the direction of the change:

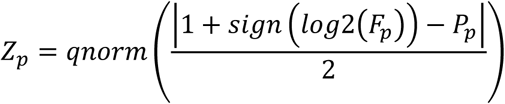

where Z_p_ is the directional probeset-specific z-score, P_p_ is the two-tailed probeset-specific p-value, F_p_ is the probeset-specific fold change, qnorm() is the standard normal inverse CDF, and sign(x) is 1 when x is >= 0 and −1 when x<0. Thus, when F_p_ is >= 1, this yields Z_p_=qnorm(1-P_p_/2) (range is [0,∞)), and when F_p_ <1, this yields the lower tail, Z_p_=qnorm(P_p_/2) (range is (−∞,0]).

A summary gene-specific p-value was calculated as 2 times the upper tail of the standard normal cumulative distribution function assessed at the absolute value of the average of the probeset-specific Z’s:

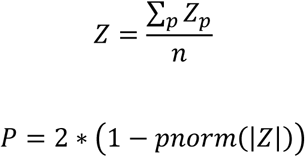

where Z is the average of the probe-specific z-scores, P is the gene specific two-tailed p-value, n is the number of probesets for a gene, and pnorm is the standard normal cumulative distribution function.

The summary gene-specific experiment-specific fold change is calculated by exponentiating the predicted value of log2 fold change from a linear regression of probeset log2 fold changes regressed on probeset z’s, evaluated at the average of the probeset z’s:

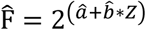

where Fhat is the predicted fold change at Z, ahat is the intercept and bhat is the slope of the linear model of log2(F_p_) modeled as a function of Z_p_.

#### Combination of gene-specific p-values and fold changes across experiments

For each gene, g, in the consensome, we counted the number of experiments, E_g_, where the gene has a nominal p-value of 0.05 or less out of Ng experiments where gene-specific data are not missing. A consensus p-value, P_g_ (CPV) was calculated as the binomial probability of observing E_g_ or more successes out of N_g_ trials, when the true probability of success is 0.05. This provided an estimate of the degree to which the fraction of experiments with alterations exceed what might be expected by chance.

The gene-specific consensus fold change, F_g_, is geometric mean of the experiment-specific fold changes, expressed as max(F_ge_, 1/F_ge_), for the gene of interest. A number of factors determine whether a target will be induced or repressed by manipulation of a given signaling node in any given experiment. These include: node isoform differential expression ^51^; cell cycle stage ^52^; biosample of study ^53^; BSM dose treatment duration; and perturbation type (loss or gain of function). To avoid these opposing alterations canceling each other out at the target transcript level in the meta-analysis, all fold changes were converted to max(F_ge_, 1/F_ge_), such that both inductive and repressive experimental manipulations counted as ‘altered’, allowing us to generate a summary measure of magnitude of perturbation.

The calculation for a hypothetical target is shown below. Genes in the consensome analysis are ranked in ascending order by CPV, with average rank reported in the case of tied CPVs.

| Gene | Experiment | Probeset | Fp | Pp | Zp | Fge | Pge | Eg | Ng | CPV | GMFC |
| --- | --- | --- | --- | --- | --- | --- | --- | --- | --- | --- | --- |
| X | E1 | PX1.1 | 4 | 0.001 | 3.29 | 2.0 | 0.08 | 2 | 4 | 0.014 | 2.83 |
|  |  | PX1.2 | 2 | 0.05 | 1.96 |  |  |  |  |  |  |
|  |  | PX1.3 | 1 | 1.0 | 0 |  |  |  |  |  |  |
|  | E2 | PX2 | 2.5 | 0.1 | 1.645 | 2.5 | 0.1 |  |  |  |  |
|  | E3 | PX3 | 0.333 | 0.02 | -2.33 | 0.333 | 0.02 |  |  |  |  |
|  | E4 | PX4 | 0.333 | 0.02 | -2.33 | 0.333 | 0.02 |  |  |  |  |

In summary, transcriptomic consensome analysis is predicated upon three assumptions: firstly, that borrowing statistical power by binning experiments according to their perturbation of a given signaling node is biologically valid; secondly, that omitting direction of differential expression from the analysis allows for direct interrogation of the strength of the regulatory relationship between a node and a target, independent of the nature of the node perturbation used in an experiment; and thirdly, that ranking targets according to the frequency of their significant differential expression, rather than by fold change, accurately reflects the strength of the regulatory relationship between a given node and its transcriptional targets.

#### Generation of cistromic consensomes

For calculation of ChIP-Seq consensomes, groups of experiments were designated whose IP nodes mapped to a defined node family. These classes were further sorted into meta-analysis classes based on mapping to the same biosample controlled vocabulary used to annotate the transcriptomic datasets. MACS2 peak calls from the ChIP-Atlas resource ^54^ for all nodes in a defined SPP family were averaged and the targets ranked based upon this value.

#### Maintenance and versioning of consensomes

SPP is continually expanding its base of data points by adding newly biocurated datasets to the resource. Accordingly, a quarterly process identifies all node/family and biosample category combinations represented by datasets added in the previous quarter and calculates new versions of the corresponding consensomes. A statement above the scatterplot and contained in the associated spreadsheet identifies the specific combination of pathway node, biosample (physiological system and organ) and species represented by the consensome, the version and date stamp, and the total number of data points, experiments and datasets on which it is based.

### Statistical analysis

Full descriptions of the statistical analyses for each experiment are included in the descriptions of those experiments below and in the Figure Legends.

### Data availability

All SPP datasets and consensomes are freely available on the SPP website under a Creative Commons Attribution 3.0 license, which provides for sharing, adaptation and both non-commercial and commercial re-use, as long as the resource is cited. Data points can be accessed programmatically via RESTful API – full documentation is available at https://beta.signalingpathways.org/docs/.

### Signaling Pathways Project web application

The Signaling Pathways Project knowledgebase is a gene-centric Java Enterprise Edition 6, web-based application around which other gene, mRNA, protein and BSM data from external databases such as NCBI are collected. After undergoing semiautomated processing and biocuration as described above, the data and annotations are stored in SPP’s Oracle 12c database. RESTful web services exposing SPP data, which are served to responsively designed views in the user interface, were created using a Flat UI Toolkit with a combination of JavaScript, D3.JS, AJAX, HTML5, and CSS3. JavaServer Faces and PrimeFaces are the primary technologies behind the user interface. SPP has been optimized for Firefox 24+, Chrome 30+, Safari 5.1.9+, and Internet Explorer 9+, with validations performed in BrowserStack and load testing in LoadUIWeb. XML describing each dataset and experiment is generated and submitted to CrossRef to mint DOIs.

### Bench validation and characterization experiments

#### Validation and characterization of *TPD52L1* in the ER human mammary gland consensome

##### ER-Hs-MG transcriptomic consensome Q-PCR

MCF-7 cells were maintained in DMEM and Ham/F12 Nutrient Mixture (DMEM/F12) supplemented with 8% Fetal Bovine Serum (FBS), Sodium Pyruvate (NaPyr) and non-essential amino acids (NEAA) and passaged every 2-3 days. For experiments, cells were plated in media lacking phenol red with 8% charcoal-stripped FBS (CFS; Gemini). Cells were plated for 48 hours and then treated for 18 hours with 17BE2 (Sigma), or FULV (Tocris). Total RNA was isolated using the Aurum Total RNA Mini-Kit according to the manufacturer’s instructions (Bio-Rad). Total RNA (0.5μg) was reverse-transcribed to cDNA using the iScript cDNA synthesis Kit (Bio-Rad). qPCR was performed using 1.625 μL of Bio-Rad SYBR green supermix, 0.125 μL of a 10 μM dilution of each forward and reverse primer, 0.25 μL of water and 1.25 μL of diluted cDNA for a total reaction volume of 3.25 μL. PCR amplification was carried out using the CFX384 qPCR system. Fold induction was calculated using the 2−ΔΔCt method ^55^, and normalized to 36B4. All data shown is representative of at least three independent experiments. Primer sequences are shown in Supplementary File 9.

##### Subcellular distribution

MCF-7 cells were kept in 5% CD-CS for 48 h prior treatment with 17BE2 10nM for 24 h. A previously published immunofluorescence protocol was followed ^56^. Briefly, cells were fixed in 4% formaldehyde in PEM buffer (80 mM potassium PIPES [pH 6.8], 5 mM EGTA, and 2 mM MgCl2), quenched with 0.1 M ammonium chloride for 10 min, and permeabilized with 0.5% Triton X-100 for 30 min. Cells were incubated at room temperature in 5% Blotto for 1 h, and then specific antibodies were added overnight at 4°C prior to 30 min of secondary antibody (AlexaFluor488 conjugated; Molecular Probes, 1:1000) and DAPI staining. Primary antibody (rabbit polyclonal, Proteintech 14732-1-AP) was diluted at 1:50. A secondary antibody only control showed no appreciable signal (data not shown). Imaging was performed on a GE Healthcare DVLive image restoration deconvolution microscope using an Olympus PlanApo 40x/0.95NA with z-stacks (0.25μm steps covering 12μm) and deconvolved. Images shown are from a maximum intensity projection.

##### Cell proliferation assay

MCF-7 cells (from BCM Tissue Culture Core via ATCC) were plated at 3×10^5^ cells per well of a six well plate in phenol red-free DMEM supplemented with 5% charcoal-stripped FBS. Cells were transfected with 50 nM of a siGENOME SMARTpool targeting human *TPD52L1* (Dharmacon, M-019567-02) or 50 nM of a siGENOME non-targeting pool #2 (Dharmacon, D-001206-14-05) using RNAiMAX (Invitrogen). After two days of knockdown, the cells were split to a 96-well plate in the same media and subsequently treated with (-/+) 10 nM water-soluble 17BE2 (Sigma) for 24 hours. After control or TPD52L1 siRNA transfections and (-/+) 17BE2 treatments, cell viabilities were measured by a CellTiter-Glo® Luminescent assay (Promega). Total RNA was isolated with an RNeasy kit (Qiagen). cDNA was made using 1 μg total RNA and Superscript III reverse transcriptase (Invitrogen) in 20 μl reactions total. To measure the relative mRNA levels, real-time reverse transcription-quantitative PCR (RT-qPCR) was performed in an Applied Biosystems Step One Plus real-time PCR system (Applied Biosystems, Foster City, CA) using 2 μl cDNA diluted 1:10, 900 nM primers, and 0.1 nM Universal Probe designed by the Roche Assay Design Center. Human TPD52L1 primers and probe were forward, 5’- CAACTGTCACAAGCCTCAAGA-3’; reverse, 5’-AGCCTCCTGCCAAGCTCt-3’; Roche probe #73; human β-actin primers and probe were previously described ^57^. Average threshold cycle (Ct) values of human β-actin mRNA were subtracted from corresponding average Ct values of *TPD52L1* mRNA to obtain ΔCt values. Relative mRNA levels were expressed as 2−ΔΔCt compared to the non-targeting siRNA control ^58^. Statistical significance was determined using the Student’s t-Test, and p values < 0.05 were considered significant.

#### Validation and characterization of *MBOAT2* in the AR-Hs-Prostate consensome

##### Cell Culture siRNA Transient Transfections and R1881 Treatments

LNCaP cells (ATCC; Baylor College of Medicine Tissue Culture Core) were plated in 12-well dishes (for gene expression analyses) or 6-well dishes (for cell viability assays) at 1×10^6^ and 2×10^6^, respectively, in charcoal stripped RPMI 1640 media (supplemented with 10% stripped-stripped fetal calf serum, penicillin/streptomycin) and transfected in triplicate with 50 nM of an *MBOAT2* targeting siRNA or a non-targeting siRNA using TransIT-TKO transfection reagent for 5 days. For gene expression analyses, 1 nM R1881 was added to cells on day 4. For cell viability assays, 0.1 nM R1881 was added to cells on day 2. All samples were then harvested on day 5.

##### Gene Expression Analyses by RT-qPCR

On day 5, the RNA from the 12-well plate LNCaP cell samples was harvested using Tri-reagent, following the manufacturer’s instructions. The RNA concentrations were quantitated by Nanodrop (ThermoFisher Nanodrop Lite). 1 µg of each RNA sample was used to make cDNAs by First-Strand cDNA Synthesis using SuperScript II Reverse Transcriptase, following the manufacturer’s protocol. cDNAs were then diluted with 180 µl of DEPC-treated water. To analyze gene expression, 2 µl of cDNAs were used in the RT-qPCR reactions along with Taqman Universal MM II, 200 nM primers (using Roche Diagnostics Universal ProbeLibrary System Assay Design *ACTB*: forward 5’-CCAACCGCGAGAAGATGA-3’, reverse 5’- CCAGAGGCGTACAGGGATAG-3’, probe #64; *MBOAT2*, forward 5’- TCAGACAGCTCTTTGGCTCA-3’, reverse 5’-ACACCCCTGTTAGAAACGTTAGAT-3’, probe #53; *KLK3*, forward 5’-CCTGTCCGTGACGTGGAT-3’, reverse 5’- CAGGGTTGGGAATGCTTCT-3’, probe #75; and *FKBP5*, forward 5’- ACAATGAAGAAAGCCCCACA-3’, reverse 5’-CACCATTCCCCACTCTTTTG-3’, probe #55,), on a StepOnePlus machine (Applied Biosystems). Expression levels of *MBOAT2*, *KLK3*, and *FKBP5* were normalized to *ACTB* and determined by the ΔCt method. PRISM software was used for statistical analyses.

##### Cell Viability Assay

On day 5, the 6-well plate LNCaP cells were briefly trypsinized and collected. Cell viability was then determined using CellTiter-Glo Luminescent Cell Viability Assay, following the manufacturer’s instruction, and a Berthold 96 well plate reading luminometer. PRISM software was used for the statistical analyses.

#### Validation and characterization of *Ppp1r3c* in the GR-Mm-Metabolic consensome

Hepa1c cells were grown in DMEM with 10% fetal bovine serum and penicillin, streptomycin and gentamycin (Life Technologies) and treated with vehicle (ethanol) or 250 nM DEX (Sigma) for 48h. Cells were lysed in TriZOL and total RNA was purified by a PureLink RNA Kit. 250 μg of RNA was reverse transcribed into cDNA using a High Capacity cDNA Reverse Transcription Kit (Life Technologies). Genes were quantified using SYBR Green following the manufacturer’s instructions on an QuantStudio 5 qPCR instrument (Applied Biosystems). Gene expression was normalized to an internal control (*Rplp0*; after evaluating several normalization genes to ensure they were unchanged by treatment). Each experiment was standardized to its own vehicle treatment. Primer sequences used are described in S9 Table.

#### Validation and characterization of *Prkab2* in the ERRs-Mm-Metabolic consensome

##### Animals

All animal protocols were approved by the Institutional Animal Care and Use Committee at City of Hope. The ERRα/Esrra-/- mice have been described and were maintained as a hybrid strain (C57BL/6/SvJ129) ^59, 60^. For baseline comparisons, littermate wild-type and ERRα/Esrra-/- mice were generated from heterozygous breeders to control for strain background. Skeletal muscle (quadriceps) was isolated from 12 week old mice fed wild-type and ERRα/Esrra-/- mice during the daytime (1000 to 1200 h), flash frozen and stored at −80°C until RNA isolation was performed.

##### Cell culture and reagents

C2C12 (ATCC, cell line CRL-1772, Manassas, VA) myoblasts (MB) were cultured in growth media (DMEM (Corning Cellgro, Manassas, VA) containing 10% FBS and differentiated in DMEM containing 2% horse serum (Atlanta Biologicals, Lawrenceville, GA) when MB reached confluence. All experiments were performed in cells below passage number 35. C2C12 myocytes were treated with 5μM XCT790 (Sigma-Aldrich, St. Louis, MO), 0.1 μM DY40 ^61^ or DMSO in growth media or differentiation media prepared with charcoal-stripped serum.

##### Plasmids and transcriptional activity assays

The Prkab2.-2.82.Luc promoter-reporter contains the region of the mouse *Prkab2* gene encompassing −2815 to +27 bp relative to the predicted TSS. The region was amplified from C57B6/J mouse genomic DNA using primers, 5- CTCGGTACCTGAGCACATTAAACCAGTAGTCC-3; 5-GAGAAGCTTTACAAGGCCCGCGACGAGGTAC-3’ (KpnI and HindIII sites in the forward and reverse primers, respectively, denoted in italics) and cloned directly into KpnI/HindIII sites of the pGL3-Basic vector. The entire cloned region was sequenced and confirmed against the corresponding region of the reference Prkab2 gene sequence in NCBI (release 106). The pcDNA3.1-Flag-Esrrg, pSG5-HA-Esrra and pcDNA-myc/his-PGC-1α have been previously described ^62^. Transient transfection in C2C12 myocytes using the calcium phosphate method and the plasmid concentrations used have been described ^63^. Luciferase activity was assayed in MB 48h post-transfection or in day 4 MT after changing confluent cells to 2% HS/DMEM. To assess IGF1 activation, MB were changed to SFM -/+ 10nM recombinant IGF1 one day following transfection and activities were measured after 24 h treatment. Luciferase activity was assayed using Dual-Glo reagents (Promega, Madison, WI) on a Tecan M200 plate reader (Männedorf, Switzerland). Firefly luciferase activity was normalized to that of *Renilla* luciferase, which was expressed downstream of the minimal thymidine kinase promoter from the pRL-TK-*Renilla* plasmid.

##### Quantitative real-time PCR

Real-time PCR was performed to quantify relative transcript levels in RNA collected from skeletal muscle isolated from mice or from day 3 MT using TRIzol reagent (Life Technologies, Carlsbad, CA), as described ^63^. RNA (1 μg) was reverse transcribed in 20 μl reactions using the BioRad iScript cDNA Synthesis Kit (BioRad Laboratories) with 1:1 mixture of oligo-dT and random hexamers for 30 min at 42°C. Resulting cDNA is used in PCR reactions (15 μl) performed in 96-well format in triplicate contained 1X SYBR green reagent (BioRad iQ SYBR Green Supermix), 0.4 μM gene specific primers and 0.5 μl of first strand reaction product (diluted 1:2) as previously described ^63^. Cycling and detection was performed using BioRad IQ5 Real Time PCR system. Experimental transcript levels were normalized to 36B4 (Rplp0) ribosomal RNA analyzed in separate reactions. The following mouse-specific primer sets were used to detect specific gene expression: AMPKα2 (*Prkab2*) forward, 5- ACCATCTCTATGCACTGTCCA -3; reverse, 5-CAGCGTGGTGACATACTTCTT-3; 36B4 (*Rplp0*) forward, 5-ATCCCTGACGCACCGCCGTGA -3; reverse, 5- TGCATCTGCTTGGAGCCCACGTT-3.

##### Statistical analysis

All cell experiments were performed in three independent trials with 3 replicates per trial. Data are presented as mean (± S.E.M.) relative activity or expression normalized to control (empty vector or vehicle treated condition). Differences between mean values for luciferase activities and real-time PCR analysis were analyzed by a one-way ANOVA followed by Fisher’s LSD post test or by unpaired Student’s t test using PRISM software (GraphPad Software, San Diego, CA). A p-value of ≤0.05 was considered significantly different.

#### Validation and characterization of *Mcrip2* in the ERRs-Mm-Metabolic and PPARGC-Mm-Metabolic consensome

##### C2C12 myotube cultures

C2C12 myoblasts (ATCC) were plated at low density in 12-well tissue culture plates in complete DMEM (10%FBS), and switched to differentiation medium (DMEM supplemented with 2% horse serum) when cultures approached confluence. For *Esrra* knockdown experiments, myotubes were infected with adenoviruses expressing Ad-shERRα or vector control at m.o.i of 200 on day 4 of differentiation, and an additional dose of Ad-shERRα or vector control (at m.o.i of 100) together with adenoviruses expressing LacZ (control), *Ppargc1a* or *Ppargc1b* (at m.o.i of 50) on day 6 of differentiation. 24 h later, RNA or DNA were harvested for RT-qPCR gene expression or ChIP analyses, respectively. For expression of the constitutively active *Esrra* (VP16-ERRα), *Esrrb* or *Esrrg*, day 6 myotubes were infected with MOI 50 of adenoviruses expressing LacZ (control), VP16-ERRα, ERRβ or ERRγ ^27, 64^. RNA was harvested 24 h later.

##### Primary brown adipocyte cultures

Pre-adipocytes were isolated from the BAT depot of mice with floxed ERR alleles, as previously described ^65^ and cultured in DMEM supplemented with 20 mM HEPES and 20% FBS prior to differentiation. To induce recombination of floxed ERR loci, pre-adipocytes at 70% confluency were incubated for 16 h with GFP- (control) or CRE-expressing lentiviruses in media containing 4 μg/ml polybrene ^65^. Upon confluency (day 0), cells were switched to DMEM supplemented with 10% FBS, 20 nM insulin, 1 nM triiodothyronine, 0.5 mM IBMX, 2 μg/ml DEX and 0.25 mM indomethacin to differentiate. On day 2 of differentiation, cells were switched to DMEM supplemented with 10% FBS, 20 nM insulin, and 1 nM triiodothyronine. For overexpression assays, mature adipocytes were infected on day 5 of differentiation with adenoviruses expressing Ppargc1a or Gadd45g at an m.o.i of 20. RNA was harvested 24 h later.

##### Q-PCR

Quantitative RT-PCR was performed using the following gene-specific primers: *Rplp0* (36B4), CTGTGCCAGCTCAGAACACTG and TGATCAGCCCGAAGAGAAG; *Mcrip2*, GCCTGTGCAGTATGTGGAGA and GGGTCCACTATGGCAACATT: *Mcrip1*, AAGAGAATGTGCGCTTCATTTA and CTAGGCACCGCTCACCAC; *Pdk4*, GTTCCTTCACACCTTCACCAC and CCTCCTCGGTCAGAAATCTTG; *Tfam*, CAAAGGATGATTCGGCTCAG and AAGCTGAATATATGCCTGCTTTTC. Relative mRNA expression was normalized using *Rplp0* (36B4) as reference gene.

##### ChIP

C2C12 myotubes were crosslinked for 10 min at 37° in 1% formaldehyde in PBS. After quenching, sonication to ∼500 bp fragments, and pre-clearing by treatment with protein A/G sepharose, soluble chromatin was immunoprecipitated with antibodies against Esrra or FLAG. Immunoprecipitated DNA was quantified by real-time PCR, using primers for the *Mcrip2* ERRE (TGAGTACTTGCGGTCCTTGA and ACCTTGGAGAAGGTTGATGG), or primers for a region distal to the *Esrra* promoter that lacks ERREs (negative control; primers described in ^66^). Data shown are the mean and standard deviation of three experimental replicates.

