## Supplementary File 5 for "The Signaling Pathways Project: an integrated ‘omics knowledgebase for mammalian cellular signaling pathways"

**Supplementary File 5. Literature and ChIP-Seq cross-validation of top-ranked transcriptomic consensome targets.**

To compare consensome rankings with canonical node-target relationships, we selected the ten top ranked targets in the following consensomes: ERs-Hs-MG; the androgen receptor in human prostate gland (AR-Hs-Prostate); the glucocorticoid receptor in mouse liver (GR-Mm-Liver); and the peroxisome proliferator-activated receptor (PPAR) family in the mouse metabolic system (PPARs-Mm-Metabolic). We then searched the research literature to identify articles in which these genes had been functionally characterized as targets of these receptors. As shown in S6 Table, 36/40 (90%) of the most highly ranked targets across all four consensomes were validated by evidence in the research literature. Interestingly, of the six transcriptomic consensome-predicted node-target relationships for which no supporting literature evidence was found, all but one were in the 85th percentile or higher of the corresponding ChIP-Seq consensome.

AR-Hs-All: Androgen receptor human all organs; ER-Hs-All: Estrogen receptors human all organs; GR-Hs-All: Glucocorticoid receptor human all organs; PPARs-Mm-Metabolic: Peroxisome proliferator activated receptors-mouse-metabolic organs; CC, percentile ranking for target in the cistromic/ChIP-Seq consensome.

| Consensome | Target | Reference | CC | Consensome | Target | References | CC |
|---|---|---|---|---|---|---|---|
| **AR-Hs-All** | *ZBTB16 (PLZF)* | [1] | 74 | **ERs-Hs-MG** | *SGK1* | [2] | 83 |
| | *UAP1* | [3] | 98 | | *TPD52L1* | - | 99 |
| | *KLK3 (PSA)* | [4] | 99 | | *SIAH2* | [5] | 98 |
| | *MBOAT2* | - | 99 | | *IGFBP4* | [6] | 100 |
| | *C5orf30* | - | 61 | | *PRSS23* | [7] | 92 |
| | *C1orf116 (SARG)* | [8] | 99 | | *HSPB8* | [9] | 99 |
| | *ATAD2* | [10] | 100 | | *IL17RB* | [11] | 92 |
| | *HOMER2* | [12] | 63 | | *EFNA1* | [13] | 84 |
| | *TMPRSS2* | [14] | 90 | | *TFF1* | [15] | 99 |
| | *PTGER4 (EP4)* | [16] | 93 | | *SLC9A3R1* | [17] | 98 |
| **GR-Mm-Liver** | *Lpin1* | [18] | 99 | **PPARs-Mm-** | *Slc25a20* | [19] | 93[rd] (Ppara), 99[th] (Pparg) |

| | | | | | |
|---|---|---|---|---|---|
| | | | **Metabolic** | | |
| *Igfbp1* | 20 | 96 | *Pdk4* | 21 | 96th (Ppar), 92nd (Ppar) |
| *Cdkn1a (p21)* | 22 | 99 | *Retsat* | 23 | 79th (Ppara), 96th (Pparg) |
| *Sult1e1* | 24 | 95 | *Hsdl2* | 25 | 94th (Ppara). 97th (Pparg) |
| *Dusp1* | 26 | 99 | *Acsl1* | 27 | 99th (Ppara), 99th (Pparg) |
| *Dusp14* | - | 94 | *Crat* | 28 | 98th (Ppara), 97th (Pparg) |
| *Crybg1* | - | 85 | *Ech1* | 29 | 99th (Ppara), 99th (Pparg) |
| *Apbb3* | - | 98 | *Acox1* | 30 | 99th (Ppara), 92nd (Pparg) |
| *Atp1b1* | 31 | 94 | *Decr1* | 30 | 99th (Ppara), 99th (Pparg) |
| *Serpina3c* | 32 | 87 | *Eci1* | 33 | 98th (Ppara), 97th (Pparg) |
