## Supplementary File 4 for "The Signaling Pathways Project: an integrated ‘omics knowledgebase for mammalian cellular signaling pathways"

| "ID" | "TYPE" | "DOI" | "FAMILY" | "PHYSIOLOGICALSYSTEM" | "ORGAN" |
|---|---|---|---|---|---|
| 145 | Transcriptomic | 10.1621/L7ryO4nfGT.1 | Abl family kinases (ABL) | ALL | ALL |
| 139 | Transcriptomic | 10.1621/OUaswUDREo.1 | Adrenergic receptors | ALL | ALL |
| 31 | Transcriptomic | 10.1621/xZXYSIb66a.1 | Ahr-like | Metabolic | Liver |
| 76 | Transcriptomic | 10.1621/uvIOTxojSB.1 | Ahr-like | Metabolic | ALL |
| 129 | Transcriptomic | 10.1621/71JfGxZa88.1 | Ahr-like | ALL | ALL |
| 38 | Transcriptomic | 10.1621/CIZ76GPaIZ.1 | AIRE | Immune | Thymus |
| 86 | Transcriptomic | 10.1621/6sFBg4bPMX.1 | AIRE | Immune | ALL |
| 138 | Transcriptomic | 10.1621/WT2Z848B2M.1 | AIRE | ALL | ALL |
| 148 | Transcriptomic | 10.1621/dZO9ARWRrA.1 | ALL | Cardiovascular | Vasculature |
| 149 | Transcriptomic | 10.1621/GdRqIZCpIL.1 | ALL | Female reproductive | Mammary gland |
| 150 | Transcriptomic | 10.1621/gcZVQLLBRq.1 | ALL | Metabolic | Liver |
| 151 | Transcriptomic | 10.1621/wAfROXIkqM.1 | ALL | Female reproductive | Uterus |
| 152 | Transcriptomic | 10.1621/EAyfNha9j6.1 | ALL | Female reproductive | Mammary gland |
| 153 | Transcriptomic | 10.1621/NKcuOFvrlW.1 | ALL | Female reproductive | Uterus |
| 154 | Transcriptomic | 10.1621/Il8wdILO5k.1 | ALL | Immune | Leukocytes |
| 155 | Transcriptomic | 10.1621/V3zF72HO4r.1 | ALL | Metabolic | Liver |
| 156 | Transcriptomic | 10.1621/xqg5au6qGj.1 | ALL | Neurosensory | CNS |
| 157 | Transcriptomic | 10.1621/3HA1NnZXGw.1 | ALL | Skeletal | Bone |
| 158 | Transcriptomic | 10.1621/ZEQwZVv9tq.1 | ALL | Male reproductive | Prostate |
| 159 | Transcriptomic | 10.1621/oCt6pq5zrS.1 | ALL | Metabolic | Skeletal muscle |
| 160 | Transcriptomic | 10.1621/NYokig3Vww.1 | ALL | Cardiovascular | Heart |
| 161 | Transcriptomic | 10.1621/1IXtkxGZZC.1 | ALL | Metabolic | Adipose tissue |
| 162 | Transcriptomic | 10.1621/pdaec2oBuX.1 | ALL | Male reproductive | Testis |
| 163 | Transcriptomic | 10.1621/SZ91Inhg1Z.1 | ALL | Other | Stem cells |
| 164 | Transcriptomic | 10.1621/RJOspi4wWX.1 | ALL | Respiratory | Lung |
| 165 | Transcriptomic | 10.1621/csscxT6esb.1 | ALL | Metabolic | Skeletal muscle |
| 166 | Transcriptomic | 10.1621/s1raevLHrx.1 | ALL | Metabolic | Liver |
| 167 | Transcriptomic | 10.1621/ow1SZEwNWc.1 | ALL | Metabolic | Adipose tissue |
| 168 | Transcriptomic | 10.1621/nUPWxTmEvF.1 | ALL | Cardiovascular | Vasculature |
| 169 | Transcriptomic | 10.1621/Wtgh4V65Je.1 | ALL | Other | Fibroblasts |
| 170 | Transcriptomic | 10.1621/17A5vw3NgQ.1 | ALL | Other | Skin |
| 171 | Transcriptomic | 10.1621/8ZaOGdbiZl.1 | ALL | Respiratory | Lung |
| 172 | Transcriptomic | 10.1621/Fx6IMA2ZhJ.1 | ALL | Gastrointestinal | Colon |
| 173 | Transcriptomic | 10.1621/accyhNC7yH.1 | ALL | Metabolic | Kidney |
| 174 | Transcriptomic | 10.1621/HKSPU9w7DC.1 | ALL | Gastrointestinal | Colon |
| 175 | Transcriptomic | 10.1621/ZjcXrmds56.1 | ALL | Immune | Leukocytes |

| | | | | | |
|---|---|---|---|---|---|
| 176 | Transcriptomic | 10.1621/U3ZhpYxJvB.1 | ALL | Other | Skin |
| 177 | Transcriptomic | 10.1621/9QPq92ImpO.1 | ALL | Blood | HSPCs |
| 178 | Transcriptomic | 10.1621/DV685hObyg.1 | ALL | Immune | Thymus |
| 179 | Transcriptomic | 10.1621/r9Fqyi9rGk.1 | ALL | Female reproductive | Placenta |
| 180 | Transcriptomic | 10.1621/DjjBxnEPOW.1 | ALL | Cardiovascular | ALL |
| 181 | Transcriptomic | 10.1621/mqp99oLOMH.1 | ALL | Female reproductive | ALL |
| 182 | Transcriptomic | 10.1621/L55kOnvHgI.1 | ALL | Metabolic | ALL |
| 183 | Transcriptomic | 10.1621/cysXM63ofu.1 | ALL | Female reproductive | ALL |
| 184 | Transcriptomic | 10.1621/FZ2YeZAJmw.1 | ALL | Female reproductive | ALL |
| 185 | Transcriptomic | 10.1621/KRqIsfPpXg.1 | ALL | Immune | ALL |
| 186 | Transcriptomic | 10.1621/b14bAiuBZT.1 | ALL | Metabolic | ALL |
| 187 | Transcriptomic | 10.1621/UTzRg52LOd.1 | ALL | Neurosensory | ALL |
| 188 | Transcriptomic | 10.1621/99Y1UCn8Y4.1 | ALL | Skeletal | ALL |
| 189 | Transcriptomic | 10.1621/m2wPH4Bz3B.1 | ALL | Male reproductive | ALL |
| 190 | Transcriptomic | 10.1621/pwFsTHQZqC.1 | ALL | Gastrointestinal | ALL |
| 191 | Transcriptomic | 10.1621/AICZOg6UBW.1 | ALL | Cardiovascular | ALL |
| 192 | Transcriptomic | 10.1621/tll8Oo8bna.1 | ALL | Metabolic | ALL |
| 193 | Transcriptomic | 10.1621/MIFFYZwRyz.1 | ALL | Male reproductive | ALL |
| 194 | Transcriptomic | 10.1621/GnLFc7ct8R.1 | ALL | Other | ALL |
| 195 | Transcriptomic | 10.1621/FUMgYbHTiy.1 | ALL | Respiratory | ALL |
| 196 | Transcriptomic | 10.1621/4ygEbjWUaY.1 | ALL | Other | ALL |
| 197 | Transcriptomic | 10.1621/EiKBKRPXNY.1 | ALL | Blood | ALL |
| 198 | Transcriptomic | 10.1621/ApbEp3dJa9.1 | ALL | Respiratory | ALL |
| 199 | Transcriptomic | 10.1621/zSva4CLFsD.1 | ALL | Gastrointestinal | ALL |
| 200 | Transcriptomic | 10.1621/zpjZMt7X4Y.1 | ALL | Immune | ALL |
| 201 | Transcriptomic | 10.1621/DktoUCPBA6.1 | ALL | Blood | ALL |
| 17 | Transcriptomic | 10.1621/lKP3CQi5sd.1 | Androgen receptor | Male reproductive | Prostate |
| 20 | Transcriptomic | 10.1621/n5BZZY6OwF.1 | Androgen receptor | Male reproductive | Testis |
| 59 | Transcriptomic | 10.1621/3I6Tnh3vUd.1 | Androgen receptor | Male reproductive | ALL |
| 63 | Transcriptomic | 10.1621/CHiLQWXWxl.1 | Androgen receptor | Male reproductive | ALL |
| 77 | Transcriptomic | 10.1621/TxVb4ifVEX.1 | Androgen receptor | Metabolic | ALL |
| 91 | Transcriptomic | 10.1621/Xuaz4dyQxT.1 | Androgen receptor | ALL | ALL |
| 115 | Transcriptomic | 10.1621/YjauVBA6hX.1 | Androgen receptor | ALL | ALL |
| 36 | Cistromic | 10.1621/Ew4bEahypc.1 | Androgen receptor | All | All |
| 69 | Cistromic | 10.1621/oNV5SeBSRR.1 | Androgen receptor | All | All |
| 20 | Cistromic | 10.1621/LPfAHbcMFr.1 | C/EBP family | All | All |
| 55 | Cistromic | 10.1621/zJO3GG4s7t.1 | C/EBP family | All | All |

| | | | | | |
|---|---|---|---|---|---|
| 141 | Transcriptomic | 10.1621/A1rw9CjTAk.1 | C-Akt kinases (AKT) | ALL | ALL |
| 33 | Cistromic | 10.1621/vPcOZt9MdC.1 | CBP/p300 | All | All |
| 56 | Cistromic | 10.1621/JromglRlyU.1 | CBP/p300 | All | All |
| 146 | Transcriptomic | 10.1621/17Gwficyu3.1 | Collagen receptor family | ALL | ALL |
| 28 | Cistromic | 10.1621/UxfhTZ64J4.1 | CREB-like factors | All | All |
| 46 | Cistromic | 10.1621/oNMFhK8x3b.1 | CREB-like factors | All | All |
| 88 | Transcriptomic | 10.1621/i1Px53udXG.1 | Cyclin-dependent kinases (CDK) | Female reproductive | ALL |
| 140 | Transcriptomic | 10.1621/gyY3kgh1Es.1 | Cyclin-dependent kinases (CDK) | ALL | ALL |
| 89 | Transcriptomic | 10.1621/KB3MOFA1eA.1 | E2F family | Female reproductive | ALL |
| 143 | Transcriptomic | 10.1621/LLAUQrs5JY.1 | E2F family | ALL | ALL |
| 42 | Cistromic | 10.1621/OB3ZexC1aL.1 | E2F family | All | All |
| 65 | Cistromic | 10.1621/CfvWEuWICo.1 | E2F family | All | All |
| 35 | Transcriptomic | 10.1621/zADqgkIXTN.1 | Epidermal growth factor receptors | Female reproductive | Mammary gland |
| 82 | Transcriptomic | 10.1621/kX52uTHASE.1 | Epidermal growth factor receptors | Female reproductive | ALL |
| 132 | Transcriptomic | 10.1621/nN8rZd1okG.1 | Epidermal growth factor receptors | ALL | ALL |
| 142 | Transcriptomic | 10.1621/JaBE1pYhgf.1 | Epidermal growth factor receptors | ALL | ALL |
| 1 | Transcriptomic | 10.1621/NCKMJxKklE.1 | Estrogen receptors | Female reproductive | Mammary gland |
| 5 | Transcriptomic | 10.1621/UH745ztFIs.1 | Estrogen receptors | Female reproductive | Uterus |
| 11 | Transcriptomic | 10.1621/TyaTomxmQ8.1 | Estrogen receptors | Metabolic | Liver |
| 39 | Transcriptomic | 10.1621/BYRcrphDrO.1 | Estrogen receptors | Female reproductive | ALL |
| 45 | Transcriptomic | 10.1621/cUbphXKOnS.1 | Estrogen receptors | Female reproductive | ALL |
| 47 | Transcriptomic | 10.1621/PX6lA4DQcb.1 | Estrogen receptors | Female reproductive | ALL |
| 52 | Transcriptomic | 10.1621/VtQFikSj4y.1 | Estrogen receptors | Metabolic | ALL |
| 92 | Transcriptomic | 10.1621/aS8hFkyFTe.1 | Estrogen receptors | ALL | ALL |
| 100 | Transcriptomic | 10.1621/quqGhwj8vn.1 | Estrogen receptors | ALL | ALL |
| 102 | Transcriptomic | 10.1621/PJF8NEyazd.1 | Estrogen receptors | ALL | ALL |
| 58 | Cistromic | 10.1621/2BKfA4znUo.1 | Estrogen receptors | All | All |
| 64 | Cistromic | 10.1621/BJ7kkEFKgd.1 | Estrogen receptors | All | All |
| 96 | Transcriptomic | 10.1621/72axf8m391.1 | Estrogen-related receptors | ALL | ALL |
| 111 | Transcriptomic | 10.1621/AMjW8Lrezy.1 | Estrogen-related receptors | ALL | ALL |
| 24 | Cistromic | 10.1621/FR3UZluQd2.1 | Estrogen-related receptors | All | All |
| 30 | Cistromic | 10.1621/Ff7Lo3xZNJ.1 | Estrogen-related receptors | All | All |
| 45 | Cistromic | 10.1621/iFskvhHhDn.1 | Ets-like | All | All |
| 57 | Cistromic | 10.1621/WjJlPOInd2.1 | Ets-like | All | All |
| 14 | Transcriptomic | 10.1621/SmZO1t1TgH.1 | Farnesoid X receptor (FXR) | Metabolic | Liver |
| 55 | Transcriptomic | 10.1621/jeDVjnfZKl.1 | Farnesoid X receptor (FXR) | Metabolic | ALL |
| 110 | Transcriptomic | 10.1621/KJBzdZDiD4.1 | Farnesoid X receptor (FXR) | ALL | ALL |

| | | | | | |
|---|---|---|---|---|---|
| 147 | Transcriptomic | 10.1621/emxBKA5xT7.1 | Fibroblast growth factor receptors | ALL | ALL |
| 37 | Transcriptomic | 10.1621/NzDv9cvgJY.1 | FOXA family | Male reproductive | Prostate |
| 85 | Transcriptomic | 10.1621/ZseuiSq6xE.1 | FOXA family | Male reproductive | ALL |
| 134 | Transcriptomic | 10.1621/jzETFZJh44.1 | FOXA family | ALL | ALL |
| 18 | Cistromic | 10.1621/zUMBX1AanX.1 | FOXA family | All | All |
| 29 | Cistromic | 10.1621/IPik89l2Ux.1 | FOXA family | All | All |
| 53 | Cistromic | 10.1621/8vyVFJI94o.1 | FOXC family | All | All |
| 22 | Cistromic | 10.1621/yXMhK2rrCV.1 | FOXD family | All | All |
| 40 | Cistromic | 10.1621/3IpbhsUOUo.1 | FOXF family | All | All |
| 54 | Cistromic | 10.1621/iBQBrWw3Ft.1 | FOXF family | All | All |
| 47 | Cistromic | 10.1621/BHD95AnIk3.1 | FOXH family | All | All |
| 50 | Cistromic | 10.1621/uWfo918s4U.1 | FOXJ family | All | All |
| 23 | Cistromic | 10.1621/UtyAUmJ2xd.1 | FOXK family | All | All |
| 43 | Cistromic | 10.1621/B98u6ho8WE.1 | FOXK family | All | All |
| 27 | Cistromic | 10.1621/dZ8lzlMKhr.1 | FOXL family | All | All |
| 52 | Cistromic | 10.1621/dGon3b5t5k.1 | FOXM family | All | All |
| 61 | Cistromic | 10.1621/hQbAQkObkl.1 | FOXO family | All | All |
| 67 | Cistromic | 10.1621/G1VUbsoJR7.1 | FOXO family | All | All |
| 32 | Cistromic | 10.1621/WEaOphxlDz.1 | FOXP family | All | All |
| 44 | Cistromic | 10.1621/bnkUfJR17j.1 | FOXP family | All | All |
| 60 | Cistromic | 10.1621/edB3TBW9yZ.1 | FOXQ family | All | All |
| 34 | Cistromic | 10.1621/7RQJp4O72F.1 | FOXR family | All | All |
| 27 | Transcriptomic | 10.1621/PfUD47GTR6.1 | Free fatty acid receptors | Immune | Leukocytes |
| 29 | Transcriptomic | 10.1621/sBgJwXnHX5.1 | Free fatty acid receptors | Metabolic | Adipose tissue |
| 73 | Transcriptomic | 10.1621/nAabU8cSjR.1 | Free fatty acid receptors | Immune | ALL |
| 75 | Transcriptomic | 10.1621/kyH5SN36wG.1 | Free fatty acid receptors | Metabolic | ALL |
| 124 | Transcriptomic | 10.1621/Y1aJckvDIO.1 | Free fatty acid receptors | ALL | ALL |
| 125 | Transcriptomic | 10.1621/bdnZTAbCHG.1 | Free fatty acid receptors | ALL | ALL |
| 3 | Transcriptomic | 10.1621/RUTfCxDO5H.1 | G protein-coupled estrogen receptor | Female reproductive | Mammary gland |
| 6 | Transcriptomic | 10.1621/jixZkdOn1G.1 | G protein-coupled estrogen receptor | Female reproductive | Uterus |
| 41 | Transcriptomic | 10.1621/1QsHjWYjms.1 | G protein-coupled estrogen receptor | Female reproductive | ALL |
| 44 | Transcriptomic | 10.1621/O24kSZBygD.1 | G protein-coupled estrogen receptor | Female reproductive | ALL |
| 94 | Transcriptomic | 10.1621/xbVgAAIvyF.1 | G protein-coupled estrogen receptor | ALL | ALL |
| 101 | Transcriptomic | 10.1621/luVZCpT6tm.1 | G protein-coupled estrogen receptor | ALL | ALL |
| 21 | Transcriptomic | 10.1621/EJOOFVuj1w.1 | Glucocorticoid receptor | Metabolic | Liver |
| 22 | Transcriptomic | 10.1621/OASW2fSKOJ.1 | Glucocorticoid receptor | Neurosensory | CNS |
| 23 | Transcriptomic | 10.1621/NLcmSbOVAO.1 | Glucocorticoid receptor | Respiratory | Lung |

| | | | | | |
|---|---|---|---|---|---|
| 33 | Transcriptomic | 10.1621/ueSYh2o5Df.1 | Glucocorticoid receptor | Metabolic | Liver |
| 64 | Transcriptomic | 10.1621/U4UTCr6hpw.1 | Glucocorticoid receptor | Metabolic | ALL |
| 65 | Transcriptomic | 10.1621/vVANLXQU2V.1 | Glucocorticoid receptor | Neurosensory | ALL |
| 66 | Transcriptomic | 10.1621/mFmzqnTftL.1 | Glucocorticoid receptor | Respiratory | ALL |
| 79 | Transcriptomic | 10.1621/hK9izkTNaO.1 | Glucocorticoid receptor | Metabolic | ALL |
| 117 | Transcriptomic | 10.1621/XJ2jpoVxN6.1 | Glucocorticoid receptor | ALL | ALL |
| 118 | Transcriptomic | 10.1621/QdbZ7Mi2ts.1 | Glucocorticoid receptor | ALL | ALL |
| 130 | Transcriptomic | 10.1621/nPeLunDng7.1 | Glucocorticoid receptor | ALL | ALL |
| 25 | Cistromic | 10.1621/jx6Qirj3WK.1 | Glucocorticoid receptor | All | All |
| 31 | Cistromic | 10.1621/tivIZ9bZss.1 | Glucocorticoid receptor | All | All |
| 137 | Transcriptomic | 10.1621/sxJtBux5x3.1 | Insulin receptor family | ALL | ALL |
| 41 | Cistromic | 10.1621/6JgQ5eArnF.1 | Jun factors | All | All |
| 73 | Cistromic | 10.1621/ZlBGp2rgVb.1 | Jun factors | All | All |
| 49 | Cistromic | 10.1621/ATZq8yYrNn.1 | Lysine acetyltransferases (KAT) | All | All |
| 59 | Cistromic | 10.1621/BciKfFV4iR.1 | Lysine acetyltransferases (KAT) | All | All |
| 34 | Transcriptomic | 10.1621/gitZiqMCYO.1 | Mineralocorticoid receptor | Metabolic | Liver |
| 36 | Transcriptomic | 10.1621/MhCH3hmqt6.1 | Mineralocorticoid receptor | Neurosensory | CNS |
| 69 | Transcriptomic | 10.1621/ZmZnYPI2e5.1 | Mineralocorticoid receptor | Female reproductive | ALL |
| 80 | Transcriptomic | 10.1621/5fEZrsClbG.1 | Mineralocorticoid receptor | Metabolic | ALL |
| 81 | Transcriptomic | 10.1621/fwWQYgJZeZ.1 | Mineralocorticoid receptor | Metabolic | ALL |
| 83 | Transcriptomic | 10.1621/rJMZSL7gXX.1 | Mineralocorticoid receptor | Neurosensory | ALL |
| 116 | Transcriptomic | 10.1621/McszX6ox4h.1 | Mineralocorticoid receptor | ALL | ALL |
| 120 | Transcriptomic | 10.1621/nHAplbKZVf.1 | Mineralocorticoid receptor | ALL | ALL |
| 123 | Transcriptomic | 10.1621/PSpyp4qa3f.1 | Mineralocorticoid receptor | ALL | ALL |
| 51 | Cistromic | 10.1621/UOFOCLy4OH.1 | Myc / Max factors | All | All |
| 63 | Cistromic | 10.1621/XOYzTIUcie.1 | Myc / Max factors | All | All |
| 48 | Cistromic | 10.1621/cqbwcCHQ8m.1 | Myocyte enhancer factor 2 | All | All |
| 62 | Cistromic | 10.1621/UHrsKRsVYZ.1 | Myocyte enhancer factor 2 | All | All |
| 38 | Cistromic | 10.1621/EcECnqRDiC.1 | Myogenic transcription factors | All | All |
| 66 | Cistromic | 10.1621/xchGgbbT5W.1 | Myogenic transcription factors | All | All |
| 21 | Cistromic | 10.1621/PLYeJjtqjA.1 | NCoR-like | All | All |
| 68 | Cistromic | 10.1621/g3yYpnU68s.1 | NCoR-like | All | All |
| 70 | Cistromic | 10.1621/vFCOycZQph.1 | NF-kappaB p50 subunit-like factors | All | All |
| 72 | Cistromic | 10.1621/KNKpZNMnv5.1 | NF-kappaB p50 subunit-like factors | All | All |
| 97 | Transcriptomic | 10.1621/wDuH53Cnco.1 | Nuclear receptor coactivator (NCOA) | ALL | ALL |
| 35 | Cistromic | 10.1621/Tr5qgPqJlJ.1 | Nuclear receptor coactivator (NCOA) | All | All |
| 37 | Cistromic | 10.1621/ybVCQJmVml.1 | Nuclear receptor coactivator (NCOA) | All | All |

| | | | | | |
|---|---|---|---|---|---|
| 4 | Transcriptomic | 10.1621/8eSo9vz4cN.1 | Peroxisome proliferator-activated receptors | Metabolic | Liver |
| 9 | Transcriptomic | 10.1621/lRwZWLJbwQ.1 | Peroxisome proliferator-activated receptors | Immune | Leukocytes |
| 19 | Transcriptomic | 10.1621/ETMvukJZD9.1 | Peroxisome proliferator-activated receptors | Metabolic | Liver |
| 26 | Transcriptomic | 10.1621/Yi6gHbYq3N.1 | Peroxisome proliferator-activated receptors | Metabolic | Adipose tissue |
| 43 | Transcriptomic | 10.1621/ALwOVBZT43.1 | Peroxisome proliferator-activated receptors | Metabolic | ALL |
| 50 | Transcriptomic | 10.1621/jwvjWIixUi.1 | Peroxisome proliferator-activated receptors | Immune | ALL |
| 58 | Transcriptomic | 10.1621/LSlhmqyj78.1 | Peroxisome proliferator-activated receptors | Metabolic | ALL |
| 60 | Transcriptomic | 10.1621/Gl3VWRkh4X.1 | Peroxisome proliferator-activated receptors | Metabolic | ALL |
| 72 | Transcriptomic | 10.1621/UW3ZhzPuDS.1 | Peroxisome proliferator-activated receptors | Cardiovascular | ALL |
| 98 | Transcriptomic | 10.1621/bzZQTlQOpH.1 | Peroxisome proliferator-activated receptors | ALL | ALL |
| 104 | Transcriptomic | 10.1621/ijRA3oG5pe.1 | Peroxisome proliferator-activated receptors | ALL | ALL |
| 107 | Transcriptomic | 10.1621/UBtWXaQHZF.1 | Peroxisome proliferator-activated receptors | ALL | ALL |
| 84 | Transcriptomic | 10.1621/WW5dXhweUe.1 | PPARG coactivator 1 (PPARGC1) | Metabolic | ALL |
| 133 | Transcriptomic | 10.1621/yZ9E585v93.1 | PPARG coactivator 1 (PPARGC1) | ALL | ALL |
| 42 | Transcriptomic | 10.1621/ZOTn3KOley.1 | Progesterone receptor | Female reproductive | ALL |
| 67 | Transcriptomic | 10.1621/27inpJ27Jq.1 | Progesterone receptor | Female reproductive | ALL |
| 90 | Transcriptomic | 10.1621/PeFrQfAMrg.1 | Progesterone receptor | ALL | ALL |
| 119 | Transcriptomic | 10.1621/xBvNn58o4A.1 | Progesterone receptor | ALL | ALL |
| 61 | Transcriptomic | 10.1621/EDZG5FOnwY.1 | Prostaglandin G/H synthases (PTGS) | Metabolic | ALL |
| 114 | Transcriptomic | 10.1621/xTwqmrHf54.1 | Prostaglandin G/H synthases (PTGS) | ALL | ALL |
| 10 | Transcriptomic | 10.1621/PdMLZ24EbL.1 | Retinoic acid receptors | Immune | Leukocytes |
| 51 | Transcriptomic | 10.1621/tIqJWsO24p.1 | Retinoic acid receptors | Immune | ALL |
| 108 | Transcriptomic | 10.1621/Ns2ZZLrczY.1 | Retinoic acid receptors | ALL | ALL |
| 128 | Transcriptomic | 10.1621/SIzL9WMaTt.1 | Retinoic acid receptors | ALL | ALL |
| 8 | Transcriptomic | 10.1621/LwmA2TbPyQ.1 | Retinoic acid-related orphan receptors | Immune | Leukocytes |
| 49 | Transcriptomic | 10.1621/hqfGdWIPTC.1 | Retinoic acid-related orphan receptors | Immune | ALL |
| 106 | Transcriptomic | 10.1621/awrRqEB2be.1 | Retinoic acid-related orphan receptors | ALL | ALL |
| 126 | Transcriptomic | 10.1621/eKFA8s5YRX.1 | Retinoic acid-related orphan receptors | ALL | ALL |
| 113 | Transcriptomic | 10.1621/8b4MfIldkt.1 | Retinoid X receptors | ALL | ALL |
| 26 | Cistromic | 10.1621/ZCcXb1XwZx.1 | SREBP factors | All | All |
| 39 | Cistromic | 10.1621/hZ3XVq6WZL.1 | SREBP factors | All | All |
| 7 | Transcriptomic | 10.1621/b1P9UJkmud.1 | Testicular receptors (NR2C) | Immune | Leukocytes |
| 48 | Transcriptomic | 10.1621/ZHKSwOkTPu.1 | Testicular receptors (NR2C) | Immune | ALL |
| 105 | Transcriptomic | 10.1621/f7u22MPXIl.1 | Testicular receptors (NR2C) | ALL | ALL |
| 127 | Transcriptomic | 10.1621/nAbR1IWv6J.1 | Testicular receptors (NR2C) | ALL | ALL |
| 136 | Transcriptomic | 10.1621/5xvSXPqcEu.1 | Toll-like receptors | ALL | ALL |
| 25 | Transcriptomic | 10.1621/oaotMz6MNy.1 | Transient Receptor Potential channels | Metabolic | Adipose tissue |

| | | | | | |
|---|---|---|---|---|---|
| 28 | Transcriptomic | 10.1621/WKeYxCbLdr.1 | Transient Receptor Potential channels | Immune | Leukocytes |
| 32 | Transcriptomic | 10.1621/fndsdkOvmx.1 | Transient Receptor Potential channels | Metabolic | Liver |
| 70 | Transcriptomic | 10.1621/UZQpjZzvJf.1 | Transient Receptor Potential channels | Female reproductive | ALL |
| 71 | Transcriptomic | 10.1621/PKu3kCpb2Y.1 | Transient Receptor Potential channels | Metabolic | ALL |
| 74 | Transcriptomic | 10.1621/PVlTgL8YVQ.1 | Transient Receptor Potential channels | Immune | ALL |
| 78 | Transcriptomic | 10.1621/XvKDi5xYoa.1 | Transient Receptor Potential channels | Metabolic | ALL |
| 95 | Transcriptomic | 10.1621/Mh4VyhIH6i.1 | Transient Receptor Potential channels | ALL | ALL |
| 121 | Transcriptomic | 10.1621/Y8EUiVRGkQ.1 | Transient Receptor Potential channels | ALL | ALL |
| 122 | Transcriptomic | 10.1621/dDFypkqQQ9.1 | Transient Receptor Potential channels | ALL | ALL |
| 135 | Transcriptomic | 10.1621/hrpg2riwJX.1 | Tumour necrosis factor receptors | ALL | ALL |
| 19 | Cistromic | 10.1621/3drXScdrbt.1 | Two zinc-finger GATA factors | All | All |
| 71 | Cistromic | 10.1621/QL7ubwiUCP.1 | Two zinc-finger GATA factors | All | All |
| 13 | Transcriptomic | 10.1621/zMOR8Gqe5m.1 | Vitamin D receptor | Immune | Leukocytes |
| 54 | Transcriptomic | 10.1621/njWvJFJ96z.1 | Vitamin D receptor | Immune | ALL |
| 109 | Transcriptomic | 10.1621/37wVZ5BUwH.1 | Vitamin D receptor | ALL | ALL |
| 2 | Transcriptomic | 10.1621/JGzKfFMKGi.1 | Xenobiotic receptors | Female reproductive | Mammary gland |
| 12 | Transcriptomic | 10.1621/JT31l2usgv.1 | Xenobiotic receptors | Metabolic | Liver |
| 15 | Transcriptomic | 10.1621/hNkpcfdWNd.1 | Xenobiotic receptors | Neurosensory | CNS |
| 16 | Transcriptomic | 10.1621/ueW68JL3zb.1 | Xenobiotic receptors | Skeletal | Bone |
| 18 | Transcriptomic | 10.1621/DXfiJcBixL.1 | Xenobiotic receptors | Metabolic | Liver |
| 24 | Transcriptomic | 10.1621/itszb4BKPh.1 | Xenobiotic receptors | Metabolic | Liver |
| 30 | Transcriptomic | 10.1621/Hdf1zT4Nkt.1 | Xenobiotic receptors | Female reproductive | Uterus |
| 40 | Transcriptomic | 10.1621/rjw4P7krvE.1 | Xenobiotic receptors | Female reproductive | ALL |
| 46 | Transcriptomic | 10.1621/JQPdBBBtyp.1 | Xenobiotic receptors | Female reproductive | ALL |
| 53 | Transcriptomic | 10.1621/oIomhnhhyY.1 | Xenobiotic receptors | Metabolic | ALL |
| 56 | Transcriptomic | 10.1621/prUUGkBcCk.1 | Xenobiotic receptors | Neurosensory | ALL |
| 57 | Transcriptomic | 10.1621/m857iaQyJG.1 | Xenobiotic receptors | Skeletal | ALL |
| 62 | Transcriptomic | 10.1621/n94e1Irpko.1 | Xenobiotic receptors | Metabolic | ALL |
| 68 | Transcriptomic | 10.1621/fvEP4al2wA.1 | Xenobiotic receptors | Metabolic | ALL |
| 93 | Transcriptomic | 10.1621/S6lNXjjD3J.1 | Xenobiotic receptors | ALL | ALL |
| 99 | Transcriptomic | 10.1621/A91BxIxhKq.1 | Xenobiotic receptors | ALL | ALL |
| 103 | Transcriptomic | 10.1621/ThMZTBHQEJ.1 | Xenobiotic receptors | ALL | ALL |

| "SPECIES" |
| --- |
| Human |
| House Mouse |
| House Mouse |
| House Mouse |
| House Mouse |
| House Mouse |
| House Mouse |
| House Mouse |
| Human |
| Human |
| House Mouse |
| Human |
| House Mouse |
| House Mouse |
| Human |
| Norway Rat |
| House Mouse |
| Human |
| Human |
| House Mouse |
| House Mouse |
| Human |
| House Mouse |
| House Mouse |
| Human |
| Norway Rat |
| Human |
| House Mouse |
| House Mouse |
| House Mouse |
| Human |
| House Mouse |
| Human |
| Norway Rat |
| House Mouse |
| House Mouse |

House Mouse
House Mouse
House Mouse
House Mouse
Human
Human
House Mouse
House Mouse
Norway Rat
Human
Norway Rat
House Mouse
Human
Human
House Mouse
House Mouse
Human
House Mouse
House Mouse
Human
Human
Human
House Mouse
Human
House Mouse
House Mouse
Human
House Mouse
Human
House Mouse
House Mouse
Human
House Mouse
House Mouse
Human
House Mouse
Human

House Mouse
Human
House Mouse
Human
House Mouse
Human
Human
Human
House Mouse
House Mouse
House Mouse
Human
Human
Human
Human
House Mouse
Human
House Mouse
Norway Rat
Human
House Mouse
Norway Rat
Norway Rat
Human
Norway Rat
House Mouse
Human
House Mouse
Human
House Mouse
Human
House Mouse
Human
House Mouse
House Mouse
House Mouse
House Mouse

Human
Human
Human
Human
House Mouse
Human
Human
Human
House Mouse
Human
Human
Human
Human
House Mouse
House Mouse
Human
Human
House Mouse
House Mouse
Human
Human
Human
Human
House Mouse
Human
House Mouse
House Mouse
Human
Human
House Mouse
Human
House Mouse
Human
House Mouse
House Mouse
House Mouse
Human

Norway Rat
House Mouse
House Mouse
Human
Norway Rat
Human
House Mouse
Norway Rat
Human
House Mouse
Human
Human
House Mouse
Human
House Mouse
Norway Rat
House Mouse
House Mouse
Norway Rat
House Mouse
House Mouse
Human
House Mouse
Norway Rat
Human
House Mouse
House Mouse
Human
House Mouse
Human
Human
House Mouse
Human
House Mouse
House Mouse
Human
House Mouse

House Mouse
Human
Norway Rat
House Mouse
House Mouse
Human
Norway Rat
Human
House Mouse
House Mouse
Norway Rat
Human
House Mouse
House Mouse
Human
House Mouse
Human
House Mouse
Norway Rat
Norway Rat
Human
Human
Human
House Mouse
Human
Human
Human
House Mouse
Human
Human
House Mouse
Human
Human
Human
House Mouse
House Mouse
House Mouse

Human
Norway Rat
House Mouse
House Mouse
Human
Norway Rat
Human
House Mouse
Norway Rat
Human
House Mouse
Human
Human
Human
Human
Human
Norway Rat
House Mouse
Human
House Mouse
Human
House Mouse
Human
House Mouse
Norway Rat
House Mouse
Human
House Mouse
Human
Human
Norway Rat
House Mouse
