## Supplementary File 3 for "The Signaling Pathways Project: an integrated ‘omics knowledgebase for mammalian cellular signaling pathways"

**Suppementary File 3. Key elements of the SPP query and reporting interface**. **A.** Ominer query form. **B.** The transcriptomic Regulation Report. The default display for single gene queries is by Category, which can be adjusted to cluster data points by biosample or species. The default display for multi-gene queries is by Target**. C.** The cistromic Regulation Report. IP antigens are identified using case-sensitive AGSs to denote experiments in different species. **D.** Fold Change information windows for transcriptomic (upper) and cistromic (lower) Regulation Reports display essential information on the data point. **E.** The Bioactive Small Molecule window displays the pharmacology of any BSMs used in the experiment**. F.** The Fold Change Detail window places the data point in the context of the wider experiment and dataset, and provides for citation of the dataset. **G.** Consensome user interface. The example shows genomic targets most frequently significantly differentially expressed in response to genetic or pharmacological manipulation of the human insulin receptor in a transcriptomic experiment. Targets are ranked by default by the consensome P value (CPV), which equates to the probability that the observed frequency of differential expression occurs by chance. Target symbols link to a SPP Regulation Report filtered by the consensome category and biosample parameters to show the underlying data points.

A.

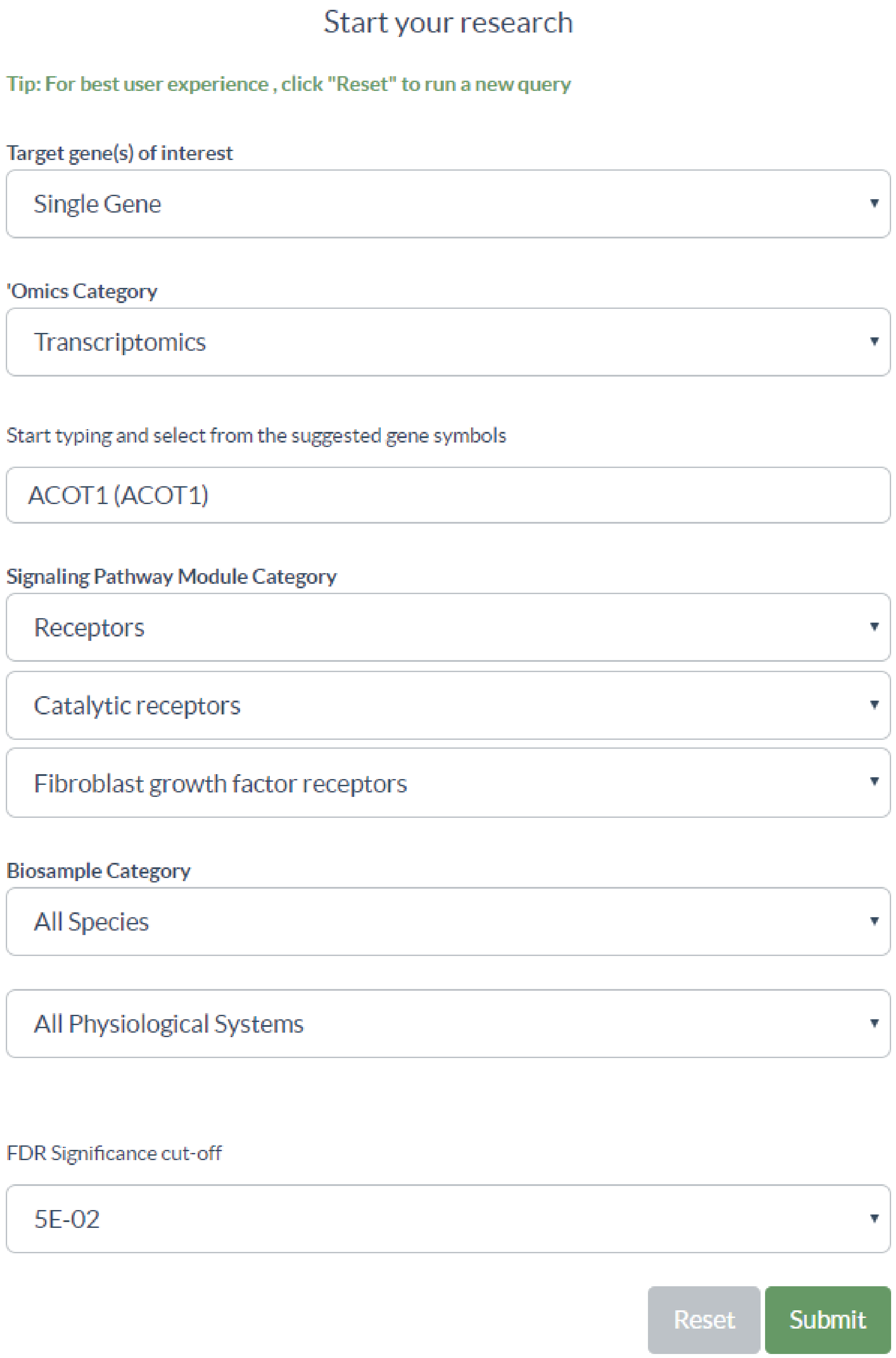
Start your research
Tip: For best user experience , click "Reset" to run a new query
Target gene(s) of interest
Single Gene
'Omics Category
Transcriptomics
Start typing and select from the suggested gene symbols
ACOT1 (ACOT1)
Signaling Pathway Module Category
Receptors
Catalytic receptors
Fibroblast growth factor receptors
Biosample Category
All Species
All Physiological Systems
FDR Significance cut-off
5E-02
Reset
Submit

B.

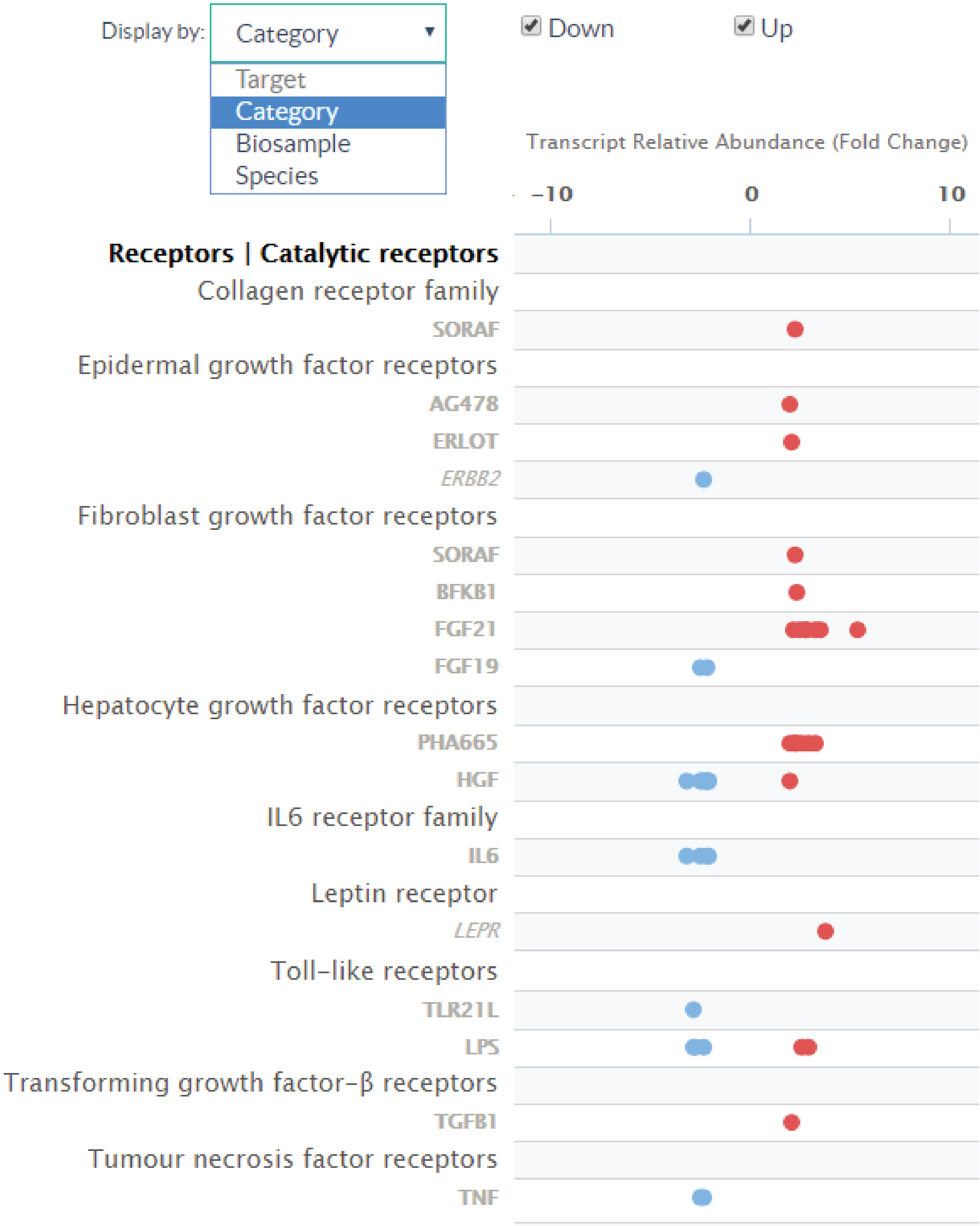
Display by:
Category
Target
Category
Biosample
Species
Down
Up
Transcript Relative Abundance (Fold Change)
−10
0
10
Receptors | Catalytic receptors
Collagen receptor family
SORAF
Epidermal growth factor receptors
AG478
ERLOT
ERBB2
Fibroblast growth factor receptors
SORAF
BFKB1
FGF21
FGF19
Hepatocyte growth factor receptors
PHA665
HGF
IL6 receptor family
IL6
Leptin receptor
LEPR
Toll−like receptors
TLR21L
LPS
Transforming growth factor−β receptors
TGFB1
Tumour necrosis factor receptors
TNF

C.

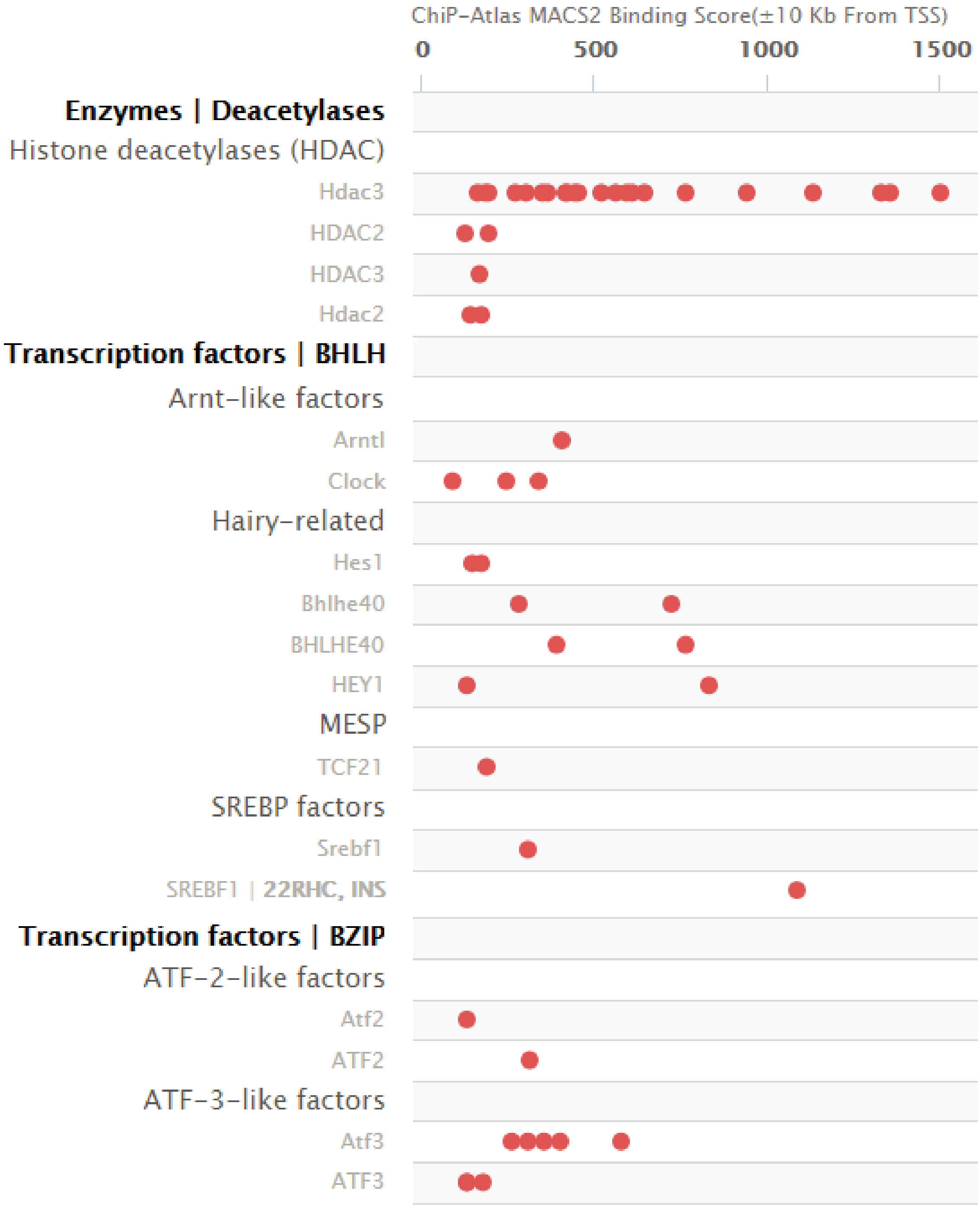
ChiP-Atlas MACS2 Binding Score(±10 Kb From TSS)
0
500
1000
1500
Enzymes | Deacetylases
Histone deacetylases (HDAC)
Hdac3
HDAC2
HDAC3
Hdac2
Transcription factors | BHLH
Arnt-like factors
Arntl
Clock
Hairy-related
Hes1
Bhlhe40
BHLHE40
HEY1
MESP
TCF21
SREBP factors
Srebf1
SREBF1 | 22RHC, INS
Transcription factors | BZIP
ATF-2-like factors
Atf2
ATF2
ATF-3-like factors
Atf3
ATF3

D.

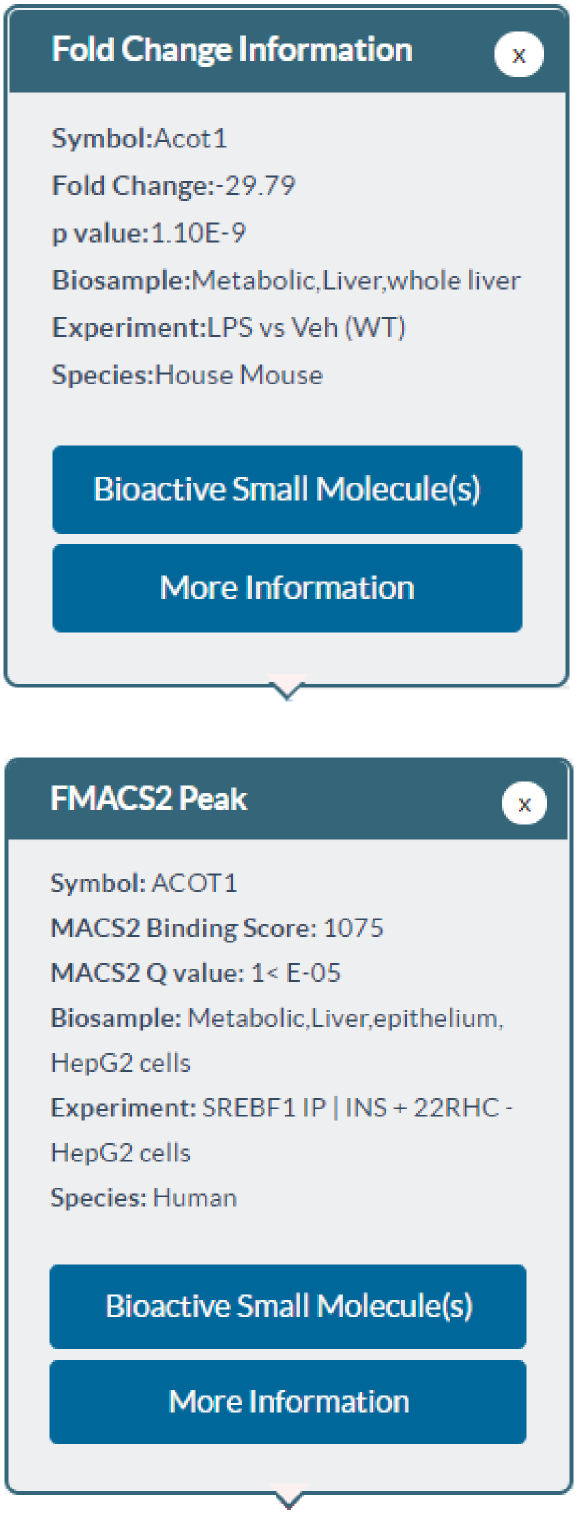
Fold Change Information
x
Symbol:Acot1
Fold Change:-29.79
p value:1.10E-9
Biosample:Metabolic,Liver,whole liver
Experiment:LPS vs Veh (WT)
Species:House Mouse
Bioactive Small Molecule(s)
More Information
FMACS2 Peak
x
Symbol: ACOT1
MACS2 Binding Score: 1075
MACS2 Q value: 1< E-05
Biosample: Metabolic,Liver,epithelium, HepG2 cells
Experiment: SREBF1 IP | INS + 22RHC - HepG2 cells
Species: Human
Bioactive Small Molecule(s)
More Information

E.

### Bioactive Small Molecule(s)

Close

**SPP Symbol:** 22RHC
**BSM Name:** 22(R)-Hydroxycholesterol
**PubChem CID:** 167685
**IUPHAR Guide to Pharmacology ID:** 2742

**SPP Symbol:** INS
**BSM Name:** insulin
**PubChem CID:**
**IUPHAR Guide to Pharmacology ID:** 5012

#### Pharmacology

**Signaling Pathway Module Category:** Receptors
**Class:** Catalytic receptors
**Family:** Insulin receptor family
**Node(s):** INSR

**Signaling Pathway Module Category:** Receptors
**Class:** Nuclear receptors
**Family:** Liver X receptors
**Node(s):** NR1H3,NR1H2

**Signaling Pathway Module Category:** Receptors
**Class:** Nuclear receptors
**Family:** Farnesoid X receptor (FXR)
**Node(s):** NR1H4

F.

### Fold Change Details

Close

**Fold Change Information**

| | |
|---|---|
| **Symbol:** | Acot1 |
| **Fold Change:** | -29.79 |
| **p value:** | 1.10E-9 |

**Experiment Information**

| | |
|---|---|
| **Name:** | LPS vs Veh (WT) |
| **Description:** | Liver was isolated from male BL6/SV129 WT mice treated with 5 µg/gr LPS or vehicle and subsequently fasted for 12 h. |
| **ID:** | 1 |
| **Biosample:** | whole liver |
| **Species:** | House Mouse |

**Dataset Information**

| | |
|---|---|
| **Name:** | Analysis of the acyl-Coenzyme A dehydrogenase, medium chain (Acadm)-dependent and lipopolysaccharide (LPS)-regulated transcriptomes in mouse liver |
| **Description:** | Liver was isolated from male BL6/SV129 Acadm KO and WT mice treated with 5 µg/gr LPS or vehicle and subsequently fasted for 12 h. |
| **DOI:** | 10.1621/kHJ7RMulfF |
| **Download Dataset Citation:** | EN |

G.

### Consensome (beta)

Download Results

**Category**: Receptors
**Class**: Catalytic receptors
**Family**: Insulin receptor family
**Species**: Human
**Physiological System**: All
**Organ**: All

Consensomes are list of genes ranked according to a meta-analysis of their differential expression in publicly archived transcriptomic datasets involving perturbations of a specific signaling pathway in a given biosample category. Consensome are intended as a guide to identifying those genes most consistently impacted by a given pathway in a given tissue context.

Calculated across **588, 339** data points from **28** experiments in **11** datasets.

Show 50 entries

Search:

| Target | Gene Name | Discovery Rate | GMFC | CPV | Percentile |
|---|---|---|---|---|---|
| *PFKFB3* | 6-phosphofructo-2-kinase/fructose-2,6-biphosphatase 3 | 0.913 | 1.383 | 1.09E-25 | 99 |
| *GRPEL1* | GrpE like 1, mitochondrial | 0.913 | 1.452 | 1.09E-25 | 99 |
| *GEM* | GTP binding protein overexpressed in skeletal muscle | 0.87 | 1.963 | 1.46E-23 | 99 |
| *CCNG2* | cyclin G2 | 0.87 | 1.996 | 1.46E-23 | 99 |
| *ARC* | activity regulated cytoskeleton associated protein | 0.87 | 1.619 | 1.46E-23 | 99 |
| *AVPI1* | arginine vasopressin induced 1 | 0.87 | 1.384 | 1.46E-23 | 99 |
| *IER3* | immediate early response 3 | 0.87 | 1.418 | 1.46E-23 | 99 |
| *YRDC* | yrdC N6-threonylcarbamoyltransferase domain containing | 0.87 | 1.682 | 1.46E-23 | 99 |
| *PMAIP1* | phorbol-12-myristate-13-acetate-induced protein 1 | 0.87 | 2.015 | 1.46E-23 | 99 |
| *DUSP2* | dual specificity phosphatase 2 | 0.87 | 1.976 | 1.46E-23 | 99 |
| *AKIRIN1* | akirin 1 | 0.87 | 1.51 | 1.46E-23 | 99 |
