## Supplementary File 2 for "The Signaling Pathways Project: an integrated ‘omics knowledgebase for mammalian cellular signaling pathways"

**Supplementary File 2. SPP UI walk-through**

**Please note: SPP is a work in progress: not all signaling pathway nodes are currently represented in the knowledgebase. Please send suggestions for dataset curation to. Kindly report any bugs (or send feedback) to.**

Annotated datasets can be browsed in the Datasets section of the SPP website.

You can ask questions of the SPP knowledgebase in the Ominer search engine, which allows users to infer regulatory relationships between signaling pathway nodes and their downstream genomic targets.

### A. Single Gene Target Queries.

1) In Ominer, select Single Gene from “Target gene(s) of Interest”

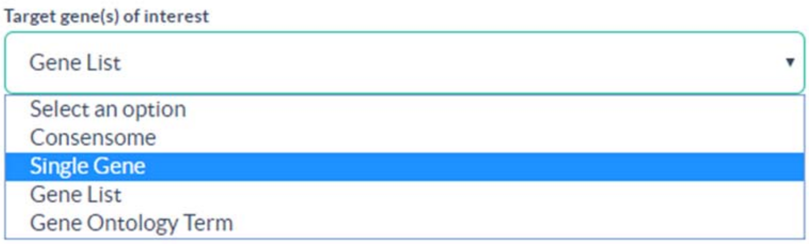


2) Select an ‘omics Category

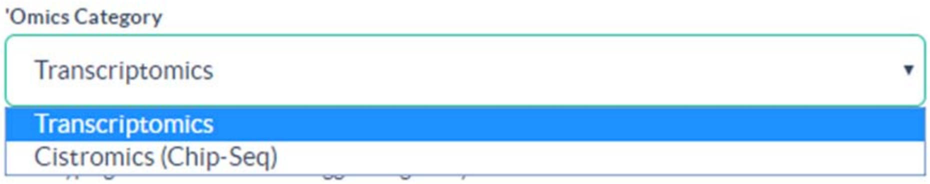


3) Start typing a gene name and select from the drop down suggestions (***you will not be able to submit the query unless you select from the drop down***).

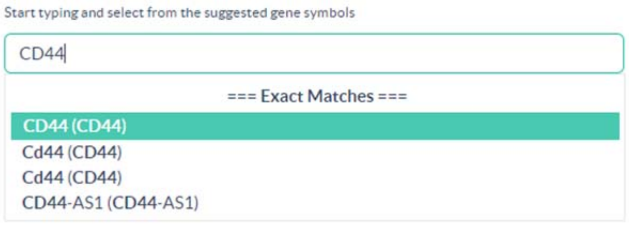


4) Specify a Signaling Pathway Module Category/Class/Family of interest. We recommend the default setting (All) since this returns the most regulatory information for a genomic target.

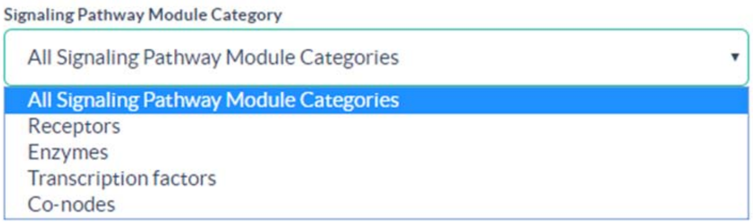


5) Select a Physiological System and/or Organ & species of interest. Again, the default settings are recommended since these will return the most regulatory information for a target of interest.

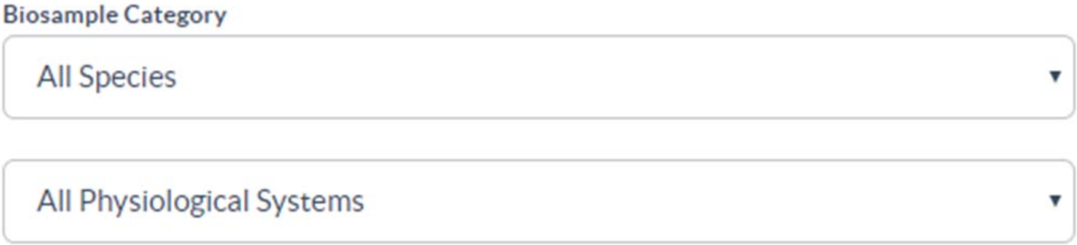


6) Select an FDR Significance Cut-off. We recommend the default (5E-02) unless your query returns more than the maximum of 3,000 data points, in which case the cut-off should be iteratively increased to reduce the number of data points.

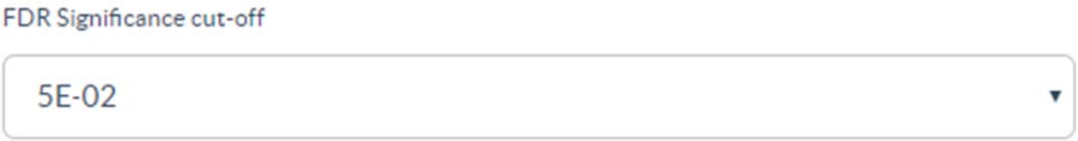


7) Submit your query

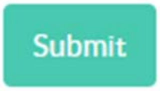


8) SPP will return a **Regulation Report** for your target gene of interest. This is a detailed synopsis of the signaling pathway nodes whose genetic or small molecule manipulation impacts expression of your target of interest (transcriptomic Reports); or the signaling pathway nodes that bind within -/+ 10 kb of the transcription start site of your target gene of interest (cistromic/ChIP-Seq reports). Users can use these Reports to hypothesize candidate signaling pathways regulating their target gene of interest.

The default display is by Target, this can be changed to Pathway Module Category, Biosample or Species using the drop-down in the top left. The layout for transcriptomic and cistromic reports is slightly different so we will briefly summarize each in turn.

**TRANSCRIPTOMIC REPORTS**

The horizontal axis is the transcript relative abundance (fold change).

The vertical axis indicates the Category, Class and Family of the manipulated node. The bottom level labels indicated any genetic (in italics e.g. overexpression, knockdown) or pharmacological (in bold, e.g. BSM administration) node manipulations.

Note: To facilitate database design, some SPP designated families contain only a single node (e.g. androgen receptor, glucocorticoid receptor).

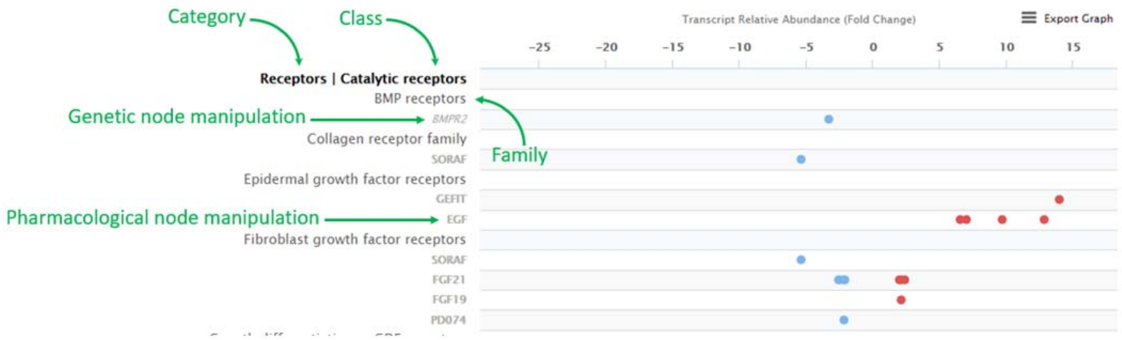


**CISTROMIC/CHIP-SEQ REPORTS**

The horizontal axis is the MACS2 peak height within -/+ 10 kb of the transcription start site of the target gene of interest. MACS2 values are obtained from the ChIP-Atlas resource (Oki et al, BioRxiv 262899).

The vertical axis indicates the Category, Class and Family of the ChIP-Seq antigen. The bottom level labels indicate, in order, the node antigen (normal font); any BSMs in the experiment design (bold) and any genetic manipulations (e.g. overexpression, knockdown) in italics.

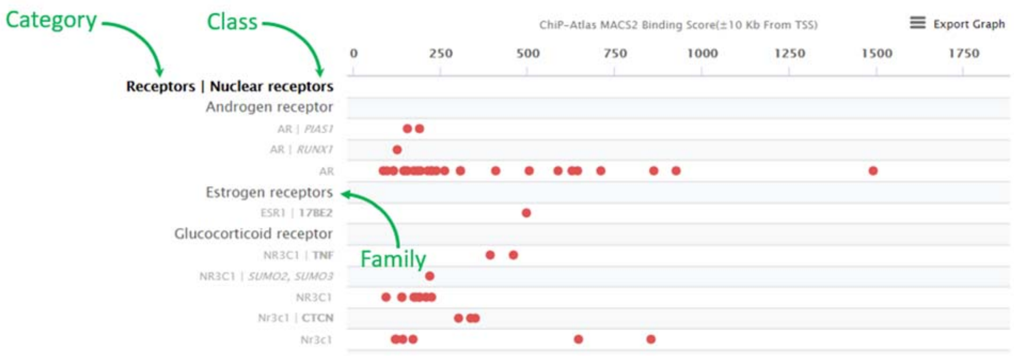


Data points in Regulation Reports are fully interactive. Click on any data point to see its details, and to navigate to a page describing the experiment, as well as a link to the full dataset page.

### B. Gene Ontology term Queries.

1) In Ominer, select “Gene Ontology Term” from “Target Gene(s) of interest”

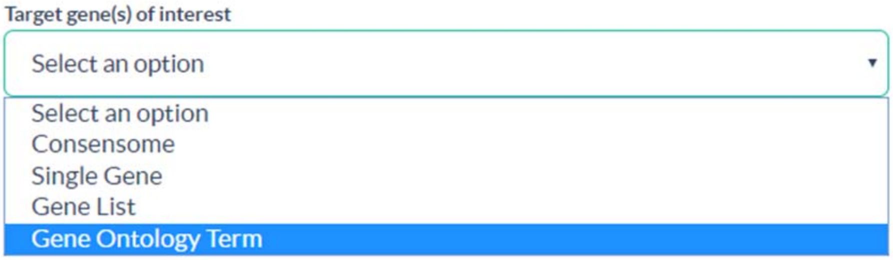


2) Select an ‘omics Category (default is transcriptomic)

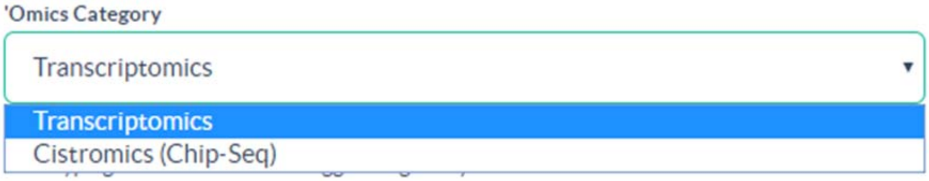


3) Start typing a Gene Ontology term and select from the drop down suggestions (***you will not be able to submit the query unless you select from the drop down***).

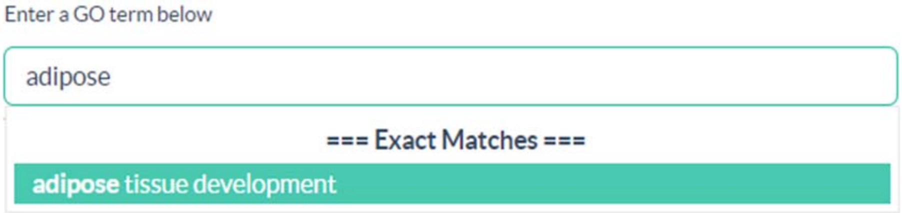


4) To reduce the load on the server, **you <u>must</u> select a Signaling Pathway Module Category (e.g. Enzymes) AND Class (e.g. Kinases) before submitting a GO term query.**

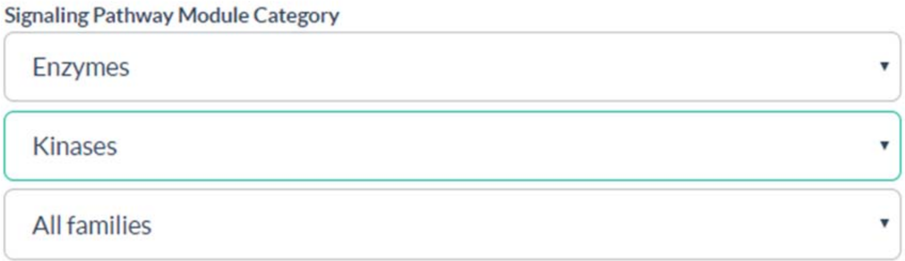


Proceed as in steps 5, 6 and 7 above. The default species is human. As a multi-gene query, the query time may be longer for GO Term queries.

The default display is by Target, this can be changed to Pathway Module Category or Biosample using the drop-down in the top left.

### C. Consensomes.

Consensomes (for consensus ‘omics) are lists of gene targets ranked by frequency of significant differential expression in response to manipulation of nodes in a given pathway node family (transcriptomic consensomes), or the average MACS2 peak scores for all nodes in a given family (cistromic/ChIP-Seq consensomes).

1) In Ominer, select “Consensome” from the “Target gene(s) of interest”

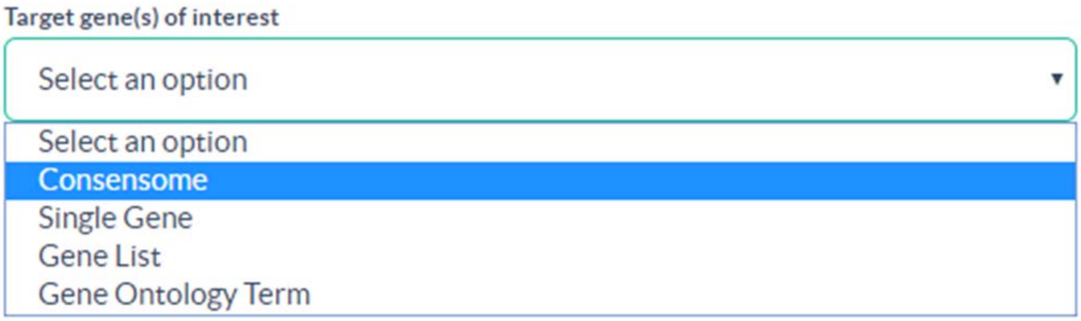


2) Select an ‘omics Category (default is transcriptomic)

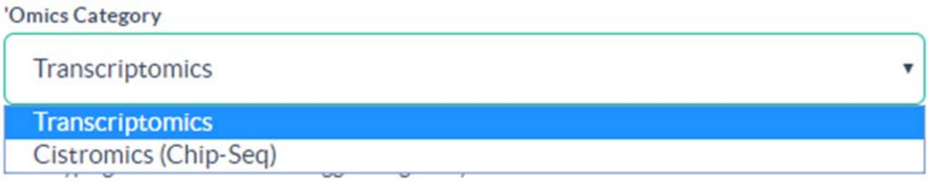


3) Select a Signaling Pathway Module Family,

You must select a Catetgory, Class **and** Family to view a consensome, otherwise the Submit button will not become active.

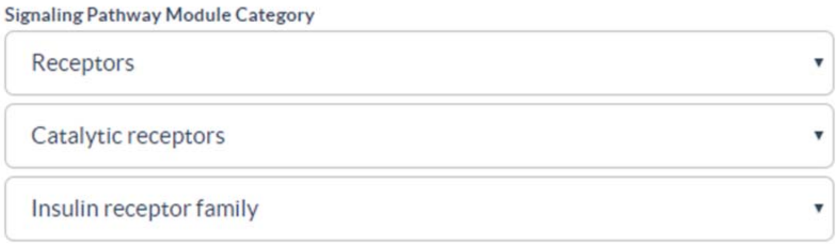


4) Select a Physiological System or Organ. If sufficient curated datasets are available, SPP calculates conensomes for specific Physiological Systems or Organs. If not, only an “All Physiological Systems” consensome will be available.

5) If a consensome is available for your family of interest, the following message will be displayed

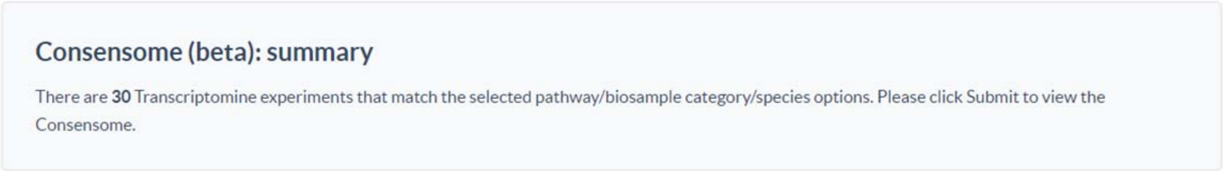


If not, you will see the following message

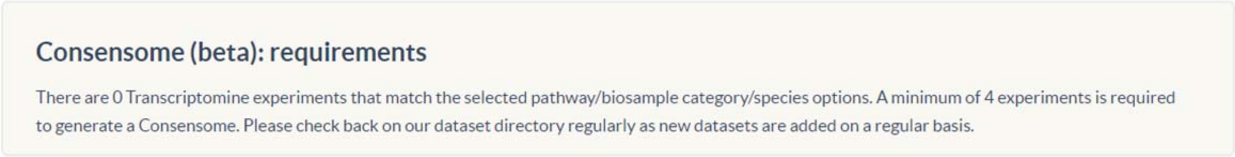


Practical staffing limitations restrict our progress to provide consensomes for all possible node families & omics categories. Some are not currently possible at all due to a shortage of deposited datasets. If you have a suggestion for a node family for which you would like to see a consensome (e.g. Patched receptors), please send an e-mail to. Feedback like this helps us direct our biocuration resources to best serve the research community.

6) Results

**Transcriptomic consensomes** are displayed in ascending order of consensome p-value, which is the probability that the observed frequency of differential expression occurred by chance. Gene target names link to a transcriptomic Regulation Report for that target showing the data points underlying the consensome.

For either consensome type, the top 10% of targets are displayed in the browser. The full conensome can be downloaded by clicking “Download Results”.

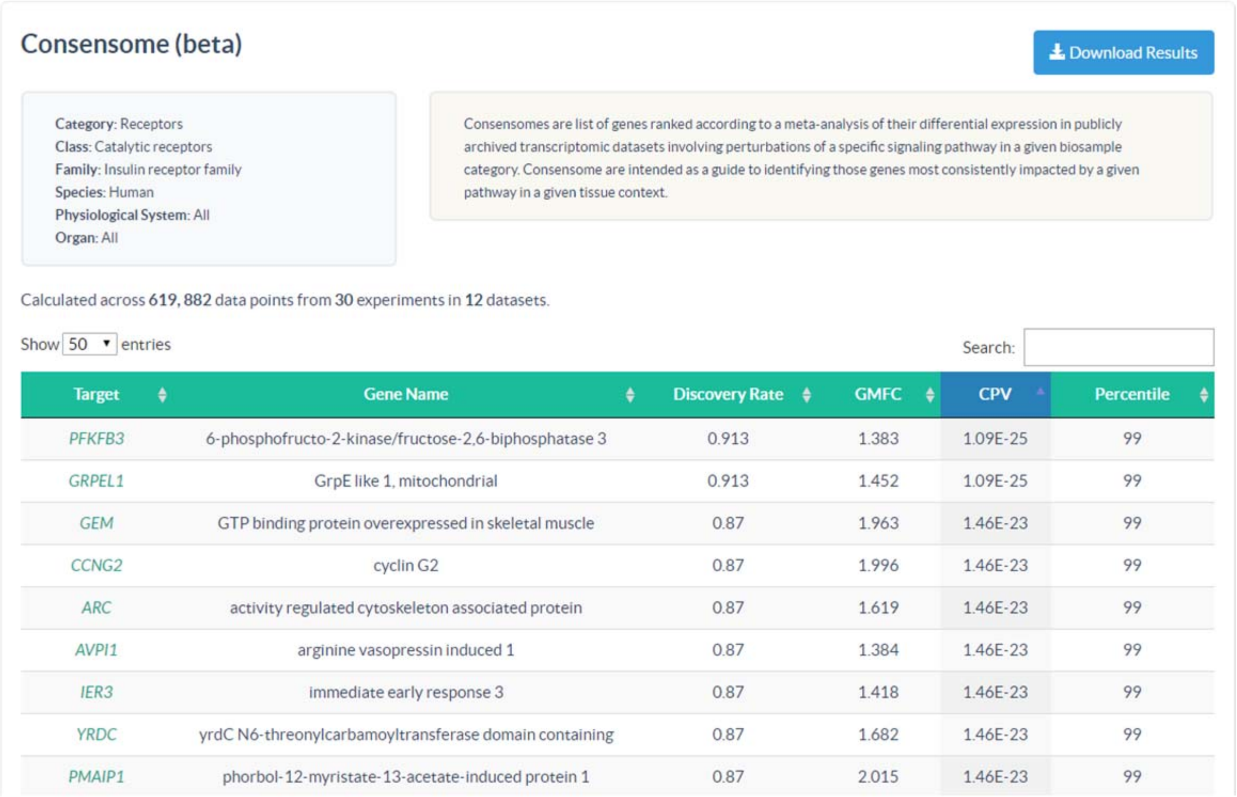

Consensome (beta)

Download Results

Category: Receptors
Class: Catalytic receptors
Family: Insulin receptor family
Species: Human
Physiological System: All
Organ: All

Consensomes are list of genes ranked according to a meta-analysis of their differential expression in publicly archived transcriptomic datasets involving perturbations of a specific signaling pathway in a given biosample category. Consensome are intended as a guide to identifying those genes most consistently impacted by a given pathway in a given tissue context.

Calculated across **619, 882** data points from **30** experiments in **12** datasets.

Show 50 entries | Search:

| Target | Gene Name | Discovery Rate | GMFC | CPV | Percentile |
|---|---|---|---|---|---|
| PFKFB3 | 6-phosphofructo-2-kinase/fructose-2,6-biphosphatase 3 | 0.913 | 1.383 | 1.09E-25 | 99 |
| GRPEL1 | GrpE like 1, mitochondrial | 0.913 | 1.452 | 1.09E-25 | 99 |
| GEM | GTP binding protein overexpressed in skeletal muscle | 0.87 | 1.963 | 1.46E-23 | 99 |
| CCNG2 | cyclin G2 | 0.87 | 1.996 | 1.46E-23 | 99 |
| ARC | activity regulated cytoskeleton associated protein | 0.87 | 1.619 | 1.46E-23 | 99 |
| AVPI1 | arginine vasopressin induced 1 | 0.87 | 1.384 | 1.46E-23 | 99 |
| IER3 | immediate early response 3 | 0.87 | 1.418 | 1.46E-23 | 99 |
| YRDC | yrdC N6-threonylcarbamoyltransferase domain containing | 0.87 | 1.682 | 1.46E-23 | 99 |
| PMAIP1 | phorbol-12-myristate-13-acetate-induced protein 1 | 0.87 | 2.015 | 1.46E-23 | 99 |

Cistromic consensomes are displayed in descending order of MACS2 peak value for all nodes in a given target family. Gene target names link to a cistromic/ChIP-Seq Regulation Report for that target showing the data points underlying the consensome.

#### D. Datasets.

If you simply wish to browse the datasets in SPP, you can do so at https://beta.signalingpathways.org/datasets/index.jsf.

You can filter the datasets by signaling pathway module category, biosample, species or ‘omics type.

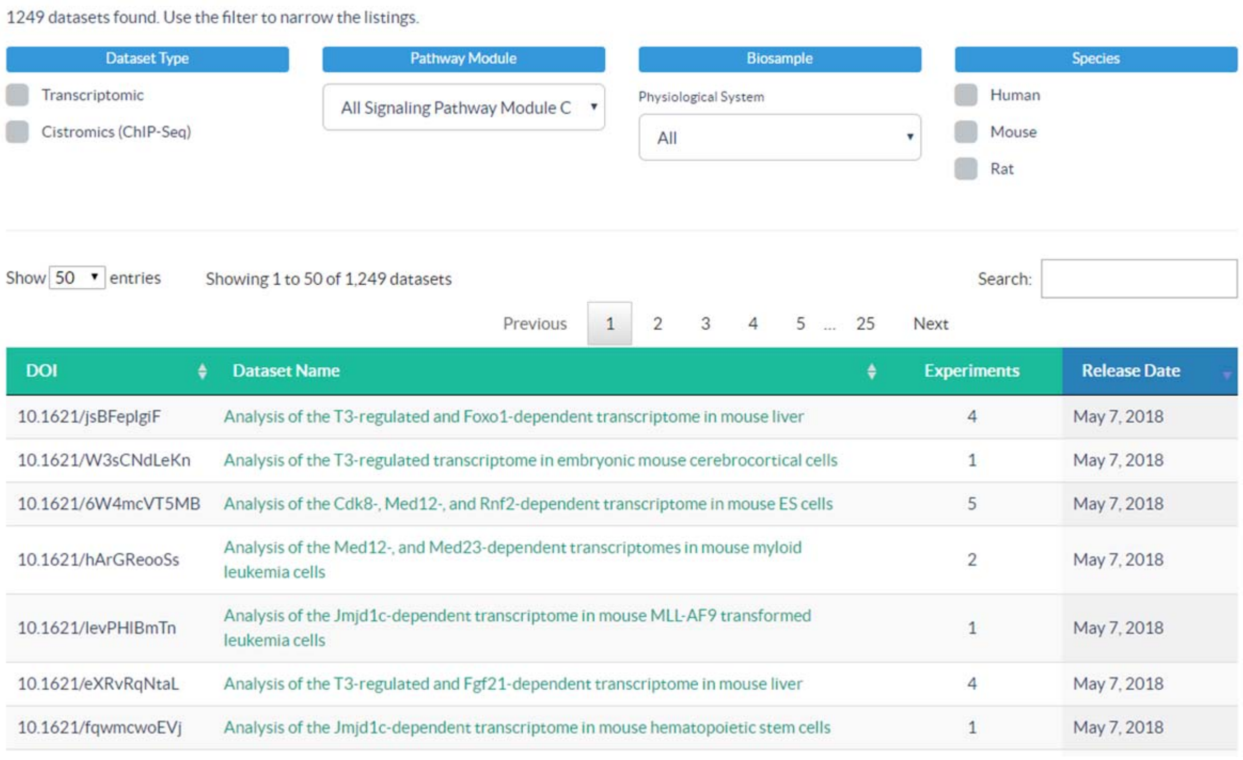


| DOI | Dataset Name | Experiments | Release Date |
|---|---|---|---|
| 10.1621/jsBFeplgiF | Analysis of the T3-regulated and Foxo1-dependent transcriptome in mouse liver | 4 | May 7, 2018 |
| 10.1621/W3sCNdLeKn | Analysis of the T3-regulated transcriptome in embryonic mouse cerebrocortical cells | 1 | May 7, 2018 |
| 10.1621/6W4mcVT5MB | Analysis of the Cdk8-, Med12-, and Rnf2-dependent transcriptome in mouse ES cells | 5 | May 7, 2018 |
| 10.1621/hArGReooSs | Analysis of the Med12-, and Med23-dependent transcriptomes in mouse myloid leukemia cells | 2 | May 7, 2018 |
| 10.1621/levPHIBmTn | Analysis of the Jmjd1c-dependent transcriptome in mouse MLL-AF9 transformed leukemia cells | 1 | May 7, 2018 |
| 10.1621/eXRvRqNtaL | Analysis of the T3-regulated and Fgf21-dependent transcriptome in mouse liver | 4 | May 7, 2018 |
| 10.1621/fqwmcwoEVj | Analysis of the Jmjd1c-dependent transcriptome in mouse hematopoietic stem cells | 1 | May 7, 2018 |

Clicking on the name displays the Dataset page, which displays essential biocurated information, as well as the targets with the highest differential expression values (transcriptomic datasets) or the targets with the highest MACS2 scores (cistromic/ChIP-Seq datasets). Clicking on the target names on the left hand side will run a default single target gene query across the entire SPP knowledgebase.

### Analysis of the T3-regulated and Foxo1-dependent transcriptome in mouse liver

#### Overview

**Dataset Name :** Analysis of the T3-regulated and Foxo1-dependent transcriptome in mouse liver

**Description :** Liver tissue was isolated from 8 week old C57BL/6 male mice 3 d after injection with Foxo1 siRNA or control siRNA followed by injection of 10 µg/kg bw T3 or vehicle.

**Dataset Type :** Transcriptomic

**Release Date :** May 08, 2018

**DOI :** 10.1621/jsBFeplgiF

**Version :** This is Version 1.0 of an annotated derivative of the original dataset, which can be found in GSE68803

**Dataset Citation :** Yen PM, Singh BK, Tripathi M and Ghosh S (2016) Analysis of the T3-regulated and Foxo1-dependent transcriptome in mouse liver, v1.0. SignalingPathway Project Datasets. 10.1621/jsBFeplgiF

**Download Citation :**

**Associated Article :** Singh BK, Sinha RA, Zhou J, Tripathi M, Ohba K, Wang ME, Astapova I, Ghosh S, Hollenberg AN, Gauthier K and Yen PM (2016) Hepatic FOXO1 Target Genes Are Co-regulated by Thyroid Hormone via RICTOR Protein Deacetylation and MTORC2-AKT Protein Inhibition. J. Biol. Chem. **291** 198-214 View Abstract | View PubMed | View Article

Download Dataset

#### Experiments

There are 3 Experiments for this dataset. Up to the 50 up and down p < 0.05 FDR data points are displayed for each. Please download the dataset to view all data points.

1 | T3 vs Con

##### Gene List

Scatter Plot | Conditions

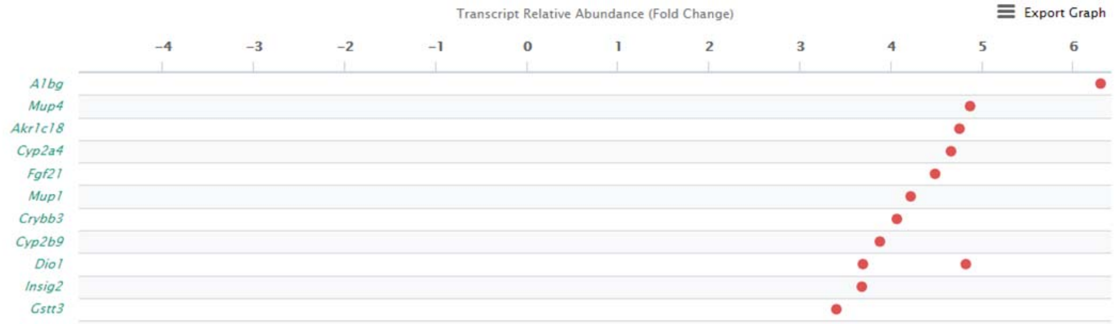
