## Supplementary figures and images for "The Signaling Pathways Project: an integrated ‘omics knowledgebase for mammalian cellular signaling pathways"

### Supplementary File 1

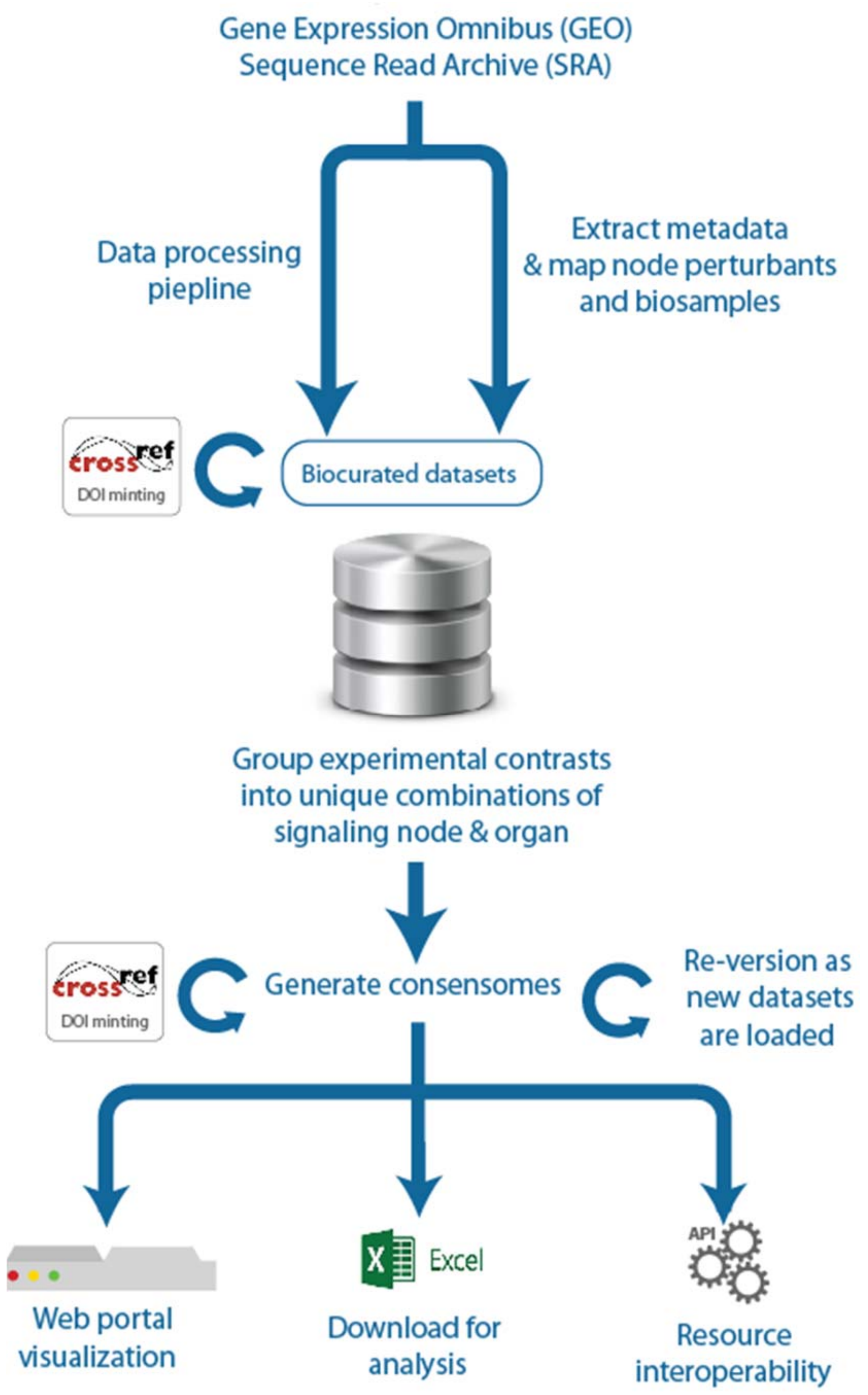


**Supplementary File 1. Flow diagram of SPP biocuration pipeline.**
