## Supplementary File 9 for "The Signaling Pathways Project: an integrated ‘omics knowledgebase for mammalian cellular signaling pathways"

**Supplementary File 9. Q-PCR primers for consensome validation**

| | |
|---|---|
| ERs-Hs-mammary gland transcriptomic consensome | |
| *36B4* | F: GGACATGTTGCTGGCCAATAA |
| | R: GGGCCCGAGACCAGTGTT |
| *ESR1* | F: TGGAGATCTTCGACATGCTG |
| | R: TCCAGAGACTTCAGGGTGCT |
| *GREB1* | F: TGGAAGGCTTGCAGCTCTTGAG |
| | R: GGAAGGGCCGTGTAGCCTTCG |
| *TPD52L1* | F: ACTCGGCATGAACCTGATGA |
| | R: CTGCGTGACTCAGGGTTTCA |
| *CXCL12* | F: ATTCTCAACACTCCAAACTGTGC |
| | R: CTTCAGCCGGGCTACAATCTG |
| *MYBL1* | F: GGCGAAGAGGTCGCGCAGTG |
| | R: TGCCATCGATGCTGGCACTGAA |
| *FHL2* | F: ATCCAAGTGCCAGGAATGCA |
| | R: GTGGCAGATGAAGCAGGTCT |
| *RAB31* | F: CCATCGCTGGAAACAAGTGC |
| | R: AACCACGATGGCACCTATGG |
| *NPY1R* | F: CCACTCTCCTCTTGGTGCTG |
| | R: TGGTTTCACTGGACCTGTACT |
| *IL17RB* | F: GCCCTTCCATGTCTGTGAAT |
| | R: ACTGAAGCTCGCGTTTGTTT |
| *CA12* | F: GTGCTCCTGCTGGTGATCTT |
| | R: TGGACCAGCTATTCTCCCCA |
| *NRIP1* | F: CCAGCCCAAAATGAAGGTGC |

| | |
|---|---|
| | R: GTTTGCTGGGTCTCTGCTCT |
| *MYC* | F: CTACCCTCTCAACGACAG<br>R: TTCTTCCTCATCTTCTTGTTC |
| *PRSS23* | F: AAACCCACTTGGCCTGCATA<br>R: GGATGTAGATGCCCACCTGC |
| *TFF1* | F: TCCCCTGGTGCTTCTATCCTAATAC<br>R: GCAGTCAATCTGTGTTGTGAGCC |
| *STC2* | F: GACCGACGCCACCAACCCAC<br>R: CCCCACATCGCCAGCGTTGA |
| *SLC7A5* | F: GCCTACTTCACCACCCTGTC<br>R: AAGACGGGGATGATCCAGGA |
| *EGR3* | F: CATGTGCGGCGTGGAGTC<br>R: TAGGTCACGGTCTTGTTGCC |
| *PTGES* | F: CAGTATTGCAGGAGCGACCC<br>R: GACGAAGCCCAGGAAAAGGA |
| GR-mouse-liver transcriptomic consensome | |
| *Ppp1r3b* | F: TGAGCCACAGATTGCTGGAG<br>R: TGGATGTCCACAGCCATCAC |
| *Ppp1r3c* | F: TGATCCATGTGCTAGATCCACG<br>R: ACTCTGCGATTTGGCTTCCTG |
| *Pcx* | F: GGCTGCAGCAAGTTTGGTTG<br>R: TAGATGTTAGCTCCGCCCTG |
| *Fgf21* | F: AGCATACCCCATCCCTGACT<br>R: AAGAGACTTTCTGGACTGCG |
| *Rplp0* | F: GAAACTGCTGCCTCACATCCG<br>R: GCTGGCACAGTGACCTCACACG |
