## Supplementary File 8 for "The Signaling Pathways Project: an integrated ‘omics knowledgebase for mammalian cellular signaling pathways"

**Supplementary File 8: Bench validation use case supplementary material**

**Use Case 2. Signaling through GR, ERR family members and insulin receptor regulates targets encoding glycogen synthase phosphatase and kinase holoenzyme regulatory subunits**

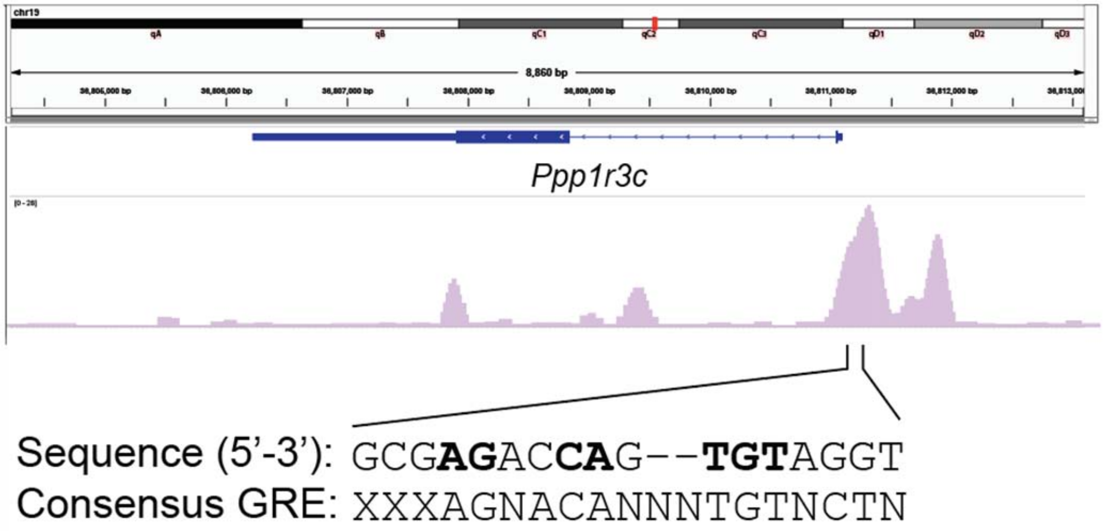


**A.** GR/NR3C1 ChIPseq data from inguinal adipose tissue (iWAT) showing two prominent peaks 5' to the first exon of PPP1R3C (gene is transcribed right to left in this image) and potential GRE within the first GR/NR3C1 peak. Based upon Ominer Regulation Report evidence for binding of GR to the *Ppp1r3c* promoter in mouse liver, we undertook sequence analysis of the murine *Ppp1r3c* promoter and identified two prominent peaks 5' to the first exon of *Ppp1r3c*, the more proximal of which contained a potential glucocorticoid response element (GRE, Fig. 5B; based on the GRE consensus {Reddy, 2009 #131}).

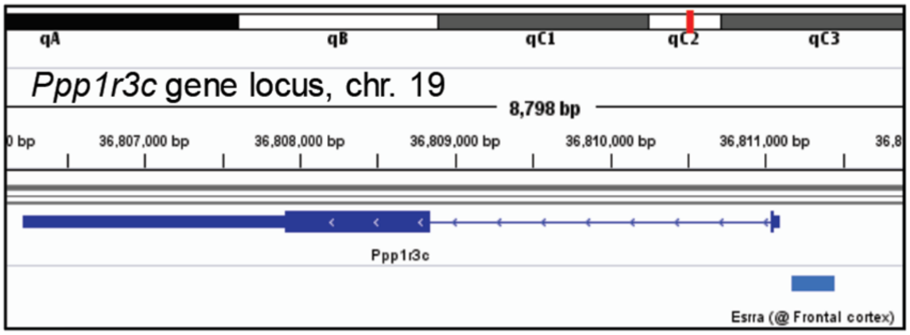

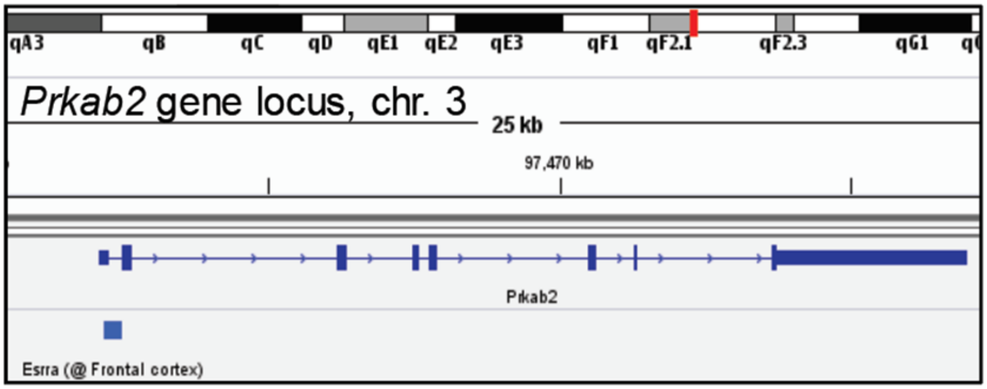


B. Evidence in the SPP cistromic Regulation Reports for *Ppp1r3c* and *Prkab2* and from IVG analysis of additional datasets supports the presence of one or more Esrra binding sites within 10 kb of the Ppp1r3c (upper) and Prkab2 (lower) TSSs.

### Use case 3. The murine ERR, PPARGC and adipose tissue consensomes implicate *Mcrip2* in adipocyte oxidative metabolism

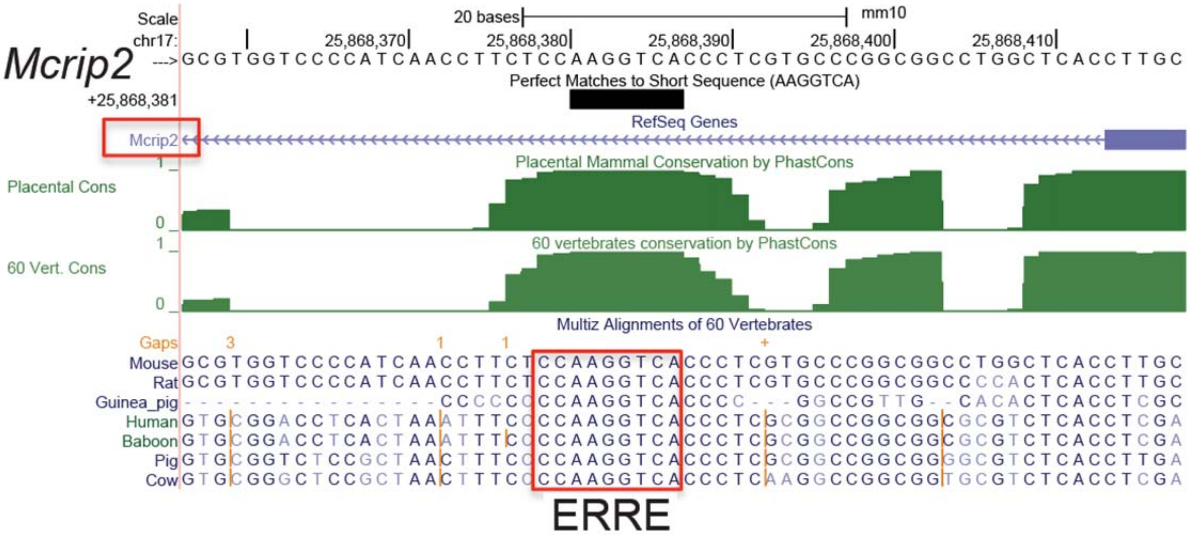


**A. Putative ERREs in Mcrip2 locus.** (source: UCSC genome browser).

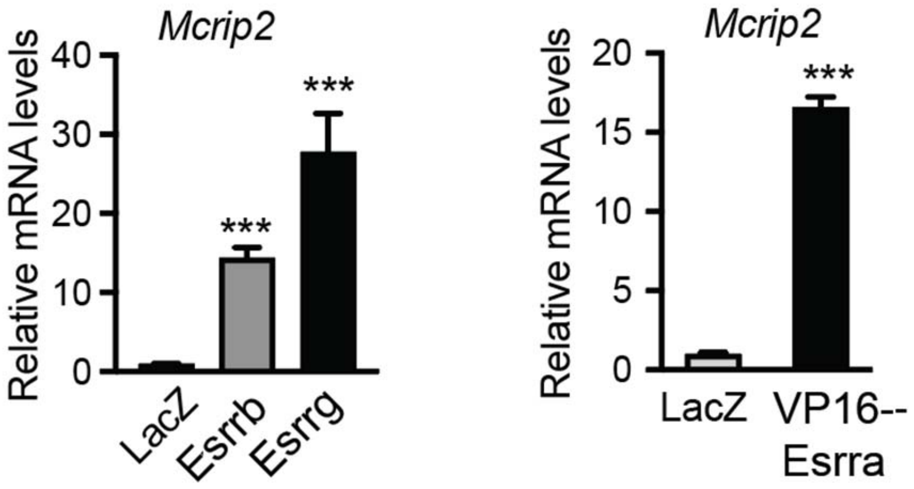


**B.** *Mcrip2* is induced by ERRs. C2C12 myotubes were infected with adenoviral vectors expressing LacZ, VP16-Esrra, Esrrb or Esrrg. RNA and data analyses are described in the Methods section.
