## Supplementary File 6 for "The Signaling Pathways Project: an integrated ‘omics knowledgebase for mammalian cellular signaling pathways"

| Gene name | Mm Approved Symbol | Percentile | CPV | Metabolic enzyme |
|---|---|---|---|---|
| ectonucleoside triphosphate diphosphohydrolase 5 | *Entpd5* | 99.99 | 9.8E-114 | Yes |
| acyl-CoA thioesterase 1 | *Acot1* | 99.99 | 1.6E-111 | Yes |
| Epoxide hydroxylase 1, microsomal xenobiotic | *Ephx1* | 99.98 | 7.3E-107 | Yes |
| Aldehyde dehydrogenase 3 family, member A2 (fatty | *Aldh3a2* | 99.98 | 2.1E-103 | Yes |
| retinol saturase (all trans retinol 13,14 reductase) | *Retsat* | 99.98 | 1.8E-99 | Yes |
| aldehyde oxidase 3 | *Aox3* | 99.97 | 1.7E-94 | Yes |
| Proline dehydrogenase (proline oxidase) | *Prodh* | 99.97 | 2.55E-94 | Yes |
| enoyl coenzyme A hydratase 1, peroxisomal | *Ech1* | 99.96 | 5.12E-91 | Yes |
| glutathione S-transferase, mu 4 | *Gstm4* | 99.96 | 1.37E-90 | Yes |
| vanin 1 | *Vnn1* | 99.96 | 4.18E-89 | |
| Enoyl-Coenzyme A, hydratase/3-hydroxyacyl Coenzy | *Ehhadh* | 99.95 | 4.7E-89 | Yes |
| ATP-binding cassette, sub-family C (CFTR/MRP), men | *Abcc3* | 99.95 | 1.39E-88 | |
| malic enzyme 1, NADP(+)-dependent, cytosolic | *Me1* | 99.94 | 2.2E-88 | Yes |
| dexamethasone-induced transcript | *Dexi* | 99.94 | 2.2E-88 | |
| MAPK regulated corepressor interacting protein 2 | *Mcrip2* | 99.94 | 2.2E-88 | |
| peroxisomal biogenesis factor 11 alpha | *Pex11a* | 99.93 | 4.27E-88 | |
| cytochrome P450, family 2, subfamily b, polypeptide | *Cyp2b10* | 99.93 | 1.25E-87 | |
| acetyl-Coenzyme A acyltransferase 1B | *Acaa1b* | 99.93 | 6.09E-87 | Yes |
| Cystathionine beta-synthase | *Cbs* | 99.92 | 6.48E-87 | Yes |
| pyruvate dehydrogenase kinase 4 | *Pdk4* | 99.92 | 1.11E-86 | Yes |
| perilipin 5 | *Plin5* | 99.91 | 1.18E-86 | |
| Histidine ammonia-lyase (histidase) | *Hal* | 99.91 | 3.56E-86 | Yes |
| glutathione S-transferase, alpha 2 (Yc2) | *Gsta2* | 99.91 | 8.58E-86 | Yes |
| glutathione S-transferase, theta 2 | *Gstt2* | 99.9 | 1.07E-85 | Yes |
| serum amyloid A 4 | *Saa4* | 99.9 | 1.07E-85 | |
| chymotrypsin-like elastase family, member 1 | *Cela1* | 99.89 | 1.58E-84 | |
| Argininosuccinate lyase | *Asl* | 99.89 | 1.54E-83 | Yes |
| nicotinamide N-methyltransferase | *Nnmt* | 99.89 | 3.63E-83 | |
| aldehyde oxidase 1 | *Aox1* | 99.88 | 7.68E-83 | Yes |
| P450 (cytochrome) oxidoreductase | *Por* | 99.88 | 1.01E-82 | |
| solute carrier family 25 (mitochondrial carnitine/acyl | *Slc25a20* | 99.88 | 1.22E-81 | |
| carbonyl reductase 1 | *Cbr1* | 99.87 | 3.32E-81 | Yes |

| | | | | |
|---|---|---|---|---|
| 3-hydroxy-3-methylglutaryl-Coenzyme A synthase 2, | *Hmgcs2* | 99.87 | 1.23E-80 | Yes |
| metallothionein 2 | *Mt2* | 99.87 | 1.23E-80 | |
| G0/G1 switch gene 2 | *G0s2* | 99.86 | 2.03E-80 | |
| choline dehydrogenase | *Chdh* | 99.86 | 3.33E-80 | Yes |
| aquaporin 8 | *Aqp8* | 99.85 | 9.66E-80 | |
| CD36 molecule | *Cd36* | 99.85 | 9.66E-80 | |
| Pyruvate kinase, liver and RBC type | *Pklr* | 99.84 | 1.05E-79 | Yes |
| growth differentiation factor 15 | *Gdf15* | 99.84 | 1.05E-79 | |
| hydroxysteroid dehydrogenase like 2 | *Hsdl2* | 99.84 | 1.05E-79 | Yes |
| glycerol-3-phosphate acyltransferase 3 | *Gpat3* | 99.83 | 2.6E-79 | |
| serine (or cysteine) peptidase inhibitor, clade A (alph | *Serpina7* | 99.83 | 2.6E-79 | |
| glutathione S-transferase, mu 2 | *Gstm2* | 99.83 | 4.23E-79 | Yes |
| transmembrane protein 98 | *Tmem98* | 99.82 | 8.52E-79 | |
| solute carrier organic anion transporter family, mem | *Slco1a4* | 99.82 | 1.44E-78 | |
| arginine vasopressin receptor 1A | *Avpr1a* | 99.81 | 1.45E-78 | |
| Epoxide hydrolase 2, cytoplasmic | *Ephx2* | 99.81 | 6.07E-78 | Yes |
| cytochrome P450, family 1, subfamily a, polypeptide | *Cyp1a2* | 99.81 | 1.01E-77 | |
| glutathione S-transferase, mu 3 | *Gstm3* | 99.8 | 1.14E-77 | Yes |
| cysteine sulfinic acid decarboxylase | *Csad* | 99.8 | 3.21E-77 | Yes |
| coagulation factor VII | *F7* | 99.79 | 3.53E-77 | |
| enoyl-Coenzyme A delta isomerase 2 | *Eci2* | 99.79 | 8.59E-77 | Yes |
| aldo-keto reductase family 1, member C19 | *Akr1c19* | 99.79 | 8.74E-77 | |
| glycosylphosphatidylinositol specific phospholipase [illegible] | *Gpld1* | 99.78 | 2.32E-76 | |
| aldehyde dehydrogenase 1 family member A7 | *Aldh1a7* | 99.78 | 2.69E-76 | Yes |
| Glutamic-oxaloacetic transaminase-1, soluble (EC 2.[illegible] | *Got1* | 99.78 | 4.3E-76 | Yes |
| ST3 beta-galactoside alpha-2,3-sialyltransferase 5 | *St3gal5* | 99.77 | 6.22E-76 | |
| cell death-inducing DFFA-like effector c | *Cidec* | 99.77 | 1.2E-75 | |
| enoyl-Coenzyme A delta isomerase 1 | *Eci1* | 99.76 | 1.96E-75 | Yes |
| ubiquitin specific peptidase 18 | *Usp18* | 99.76 | 1.96E-75 | |
| UDP-glucose dehydrogenase | *Ugdh* | 99.76 | 2.62E-75 | Yes |
| peroxisomal membrane protein 4 | *Pxmp4* | 99.75 | 4.34E-75 | |
| acyl-CoA thioesterase 4 | *Acot4* | 99.75 | 7.17E-75 | Yes |
| solute carrier family 22 (organic cation transporter), | *Slc22a5* | 99.74 | 1.64E-74 | |

| | | | | |
|---|---|---|---|---|
| lysophosphatidylcholine acyltransferase 3 | *Lpcat3* | 99.74 | 8.45E-74 | Yes |
| chemokine (C-C motif) ligand 9 | *Ccl9* | 99.73 | 2.23E-73 | |
| protein S (alpha) | *Pros1* | 99.73 | 2.23E-73 | |
| glutathione S-transferase, alpha 4 | *Gsta4* | 99.73 | 2.36E-73 | Yes |
| cellular repressor of E1A-stimulated genes 1 | *Creg1* | 99.72 | 3.14E-73 | |
| cAMP responsive element binding protein 3-like 3 | *Creb3l3* | 99.72 | 5.14E-73 | |
| neuritin 1 | *Nrn1* | 99.72 | 8.43E-73 | |
| early growth response 1 | *Egr1* | 99.71 | 9.47E-73 | |
| carbonic anhydrase 14 | *Car14* | 99.71 | 1.84E-72 | |
| protein phosphatase 1, regulatory subunit 3C | *Ppp1r3c* | 99.71 | 2.4E-72 | |
| transmembrane protein 97 | *Tmem97* | 99.7 | 2.97E-72 | |
| androgen dependent TFPI regulating protein | *Adtrp* | 99.69 | 3.09E-72 | |
| PERP, TP53 apoptosis effector | *Perp* | 99.69 | 4.32E-72 | |
| inhibin beta-C | *Inhbc* | 99.69 | 4.77E-72 | |
| solute carrier family 46, member 3 | *Slc46a3* | 99.68 | 7.04E-72 | |
| acyl-CoA synthetase long-chain family member 1 | *Acsl1* | 99.68 | 7.65E-72 | Yes |
| RIKEN cDNA 4931406C07 gene | *4931406C07Rik* | 99.67 | 3.9E-71 | |
| transcription elongation factor A (SII), 3 | *Tcea3* | 99.67 | 3.9E-71 | |
| phospholysine phosphohistidine inorganic pyrophos[illegible] | *Lhpp* | 99.67 | 5.87E-71 | Yes |
| 2,4-dienoyl-CoA reductase 1 | *Decr1* | 99.66 | 6.23E-71 | Yes |
| solute carrier family 2 (facilitated glucose transporte[illegible] | *Slc2a2* | 99.66 | 6.23E-71 | |
| cytochrome P450, family 2. subfamily c, polypeptide [illegible] | *Cyp2c37* | 99.66 | 6.38E-71 | |
| carboxypeptidase N, polypeptide 2 | *Cpn2* | 99.65 | 9.5E-71 | |
| phospholipase A2 group XIIA | *Pla2g12a* | 99.65 | 3.17E-70 | Yes |
| hairy and enhancer of split 6 | *Hes6* | 99.64 | 5.05E-70 | |
| pipecolic acid oxidase | *Pipox* | 99.64 | 5.05E-70 | |
| perilipin 2 | *Plin2* | 99.64 | 8.02E-70 | |
| ATP-binding cassette, sub-family A (ABC1), member [illegible] | *Abca8a* | 99.63 | 1.26E-69 | |
| ATP binding cassette subfamily G member 5 | *Abcg5* | 99.63 | 1.26E-69 | |
| cytochrome P450, family 2, subfamily c, polypeptide [illegible] | *Cyp2c54* | 99.62 | 1.53E-69 | |
| GrpE-like 2, mitochondrial | *Grpel2* | 99.62 | 1.62E-69 | |
| carboxylesterase 1E | *Ces1e* | 99.62 | 2.02E-69 | Yes |
| fat storage-inducing transmembrane protein 2 | *Fitm2* | 99.61 | 4.98E-69 | |
| coagulation factor XI | *F11* | 99.61 | 5.16E-69 | |

| | | | | |
|---|---|---|---|---|
| solute carrier family 7 (cationic amino acid transport | *Slc7a2* | 99.61 | 6.45E-69 | |
| cytochrome P450, family 4, subfamily f, polypeptide | *Cyp4f14* | 99.6 | 7.9E-69 | |
| lectin, galactoside-binding, soluble, 3 binding proteir | *Lgals3bp* | 99.6 | 9.23E-69 | |
| Hydroxy-delta-5-steroid dehydrogenase, 3 beta- and | *Hsd3b2* | 99.59 | 1.02E-68 | Yes |
| glutathione S-transferase pi 3 | *Gstp3* | 99.59 | 1.03E-68 | |
| kallikrein B, plasma 1 | *Klkb1* | 99.59 | 1.27E-68 | |
| glucosamine-6-phosphate deaminase 1 | *Gnpda1* | 99.58 | 1.65E-68 | Yes |
| cytochrome P450, family 4, subfamily a, polypeptide | *Cyp4a31* | 99.58 | 4.05E-68 | |
| amidohydrolase domain containing 1 | *Amdhd1* | 99.57 | 5.81E-68 | Yes |
| protease, serine 8 (prostasin) | *Prss8* | 99.57 | 5.81E-68 | |
| retinol dehydrogenase 16 | *Rdh16* | 99.56 | 8.1E-68 | Yes |
| tubulin folding cofactor E-like | *Tbcel* | 99.56 | 8.1E-68 | |
| serum amyloid P-component | *Apcs* | 99.56 | 1.27E-67 | |
| abhydrolase domain containing 6 | *Abhd6* | 99.56 | 1.34E-67 | Yes |
| out at first homolog | *Oaf* | 99.55 | 1.68E-67 | |
| flavin containing monooxygenase 5 | *Fmo5* | 99.55 | 1.72E-67 | |
| Sulfite oxidase | *Suox* | 99.54 | 2.12E-67 | Yes |
| oxidative stress induced growth inhibitor 1 | *Osgin1* | 99.54 | 3.11E-67 | |
| progestin and adipoQ receptor family member IX | *Paqr9* | 99.54 | 3.11E-67 | |
| Sarcosine dehydrogenase | *Sardh* | 99.53 | 6.42E-67 | Yes |
| solute carrier family 41, member 2 | *Slc41a2* | 99.53 | 8.38E-67 | |
| UDP glucuronosyltransferase 2 family, polypeptide B | *Ugt2b35* | 99.52 | 9.25E-67 | |
| cytochrome P450, family 4, subfamily a, polypeptide | *Cyp4a14* | 99.52 | 1E-66 | |
| peptidoglycan recognition protein 2 | *Pglyrp2* | 99.52 | 1.29E-66 | |
| acyl-CoA synthetase long-chain family member 5 | *Acsl5* | 99.51 | 1.32E-66 | Yes |
| UDP-glucose pyrophosphorylase 2 | *Ugp2* | 99.51 | 1.57E-66 | Yes |
| homocysteine-inducible, endoplasmic reticulum stre | *Herpud1* | 99.51 | 2.04E-66 | |
| abhydrolase domain containing 14b | *Abhd14b* | 99.5 | 2.69E-66 | Yes |
| biliverdin reductase B (flavin reductase (NADPH)) | *Blvrb* | 99.5 | 2.69E-66 | Yes |
| RIKEN cDNA 2010003K11 gene | *2010003K11Rik* | 99.49 | 4.37E-66 | |
| carboxylesterase 1G | *Ces1g* | 99.49 | 5.06E-66 | Yes |
| cytochrome P450, family 39, subfamily a, polypeptid | *Cyp39a1* | 99.49 | 6.66E-66 | |
| cytochrome P450, family 4, subfamily a, polypeptide | *Cyp4a12a* | 99.48 | 6.96E-66 | |
| tubulin, beta 4B class IVB | *Tubb4b* | 99.48 | 9.06E-66 | |

| | | | | |
|---|---|---|---|---|
| Dimethylglycine dehydrogenase | *Dmgdh* | 99.46 | 1.22E-65 | Yes |
| monoglyceride lipase | *Mgll* | 99.46 | 1.22E-65 | Yes |
| Guanidinoacetate methyltransferase | *Gamt* | 99.46 | 1.22E-65 | Yes |
| Lipin 2 | *Lpin2* | 99.46 | 1.22E-65 | Yes |
| RAB9, member RAS oncogene family | *Rab9* | 99.46 | 1.22E-65 | |
| sulfiredoxin 1 homolog (S. cerevisiae) | *Srxn1* | 99.46 | 1.22E-65 | |
| nuclear factor, interleukin 3, regulated | *Nfil3* | 99.45 | 2.94E-65 | |
| Catechol-O-methyltransferase | *Comt* | 99.44 | 5.27E-65 | Yes |
| cytochrome P450, family 2, subfamily c, polypeptide | *Cyp2c70* | 99.44 | 5.27E-65 | |
| phosphatidylinositol glycan anchor biosynthesis, clas | *Pigp* | 99.44 | 5.27E-65 | |
| RDH16 family member 2 | *Rdh16f2* | 99.44 | 7.11E-65 | |
| pleckstrin homology domain containing, family F (wi | *Plekhf1* | 99.43 | 8.91E-65 | |
| Acyl-Coenzyme A dehydrogenase, C-4 to C-12 straigh | *Acadm* | 99.43 | 9.52E-65 | Yes |
| angiotensinogen (serpin peptidase inhibitor, clade A, | *Agt* | 99.42 | 1.47E-64 | |
| hepatocyte growth factor activator | *Hgfac* | 99.42 | 1.47E-64 | |
| orosomucoid 2 | *Orm2* | 99.41 | 2.27E-64 | |
| retinol binding protein 1, cellular | *Rbp1* | 99.41 | 2.27E-64 | |
| transmembrane protein 37 | *Tmem37* | 99.41 | 2.65E-64 | |
| ribonuclease, RNase A family 4 | *Rnase4* | 99.4 | 3.49E-64 | |
| 2,4-dienoyl-CoA reductase 2 | *Decr2* | 99.4 | 4.14E-64 | Yes |
| angiogenin, ribonuclease, RNase A family, 5 | *Ang* | 99.4 | 4.14E-64 | |
| ATP-binding cassette, sub-family C (CFTR/MRP), men | *Abcc4* | 99.39 | 4.91E-64 | |
| complement component 8, alpha polypeptide | *C8a* | 99.39 | 6.44E-64 | |
| solute carrier family 37 (glucose-6-phosphate transp | *Slc37a4* | 99.39 | 7.36E-64 | |
| serine dehydratase | *Sds* | 99.38 | 8.2E-64 | Yes |
| thymidine kinase 1 | *Tk1* | 99.38 | 1.74E-63 | Yes |
| fatty acid binding protein 2, intestinal | *Fabp2* | 99.37 | 1.76E-63 | |
| Galactokinase-1 | *Galk1* | 99.37 | 2.67E-63 | Yes |
| indolethylamine N-methyltransferase | *Inmt* | 99.37 | 2.67E-63 | |
| MACRO domain containing 1 | *Macrod1* | 99.36 | 3.23E-63 | Yes |
| BAI1-associated protein 2-like 1 | *Baiap2l1* | 99.36 | 3.23E-63 | |
| fibroblast growth factor 21 | *Fgf21* | 99.35 | 4.08E-63 | |
| gulonolactone (L-) oxidase | *Gulo* | 99.35 | 5.01E-63 | Yes |
| glutathione S-transferase, theta 3 | *Gstt3* | 99.35 | 6.23E-63 | Yes |

| | | | | |
|---|---|---|---|---|
| monocyte to macrophage differentiation-associated | *Mmd* | 99.34 | 7.75E-63 | |
| Aminolevulinate, delta-, synthase-2 | *Alas2* | 99.34 | 8.68E-63 | Yes |
| carboxylesterase 1D | *Ces1d* | 99.33 | 1.33E-62 | Yes |
| myosin IB | *Myo1b* | 99.33 | 1.33E-62 | |
| growth arrest and DNA-damage-inducible 45 beta | *Gadd45b* | 99.32 | 1.39E-62 | |
| low density lipoprotein receptor-related protein 4 | *Lrp4* | 99.32 | 1.39E-62 | |
| NAD(P)H dehydrogenase, quinone 1 | *Nqo1* | 99.32 | 1.39E-62 | |
| Lipoprotein lipase | *Lpl* | 99.32 | 1.44E-62 | Yes |
| macrophage expressed gene 1 | *Mpeg1* | 99.31 | 1.45E-62 | |
| ChaC, cation transport regulator 1 | *Chac1* | 99.31 | 1.84E-62 | |
| aldehyde dehydrogenase 1 family member A1 | *Aldh1a1* | 99.29 | 2.03E-62 | Yes |
| glutathione S-transferase, mu 1 | *Gstm1* | 99.29 | 2.03E-62 | Yes |
| interferon induced transmembrane protein 3 | *Ifitm3* | 99.29 | 2.03E-62 | |
| phosphatidylcholine transfer protein | *Pctp* | 99.29 | 2.03E-62 | |
| solute carrier family 29 (nucleoside transporters), m | *Slc29a1* | 99.29 | 2.03E-62 | |
| solute carrier family 6 (neurotransmitter transporter | *Slc6a12* | 99.29 | 2.03E-62 | |
| hydroxysteroid (17-beta) dehydrogenase 2 | *Hsd17b2* | 99.28 | 2.3E-62 | Yes |
| Alpha-aminoadipic semialdehyde synthase | *Aass* | 99.28 | 3.09E-62 | Yes |
| Phenylalanine hydroxylase | *Pah* | 99.28 | 3.09E-62 | Yes |
| RAB30, member RAS oncogene family | *Rab30* | 99.27 | 3.88E-62 | |
| zinc binding alcohol dehydrogenase, domain contain | *Zadh2* | 99.26 | 5.97E-62 | Yes |
| phenazine biosynthesis-like protein domain containi | *Pbld2* | 99.26 | 5.97E-62 | |
| cilia and flagella associated protein 20 | *Cfap20* | 99.26 | 6.61E-62 | |
| PDZ domain containing 1 | *Pdzk1* | 99.26 | 6.61E-62 | |
| Tyrosine aminotransferase, soluble | *Tat* | 99.25 | 7.14E-62 | Yes |
| lymphocyte antigen 6 complex, locus D | *Ly6d* | 99.25 | 9.18E-62 | |
| transglutaminase 1, K polypeptide | *Tgm1* | 99.24 | 1.08E-61 | Yes |
| RIKEN cDNA 4931408D14 gene | *4931408D14Rik* | 99.24 | 1.67E-61 | |
| keratin 23 | *Krt23* | 99.24 | 1.7E-61 | |
| Xanthine dehydrogenase (xanthine oxidase) | *Xdh* | 99.23 | 2.33E-61 | Yes |
| cysteine rich protein 2 | *Crip2* | 99.23 | 2.33E-61 | |
| glycerophosphodiester phosphodiesterase 1 | *Gde1* | 99.23 | 2.33E-61 | |
| RIKEN cDNA B230114P17 gene | *B230114P17Rik* | 99.22 | 2.75E-61 | |
| retinol dehydrogenase 9 | *Rdh9* | 99.22 | 2.98E-61 | Yes |

| | | | | |
|---|---|---|---|---|
| Acyl-Coenzyme A oxidase 1, palmitoyl | *Acox1* | 99.21 | 3.53E-61 | Yes |
| retinoic acid receptor responder (tazarotene inducec | *Rarres1* | 99.21 | 3.84E-61 | |
| glutathione peroxidase 7 | *Gpx7* | 99.2 | 4.67E-61 | Yes |
| insulin-like growth factor binding protein 2 | *Igfbp2* | 99.2 | 5.34E-61 | |
| metallothionein 1 | *Mt1* | 99.2 | 5.69E-61 | |
| carboxylesterase 1F | *Ces1f* | 99.19 | 7.01E-61 | Yes |
| 5-oxoprolinase (ATP-hydrolysing) | *Oplah* | 99.19 | 7.01E-61 | |
| cyclin-dependent kinase inhibitor 1A (P21) | *Cdkn1a* | 99.18 | 1.21E-60 | |
| lectin, galactose binding, soluble 4 | *Lgals4* | 99.18 | 1.32E-60 | |
| cytidine deaminase | *Cda* | 99.18 | 1.63E-60 | Yes |
| synaptosomal-associated protein, 47 | *Snap47* | 99.17 | 1.75E-60 | |
| Alkaline phosphatase, liver/bone/kidney | *Alpl* | 99.17 | 1.86E-60 | Yes |
| ATP binding cassette subfamily G member 2 (Junior b | *Abcg2* | 99.17 | 2.04E-60 | |
| high mobility group box 3 | *Hmgb3* | 99.16 | 3.49E-60 | |
| dehydrogenase/reductase (SDR family) member 4 | *Dhrs4* | 99.15 | 3.97E-60 | Yes |
| cytochrome P450, family 3, subfamily a, polypeptide | *Cyp3a13* | 99.15 | 3.97E-60 | |
| phospholipid scramblase 2 | *Plscr2* | 99.15 | 3.97E-60 | |
| carboxylesterase 2G | *Ces2g* | 99.15 | 5.7E-60 | Yes |
| complement component 2 (within H-2S) | *C2* | 99.14 | 5.97E-60 | |
| carboxylesterase 2E | *Ces2e* | 99.14 | 8.1E-60 | Yes |
| major facilitator superfamily domain containing 2A | *Mfsd2a* | 99.13 | 8.93E-60 | |
| inhibin beta-E | *Inhbe* | 99.13 | 9.08E-60 | |
| A kinase (PRKA) interacting protein 1 | *Akip1* | 99.13 | 1.73E-59 | |
| leukocyte cell-derived chemotaxin 2 | *Lect2* | 99.12 | 2.82E-59 | |
| Heme oxygenase 1 | *Hmox1* | 99.11 | 2.94E-59 | Yes |
| Aconitase, mitochondrial | *Aco2* | 99.11 | 2.94E-59 | Yes |
| solute carrier family 9 (sodium/hydrogen exchanger) | *Slc9a3r1* | 99.11 | 2.94E-59 | |
| SPARC related modular calcium binding 1 | *Smoc1* | 99.11 | 2.94E-59 | |
| translocase of inner mitochondrial membrane 8A1 | *Timm8a1* | 99.11 | 2.94E-59 | |
| doublecortin-like kinase 3 | *Dclk3* | 99.1 | 4.03E-59 | |
| Retinol dehydrogenase-5 | *Rdh5* | 99.09 | 4.4E-59 | Yes |
| etoposide induced 2.4 mRNA | *Ei24* | 99.09 | 4.4E-59 | |
| interferon regulatory factor 7 | *Irf7* | 99.09 | 4.4E-59 | |
| cytidine 5'-triphosphate synthase | *Ctps* | 99.08 | 7.85E-59 | |

| | | | | |
|---|---|---|---|---|
| Cystathionine gamma-lyase | *Cth* | 99.07 | 9.21E-59 | Yes |
| N-acetyltransferase 8 (GCN5-related) | *Nat8* | 99.07 | 9.21E-59 | |
| plasma membrane proteolipid | *Pllp* | 99.07 | 9.21E-59 | |
| serine (or cysteine) peptidase inhibitor, clade F, men | *Serpinf2* | 99.07 | 9.21E-59 | |
| solute carrier family 17 (sodium phosphate), membe | *Slc17a2* | 99.07 | 9.21E-59 | |
| B cell translocation gene 2, anti-proliferative | *Btg2* | 99.06 | 1.39E-58 | |
| Malonyl-CoA decarboxylase | *Mlycd* | 99.06 | 1.44E-58 | Yes |
| proteoglycan 4 (megakaryocyte stimulating factor, a | *Prg4* | 99.05 | 1.84E-58 | Yes |
| basic helix-loop-helix domain containing, class B9 | *Bhlhb9* | 99.05 | 1.98E-58 | |
| aldehyde dehydrogenase 16 family, member A1 | *Aldh16a1* | 99.04 | 2.81E-58 | |
| immediate early response 2 | *Ier2* | 99.04 | 2.81E-58 | |
| methyltransferase like 7B | *Mettl7b* | 99.04 | 2.81E-58 | |
| Carbamoyl-phosphate synthetase 1, mitochondrial | *Cps1* | 99.03 | 3.15E-58 | Yes |
| Ornithine aminotransferase | *Oat* | 99.03 | 3.15E-58 | Yes |
| Ketohexokinase (fructokinase) | *Khk* | 99.02 | 3.22E-58 | Yes |
| Carnitine palmitoyltransferase II | *Cpt2* | 99.02 | 3.22E-58 | Yes |
| Isocitrate dehydrogenase, soluble | *Idh1* | 99.02 | 3.22E-58 | Yes |
| 17-beta-hydroxysteroid dehydrogenase X | *Hsd17b10* | 99.02 | 3.22E-58 | Yes |
| carboxymethylenebutenolidase-like (Pseudomonas) | *Cmbl* | 99.02 | 3.22E-58 | |
| collagen, type XVIII, alpha 1 | *Col18a1* | 99.02 | 3.22E-58 | |
| poly (ADP-ribose) polymerase family, member 16 | *Parp16* | 99.02 | 3.22E-58 | |
| basic helix-loop-helix family, member e40 | *Bhlhe40* | 99 | 4.8E-58 | |
| nuclear receptor subfamily 1, group I, member 2 | *Nr1i2* | 99 | 4.8E-58 | |

| Metabolic disease associated with deficiency in human ortholog | Pathway in which enzyme regulates rate-limiting step |
|---|---|
| Hypercholanemia, familial, 607748 (3), Autosomal recessive | |
| Sjogren-Larsson syndrome, 270200 (3), Autosomal recessive | Oxidation of fatty aldehydes to fatty acids |
| Hyperprolinemia, type I, 239500 (3), Autosomal recessive; {Schizophrenia, s | Oxidation of proline to glutamate |
| Fanconi renotubular syndrome 3, 615605 (3), Autosomal dominant | |
| Homocystinuria, B6-responsive and nonresponsive types, 236200 (3), Autos | Trans-sulfuration pathway |
| | Regulates PDH, which catalyzes the rate limiting oxidation of pyruvate to AcC |
| [Histidinemia], 235800 (3), Autosomal recessive, Autosomal dominant | |
| Argininosuccinic aciduria, 207900 (3), Autosomal recessive | Arginine synthesis |
| | Conversion of PGE2 to PGF2alpha |

| | |
|---|---|
| HMG-CoA synthase-2 deficiency, 605911 (3), Autosomal recessive | Ketogenesis |
| Adenosine triphosphate, elevated, of erythrocytes, 102900 (3), Autosomal [illegible] | Glycolysis |
| {Hypercholesterolemia, familial, due to LDLR defect, modifier of}, 143890 (3), Autosomal dominant | |
| | Taurine biosynthesis |
| | Retinoic acid biosynthesis |
| Aspartate aminotransferase, serum level of, QTL1, 614419 (3) | |

2,4-dienoyl-CoA reductase deficiency (DECRD)

Unsaturated fatty acid oxidation in mitochondria

Linoleic acid metabolism

| | |
|---|---|
| Adrenal hyperplasia, congenital, due to 3-beta-hydroxysteroid dehydrogena | Aldosterone production |
| | Hexosamine biosynthesis |
| | All-trans retinoic acid biosynthesis |
| | 2-arachidonoglycerol biosynthesis |
| Sulfite oxidase deficiency, 272300 (3), Autosomal recessive | |
| [Sarcosinemia], 268900 (3), Autosomal recessive | |

| | |
|---|---|
| Dimethylglycine dehydrogenase deficiency, 605850 (3), Autosomal recessiv | Betaine metabolism |
| | Monoacylglycerol catabolism |
| Cerebral creatine deficiency syndrome 2, 612736 (3), Autosomal recessive | |
| Majeed syndrome, 609628 (3) | |
| {Panic disorder, susceptibility to}, 167870 (3), Autosomal dominant; {Schizophrenia, susceptibility to}, 181500 (3), Autosomal dominant | |
| Acyl-CoA dehydrogenase, medium chain, deficiency of, 201450 (3), Autosor | Medium-chain fatty acid β-oxidation |
| | Unsaturated fatty acid oxidation in mitochondria |
| | Nucleotide salvage pathway |
| Galactokinase deficiency with cataracts, 230200 (3), Autosomal recessive | |
| | Ascorbic acid biosynthesis |

| | |
|---|---|
| Anemia, sideroblastic, 1, 300751 (3), X-linked recessive; Protoporphyria, ery | Heme biosynthesis |
| Combined hyperlipidemia, familial, 144250 (3), Autosomal dominant; [High | Hydrolysis of core TGs from TG-rich lipoproteins |
| | Retinoic acid biosynthesis |
| Hyperlysinemia, 238700 (3), Autosomal recessive; Saccharopinuria, 268700 | Lysine catabolism |
| [Hyperphenylalaninemia, non-PKU mild], 261600 (3), Autosomal recessive; | Phenylalanine catabolism |
| Tyrosinemia, type II, 276600 (3), Autosomal recessive | Tyrosine catabolism |
| Autosomal recessive congenital ichthyosis | Formation of cross-linked protein envelope in terminal differentiation of skin |
| Xanthinuria, type I, 278300 (3), Autosomal recessive | Purine metabolism |
| | All-trans retinoic acid biosynthesis |

Peroxisomal acyl-CoA oxidase deficiency, 264470 (3), Autosomal recessive Peroxisomal fatty acid β-oxidation pathway

Hypophosphatasia, adult, 146300 (3), Autosomal recessive, Autosomal dominant; Hypophosphatasia, childhood, 241510 (3), Autosomal recessive; Hypop

Heme oxygenase-1 deficiency, 614034 (3); {Pulmonary disease, chronic obs Heme degradation

Infantile cerebellar-retinal degeneration, 614559 (3), Autosomal recessive; ?Optic atrophy 9, 616289 (3), Autosomal recessive

Fundus albipunctatus, 136880 (3), Autosomal recessive, Autosomal domina All-trans retinoic acid synthesis

| | |
|---|---|
| Cystathioninuria, 219500 (3), Autosomal recessive; Homocysteine, total pla | Cysteine synthesis |
| Malonyl-CoA decarboxylase deficiency, 248360 (3), Autosomal recessive | Conversion of malonyl-CoA to acetyl-CoA and $CO_2$ |
| Carbamoylphosphate synthetase I deficiency, 237300 (3), Autosomal recess | Urea cycle |
| Gyrate atrophy of choroid and retina with or without ornithinemia, 258870 (3), Autosomal recessive | |
| [Fructosuria], 229800 (3), Autosomal recessive | Conversion of fructose to fructose-1-phosphate |
| CPT II deficiency, infantile, 600649 (3), Autosomal recessive; CPT II deficienc | Mitochondrial fatty acid beta-oxidation |
| {Glioma, susceptibility to, somatic}, 137800 (3) | TCA cycle |
| HSD10 mitochondrial disease, 300438 (3), X-linked dominant | |

CoA

phosphatasia, infantile, 241500 (3), Autosomal recessive; Odontohypophosphatasia, 146300 (3), Autosomal recessive, Autosomal dominant
