## Supplementary File 7 for "The Signaling Pathways Project: an integrated ‘omics knowledgebase for mammalian cellular signaling pathways"

**Supplementary File 7. Genes encoding metabolic enzymes in the 99th percentile of the All nodes-Mm-liver transcriptomic consensome whose deficiency is associated with a human metabolic disorder.** Gene symbol links point to SPP transcriptomic Regulation Reports filtered for mouse liver. Disease links point to OMIM entries highlighted for the corresponding human gene.

| Target | Gene product | CPV | Hepatic metabolic pathway | Known human deficiency disease |
|---|---|---|---|---|
| **Lipid metabolism** | | | | |
| *Ephx1* | Epoxide hydroxylase 1, microsomal xenobiotic | 7.27E-107 | Conversion of epoxides to trans-dihydrodiols for conjugation and excretion | Familial hypercholanemia |
| *Aldh3a2* | Aldehyde dehydrogenase 3 family, member A2 (fatty aldehyde dehydrogenase) | 2.11E-103 | Rate-limiting oxidation of fatty aldehydes to fatty acids | Sjogren-Larsson syndrome |
| *Ehhadh* | Enoyl-Coenzyme A, hydratase/3-hydroxyacyl Coenzyme A dehydrogenase | 4.70E-89 | Essential for the production of medium-chain dicarboxylic acids | Fanconi renotubular syndrome 3 |
| *Hmgcs2* | 3-hydroxy-3-methylglutaryl-coenzyme A synthase 2, mitochondrial | 1.23E-80 | Rate-limiting enzyme of ketogenesis | HMG-CoA synthase-2 deficiency |
| *Ephx2* | Epoxide hydrolase 2, cytoplasmic | 6.06E-78 | Conversion of epoxides to trans-dihydrodiols for conjugation and excretion | Familial hypercholesterolemia |
| *Decr1* | 2,4-dienoyl-CoA reductase 1 | 6.23E-71 | Rate-limiting step of unsaturated fatty acid oxidation in mitochondria | DECR deficiency |
| *Acadm* | Acyl-Coenzyme A dehydrogenase, C-4 to C-12 straight chain | 9.52E-65 | Rate limiting step of medium-chain fatty acid β-oxidation | ACADM deficiency |
| *Lpin2* | Lipin 2 | 1.22E-65 | Conversion of phosphatidic acid to diacylglycerol during triglyceride biosynthesis | Majeed syndrome |
| *Lpl* | Lipoprotein lipase | 1.44E-62 | Rate-limiting hydrolysis of core TGs from TG-rich lipoproteins | Familial combined hyperlipidemia |
| *Acox1* | Acyl-Coenzyme A oxidase 1, palmitoyl | 3.53E-61 | Rate-limiting enzyme in the peroxisomal fatty acid β-oxidation pathway | Peroxisomal acyl-CoA oxidase deficiency |
| *Cpt2* | Carnitine palmitoyltransferase II | 3.22E-58 | Rate limiting step in mitochondrial fatty acid beta-oxidation | Neonatal CPT II deficiency |
| *Mlycd* | Malonyl-CoA decarboxylase | 1.44E-58 | Conversion of malonyl-CoA to acetyl-CoA and $CO_2$ | MLYCD deficiency |
| **Carbohydrate metabolism** | | | | |

| | | | | |
|---|---|---|---|---|
| *Pklr* | pyruvate kinase L/R | 1.04E-79 | Rate-limiting step in glycolysis | Pyruvate kinase deficiency |
| *Got1* | Aspartate aminotransferase, cytoplasmic | 4.30E-76 | Transamination of aspartate & oxaloacetate in gluconeogenesis | Aspartate aminotransferase, serum level of, QTL1 |
| *Galk1* | Galactokinase-1 | 2.66E-63 | Phosphorylation of α-D-galactose to galactose 1-phosphate in the Leloir pathway | Galactokinase deficiency with cataracts |
| *Aco2* | Aconitase, mitochondrial | 2.93E-59 | Stereo-specific isomerization of citrate to isocitrate | Infantile cerebellar-retinal degeneration |
| *Idh1* | Isocitrate dehydrogenase, soluble | 3.21E-58 | Rate-limiting step of the TCA cycle | Glass syndrome, Ollier disease |
| *Khk* | Ketohexokinase (fructokinase) | 3.21E-58 | Rate-limiting conversion of fructose to fructose-1-phosphate | Fructosuria |
| **Amino acid metabolism** | | | | |
| *Prodh* | Proline dehydrogenase (proline oxidase) | 2.54E-94 | Rate-limiting oxidation of proline to glutamate | Hyperprolinemia |
| *Hal* | Histidine ammonia-lyase | 3.55E-86 | Initial reaction in histidine catabolism | Histidinemia |
| *Asl* | Argininosuccinate lyase | 1.54E-83 | Rate-limiting step in arginine synthesis | Argininosuccinic aciduria |
| *Sardh* | Sarcosine dehydrogenase | 6.41E-67 | N-demethylation of sarcosine to glycine | Sarcosinemia |
| *Dmgdh* | Dimethylglycine dehydrogenase | 1.22E-65 | Rate-limiting step in betaine metabolism | DMGDH deficiency |
| *Aass* | Alpha-aminoadipic semialdehyde synthase | 3.09E-62 | Rate-limiting step in lysine catabolism | Hyperlysinemia |
| *Pah* | Phenylalanine hydroxylase | 3.09E-62 | Rate-limiting step in phenylalanine catabolism | Phenylketonuria |
| *Tat* | Tyrosine aminotransferase, soluble | 7.14E-62 | Rate-limiting step in tyrosine catabolism | Type II tyrosinemia |
| *Cth* | Cystathionine gamma-lyase | 9.20E-59 | Rate-limiting step in cysteine synthesis | Cystathioninuria |
| *Oat* | Ornithine aminotransferase | 3.14E-58 | Glutamate biosynthesis | Gyrate atrophy |
| **Other metabolic pathways** | | | | |
| *Cbs* | Cystathionine beta-synthase | 6.47E-87 | Rate-limiting and initial enzyme in the trans-sulfuration pathway. | Homocysteinemia |
| *Hsd3b2* | 3beta-hydroxysteroid dehydrogenase/delta(5)-delta(4)isomerase type II | 1.01E-68 | Rate limiting in aldosterone production | Congenital adrenal hyperplasia |
| *Suox* | Sulfite oxidase | 2.12E-67 | Oxidation of sulfite to sulfate | Sulfite oxidase |

| | | | | |
|---|---|---|---|---|
| | | | | deficiency |
| *Gamt* | Guanidinoacetate N-methyltransferase | 1.22E-65 | Creatine biosynthesis | Cerebral creatine deficiency |
| *Comt* | Catechol-O-methyltransferase | 5.26E-65 | Degradation of catecholamines | Panic disorder, schizophrenia |
| *Alas2* | Aminolevulinate, delta-, synthase-2 | 8.68E-63 | Rate-limiting in heme biosynthesis | Sideroblastic anemia |
| *Alpl* | Alkaline phosphatase, liver/bone/kidney | 1.86E-60 | General hydrolysis of phosphate esters | Adult hypophosphatasia |
| *Tgm1* | Transglutaminase 1 | 1.08E-61 | Rate limiting in formation of cross-linked protein envelope during terminal skin differentiation | Autosomal recessive congenital ichthyosis |
| *Xdh* | Xanthine dehydrogenase (xanthine oxidase) | 2.33E-61 | Rate-limiting enzyme in purine metabolism | Type I xanthinuria |
| *Hmox1* | Heme oxygenase 1 | 2.93E-59 | Rate-limiting enzyme of heme degradation | HMOX1 deficiency |
| *Cps1* | Carbamoyl-phosphate synthetase 1, mitochondrial | 3.14E-58 | First rate-limiting mitochondrial enzyme in the urea cycle | CPS1 deficiency |
| *Hsd17b10* | 17-beta-hydroxysteroid dehydrogenase X | 3.21E-58 | Beta-oxidation at position 17 of androgens and estrogens | HSD10 mitochondrial disease |
| *Rdh5* | Retinol dehydrogenase-5 | 4.39E-59 | Rate limiting step in all-trans retinoic acid synthesis | Fundus albipunctatus |
